# A novel expression domain of *extradenticle* underlies the evolutionary developmental origin of the chelicerate patella

**DOI:** 10.1101/2024.05.16.594547

**Authors:** Benjamin C. Klementz, Georg Brenneis, Isaac A. Hinne, Ethan M. Laumer, Sophie M. Neu, Grace M. Hareid, Guilherme Gainett, Emily V.W. Setton, Catalina Simian, David E. Vrech, Isabella Joyce, Austen A. Barnett, Nipam H. Patel, Mark S. Harvey, Alfredo V. Peretti, Monika Gulia-Nuss, Prashant P. Sharma

## Abstract

Neofunctionalization of duplicated gene copies is thought to be an important process underlying the origin of evolutionary novelty and provides an elegant mechanism for the origin of new phenotypic traits. One putative case where a new gene copy has been linked to a novel morphological trait is the origin of the arachnid patella, a taxonomically restricted leg segment. In spiders, the origin of this segment has been linked to the origin of the paralog *dachshund-2*, suggesting that a new gene facilitated the expression of a new trait. However, various arachnid groups that possess patellae do not have a copy of *dachshund-2*, disfavoring the direct link between gene origin and trait origin. We investigated the developmental genetic basis for patellar patterning in the harvestman *Phalangium opilio*, which lacks *dachshund-2*. Here, we show that the harvestman patella is established by a novel expression domain of the transcription factor *extradenticle*. Leveraging this definition of patellar identity, we surveyed targeted groups across chelicerate phylogeny to assess when this trait evolved. We show that a patellar homolog is present in Pycnogonida (sea spiders) and various arachnid orders, suggesting a single origin of the patella in the ancestor of Chelicerata. A potential loss of the patella is observed in Ixodida. Our results suggest that the modification of an ancient gene, rather than the neofunctionalization of a new gene copy, underlies the origin of the patella. Broadly, this work underscores the value of comparative data and broad taxonomic sampling when testing hypotheses in evolutionary developmental biology.

## Introduction

One of the central goals of evolutionary developmental biology remains demonstrating the mechanistic basis by which evolutionary novelty arises, including both morphological and molecular innovation through modifications of development (reviewed in Hall [2012]). Two competing hypotheses pertain to the genetic basis for the origin of novelty. The first is the acquisition of a novel function by an existing gene. This can be accomplished via diverse mechanisms, including heterochrony (de Beer 1930; Raff 1992; MacDonald and Hall 2001; Smith 2003; Tills et al. 2011), the modification of gene expression domains (Carpio et al. 2004; Janssen et al. 2021), alterations in *cis* regulatory interactions (Mazo-Vargas et al. 2022; Tendolkar et al. 2024), and the cooption of existing gene regulatory networks (Moczek and Rose 2009; Setton and Sharma 2018). The second explanation ties evolutionary novelty with the birth of new genes, with new functions either generated *de novo* (Cai et al. 2008; Knowles and McLysaght 2009) or via divergence following tandem or whole genome duplication events (Haldane 1990; Dittmar and Liberles 2011). While pseudogenization and eventual loss are understood to be the most likely long-term outcome for duplicated genes (Ohno 1970; Brunet et al. 2006; Gout et al. 2023), copies that persist may either subdivide the ancestral gene’s function (subfunctionalization) and thereby reduce pleiotropy, or one daughter copy may acquire a novel function (neofunctionalization) (Lynch and Conery 2000). These processes are non-mutually exclusive and may act in tandem (He and Zhang 2005). However, neofunctionalization of new gene copies is an especially compelling phenomenon when it is linked to novel traits.

One prominent case of neofunctionalization appears to underly the origin of the patella, a segment (or podomere) found in the pedipalps and walking legs of a subset of chelicerate arthropods (sea spiders, horseshoe crabs, and terrestrial arachnids) (Turetzek et al. 2016) (fig. 1*a*). Much of the evolutionary success of Arthropoda, the most species-rich of the metazoan phyla, can be attributed to their eponymous jointed appendages. Subdivision of these appendages into discrete, sclerotized podomeres not only improves appendage flexibility and range of motion but also provides additional substrates for morphological innovation and regionalization (Brusca and Brusca 2003). This regionalization has given rise to a wide range of appendage forms and functions, including modification of the first pair of walking legs into the venom-injecting forcipules of centipedes, the raptorial appendages of mantis shrimps, the grasping pedipalps of scorpions, or the antenniform sensory legs of whip spiders and vinegaroons, ultimately enabling the phylum to exploit all but the most extreme habitats. The presence of the patella, however, differentiates the six-segmented leg of a typical mandibulate arthropod (myriapods, crustaceans, and insects) from the seven-segmented walking legs of arachnids like spiders and scorpions. Chelicerate pedipalps also possess a patella but are six-segmented due to absence of the metatarsal segment.

**Figure 1.**
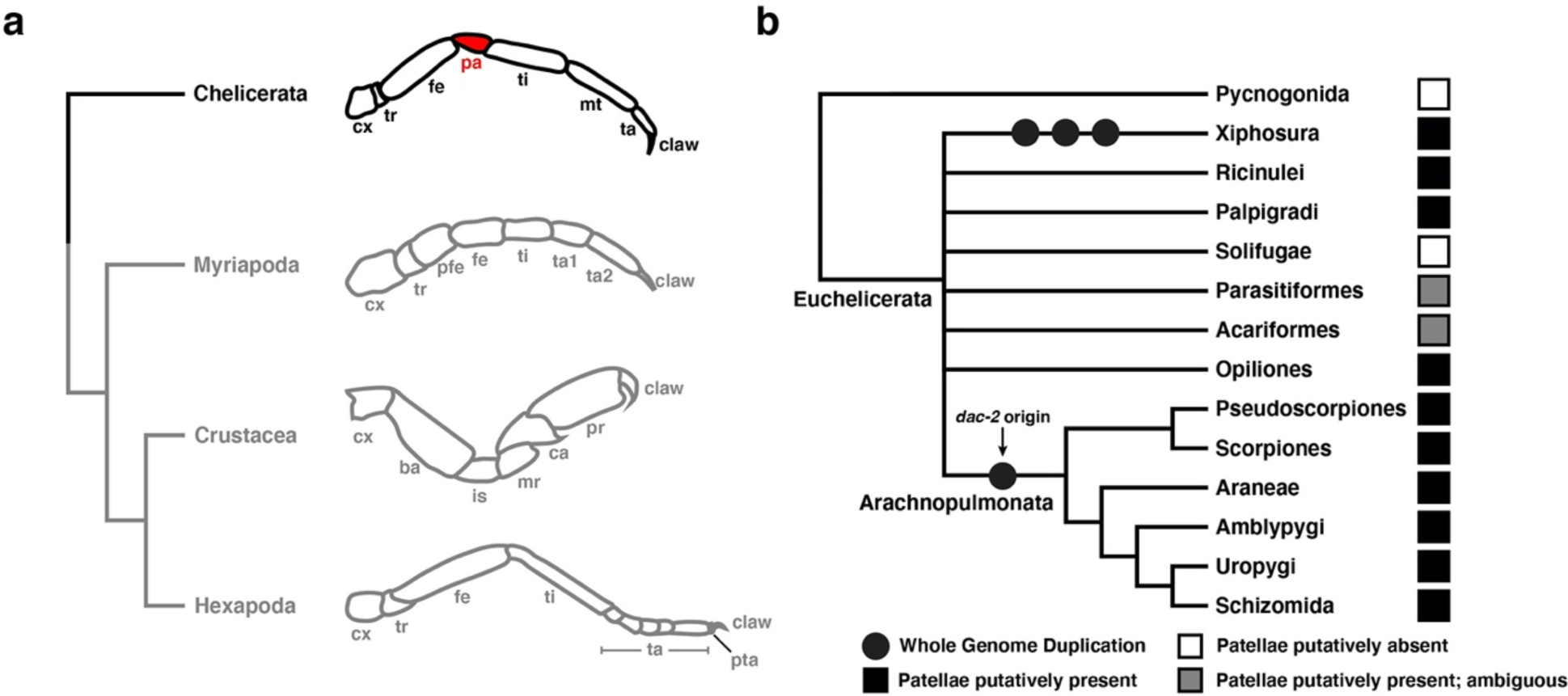
The patella differentiates the appendages of most chelicerate orders from other arthropods. (**a**) Exemplars of leg architecture across major arthropod lineages. Appendage schematics from top to bottom: spider leg; centipede leg; amphipod cheliped; insect leg. (**b**) Simplified phylogeny of Chelicerata. Icons indicate condition of patella. Origin of *dac-2* has been inferred to originate from whole-genome duplication in the arachnopulmonate ancestor. Abbreviations: cx, coxa; tr, trochanter; fe, femur; pa, patella; ti, tibia; mt, metatarsus; ta, tarsus; pfe, prefemur; ba, basis; is, ischium; mr, merus; ca, carpus; pr, propodus; pta, pretarsus. Chelicerate tree topology based on (Ballesteros et al. 2022) with unstable nodes collapsed.

The evolutionary origin of the patella was recently inferred to result from the neofunctionalization of a duplicated paralog of the conserved appendage patterning gene *dachshund* (*dac*) (Turetzek et al. 2016). During establishment of the arthropod proximo-distal (PD) limb axis, *dac* is one of four primary transcription factors (alongside *homothorax* (*hth*), *extradenticle* (*exd*), and *Distal-less* (*Dll*)), that regionalizes the PD axis; in the fruit fly *Drosophila melanogaster,* the canonical *dac* loss-of-function phenotype exhibits a deletion of medial segments (Dong et al. 2001). Bioinformatic and gene expression surveys have also inferred that a single copy of *dac* was present in the common ancestor of Panarthropoda (Tardigrada, Onychophora, and Arthropoda) (Prpic and Tautz 2003; Angelini and Kaufman 2005; Janssen et al. 2010; Sharma et al. 2012a; Barnett and Thomas 2013). Previous functional studies in pancrustaceans and one arachnid species demonstrated a conserved role for *dac* in patterning the medial territory of the walking leg across Arthropoda (Angelini and Kaufman 2004; Sewell et al. 2008; Angelini et al. 2009; Angelini et al. 2012; Sharma et al. 2012a; Sharma et al. 2013; Sugime et al. 2019). Intriguingly, two copies of *dac* occur in some arachnids like spiders, and it was previously shown that the two copies exhibit dissimilar expression domains during embryogenesis of two spider species (Turetzek et al. 2016). One copy, *dac-1,* retains the conserved medial expression domain characteristic of the *dac* single copy ortholog in other arthropods, whereas the second, *dac-2,* exhibits a proximal domain in the coxa and body wall, as well as a novel distal ring of expression in the presumptive patellar segment (Turetzek et al. 2016). Maternal RNA interference (RNAi) against *dac-2* in the spider *Parasteatoda tepidariorum* resulted in a loss of the patellar-tibial boundary and consequent fusion of these segments. This result was interpreted to mean that the origin of the patella was caused by neofunctionalization of the *dac-2* copy, as *dac-2* was shown to be present at least in the common ancestor of spiders and scorpions (and by extension, across chelicerates). RNAi against *dac-1* was not reported in that work, and no functional data are available for *dac-1* in any spider species. As such, the inference of an ancestral function for *dac-1* (PD axis patterning in the medial segments) is based solely upon gene expression patterns in the two spider models (Turetzek et al. 2016).

On their own, these results certainly implicate *dac-2* as responsible for the origin of the patella. But, this reconstruction is problematic for several, interconnected reasons, especially given recent advances in chelicerate genomics. First, *dac-2* is not common to all arachnids, but rather, restricted only to a subdivision of chelicerate orders called Arachnopulmonata, which are united by a shared whole-genome duplication (which is thought to have given rise to *dac-2*; Nolan et al. 2020) (fig. 1*b*). Many lineages of apulmonate arachnids (e.g., Opiliones, Ricinulei) that diverged prior to this genome duplication event have only a single copy of *dac*, but putatively possess a patella. Thus, *dac-2* is significantly younger than the origin of this novel podomere. In typical cases of novelty arising from neofunctionalization, a novel trait occurs either phylogenetically coincident with, or subsequent to, the appearance of the new gene that facilitates its expression. Notably, several copies of *dac* also occur in Xiphosura, but these are unrelated to the arachnopulmonate duplication as Xiphosura has experienced three rounds of comparatively recent and lineage-specific whole genome duplication (resulting in nine copies of *dac*; Nong et al. 2021). Thus, though Xiphosura also possess patellae, they do not possess an ortholog of spider *dac-2.* These patterns suggest that the origin of the patella is older than, and therefore not a consequence of, *dac-2* origin.

Second, a clear definition of the patella is elusive, and its presence is controversial in various chelicerate orders (fig. 2; supplementary tables S1, S2). Traditionally, the patella has been contextually defined as the small, compact podomere falling between the larger femur and tibia; it is therefore the fourth podomere of the chelicerate walking leg. Scorpions, however, possess an elongate patella that exceeds the size of the more distal tibia and creates the prominent rectangular bend characteristic of their walking legs (fig. 2*h*). The fourth podomere of pseudoscorpion appendages is likewise elongated (fig. 2*g*). Historically, the putative homology of this podomere was subject of much debate with several prominent works refuting presence of the patella throughout the order, instead opting to ascribe a bipartite femur (e.g., Savory 1964), while others argued for its presence as a synapomorphy of Arachnida (e.g., Snodgrass 1958). Yet, convergent fusions across pseudoscorpion families, between metatarsus and tarsus, as well as between both segments of the bipartite femur, confounded consistent alignment of their podomeres with other arachnid orders. More recent works, however, have routinely defined the fourth podomere of pseudoscorpions as the patella, irrespective of podomere size (Shultz 1989; Harvey 1992; Michalski et al. 2022).

**Figure 2.**
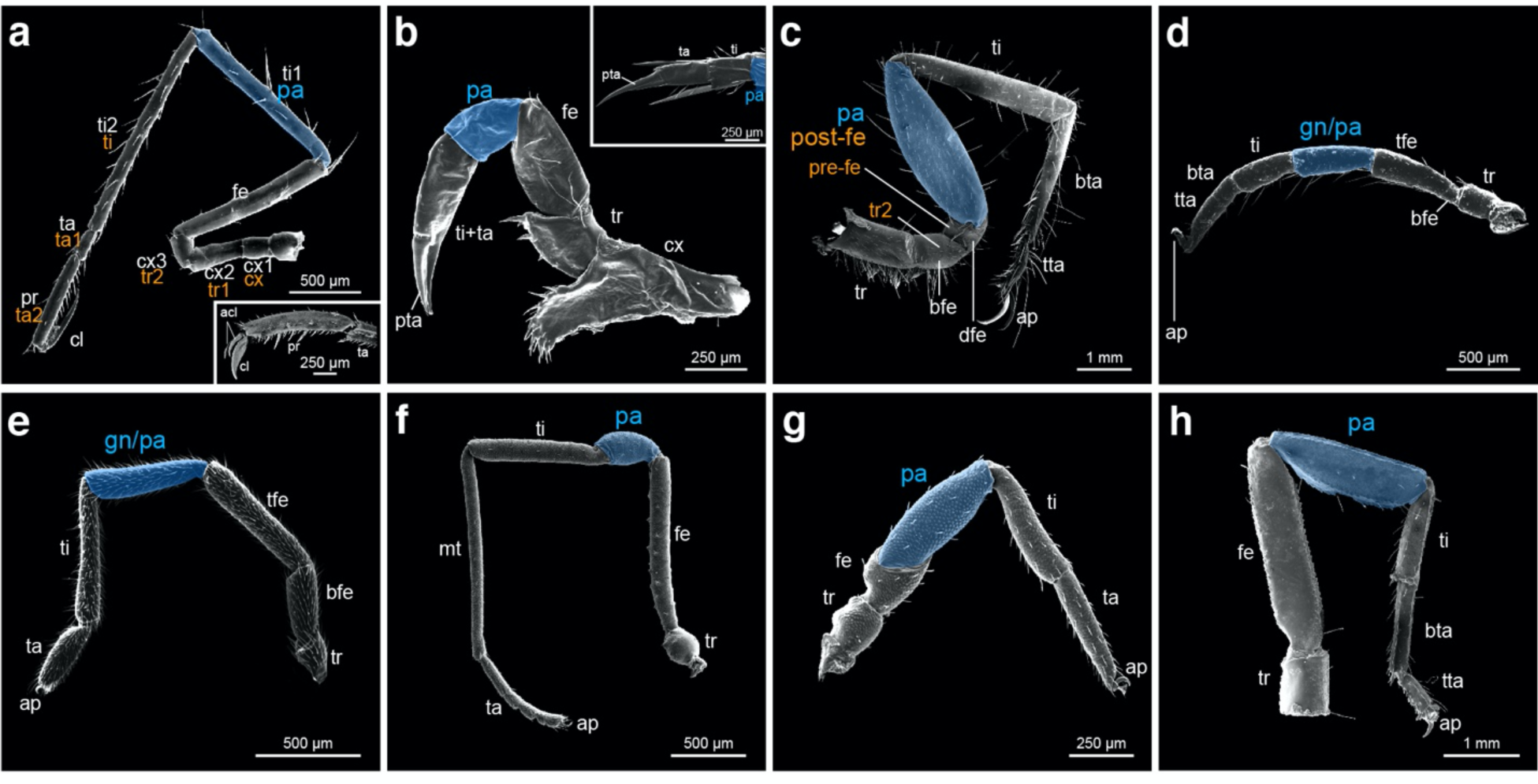
The patella is inconsistently defined across Chelicerata. (**a-h**) Scanning electron micrographs of legs for selected orders. (**a**) Leg four of sea spider *Nymphon* sp. (Pycnogonida). Inset: distal podomere morphology of *Nymphon gracile* leg. (**b**) Leg four of horseshoe crab *Limulus polyphemus* (Xiphosura). Inset: distal podomere morphology of the pusher leg (leg V). (**c**) Leg four of camel spider *Mummucia sp.* (Solifugae). (**d**) Leg four of tick *Ixodes scapularis* (Parasitiformes, Ixodida). (**e**) Leg one of velvet mite *Trombidium* sp. (Acariformes, Trombidiformes). (**f**) Leg three of harvestman *Zalmoxis furcifer* (Opiliones). (**g**) Leg one of pseudoscorpion *Pselaphochernes parvus* (Pseudoscorpiones). (**h**) Leg one of scorpion *Scorpio palmatus* (Scorpiones). Blue shading: putative patellar homologs. White and orange text indicate alternative nomenclature. Abbreviations: acl: auxiliary claw; cx, coxa; tr, trochanter; fe, femur; bfe, basifemur; tfe, telofemur; pa, patella; gn, genu; ti, tibia; mt, metatarsus; ta, tarsus; bta, basitarsus; tta, telotarsus; pr, propodus; cl, claw; pta, pretarsus; ap, apotele.

Exacerbating this confusion, the phylogenetic placement of Pseudoscorpiones had long been unstable. Pseudoscorpions were initially thought to be closely related to Solifugae (camel spiders), which similarly lack an obvious patellar segment but clearly possess a metatarsus. The putative shared absence of a patella had thus been historically leveraged as one of several morphological characters uniting Pseudoscorpiones and Solifugae in the clade Apatellata (Haplocnemata) (van der Hammen 1977; van der Hammen 1985; van der Hammen 1986), though this relationship has never been recovered by analyses of molecular sequence data (e.g., Regier et al. 2010; Sharma et al. 2014; Ballesteros and Sharma 2019; Ballesteros et al. 2022). Likewise, Solifugae have historically demonstrated an unstable phylogenetic position across molecular analyses (Regier et al. 2010; Sharma et al. 2014; Ballesteros et al. 2019; Ballesteros and Sharma 2019; Ballesteros et al. 2022). However, recent sequencing efforts to produce the first embryonic transcriptomes of both Pseudoscorpiones and Solifugae refuted the Apatellata hypothesis on the basis of a shared whole genome duplication between pseudoscorpions and the remaining arachnopulmonates, to the exclusion of Solifugae (Ontano et al. 2021; Gainett et al. 2023) (fig 1*b*).

At present, although a sister group relationship between the two orders has been overturned, debate persists over the homology of their podomeres. An additional podomere is found in the third and fourth walking legs of solifuges, as well as in all walking legs of groups like Ricinulei (hooded tick spiders), Trigonotarbida (extinct), and some Eurypterida (extinct) (Shultz 1989), but many workers consider this segment to be homologous to the more distal segment of a bipartite femur (i.e., basifemur and distifemur; Punzo 1998). Many other chelicerate orders exhibit discordant leg segment alignments, each with their own lineage-specific nuances. The segmental homologies of Acariformes (mites) and Parasitiformes (mites and ticks; together forming the likely polyphyletic Acari) are likewise obscured. As in pseudoscorpions, many groups of acariform mites possess a subdivision of the femur forming basi- and telofemur, which therefore represents the fourth podomere of their walking legs. The adjacent segment, rather than patella, is instead referred to as the genu. However, this terminology was largely adopted subsequent to the early work of A.D. Michael on British oribatid mites which themselves possess a secondary fusion of both femora in all but the most early-diverging families (Michael 1884), synthesizing alternative terms such as *la jambe* (Dugès 1834), femur (Fumouze and Robin 1867), or 1^st^ article (Donnadieu 1875), and use of genu by earlier authors (Nicolet 1855). The alignment of genu and patella in the fourth podomere of Oribatida and many other arachnid orders had thus been supposed to reflect homology and select authors have since used the terms interchangeably in both taxonomic and comparative arachnological works (e.g., Shultz 1989; Evans 1992; Harvey 1998; Krantz et al. 2009). Similarly, parasitiform mites and ticks also possess a genu segment and largely retain ancestral basi- and telofemora (Krantz et al. 2009). Opilioacarida, the putative sister group to the remaining Parasitiformes, present additional subdivision of the trochanter (basi-, telotrochanter), although patterns of segment innervations suggest the subdivision is superficial (van der Hammen 1970).

These patterns of segment gain and loss across chelicerate orders undermine definitions for the patella that are based upon either position or shape. Inferring the origin of the patella is further stymied by leg architecture in Pycnogonida, the sister group of the remaining chelicerates. In sea spiders, numerous historical and primarily descriptive works have failed to converge on a consistent terminology for the pycnogonid podomeres, complicating interpretations of segmental homology. All studies on extant species report the presence of eight podomeres (when not including the distal claw), but most sea spider workers infer the leg to consist of three coxae, a femur, two tibiae, and two distal segments (Hoek 1881; Sars 1891; Meinert 1899; Helfer and Schlottke 1935; King 1973; Arnaud and Bamber 1988; Brusca and Brusca 2003). Only a minority of works have inferred the fifth podomere to be the putative patellar segment (Snodgrass 1958; Dencker 1974; Schram and Hedgpeth 1978; Shultz 1989; Sabroux et al. 2023).

Thus, both the developmental genetic basis for the origin of this podomere, as well as its incidence across Chelicerata, is poorly understood. Breaking this impasse therefore requires the identification of a potential genetic mechanism underlying patella formation that is phylogenetically consistent with the age of the trait, and that can also be surveyed across chelicerate taxa to test for the presence of a patella. Of the four transcription factors required for the establishment of the arthropod PD axis (Dong et al. 2001), the arachnid homolog of *extradenticle* (*exd*) exhibits both ancestral and novel expression domains. Shared with other arthropods, arachnid *exd* has a conserved proximal domain (overlapping with *homothorax* expression; the two genes operate as a heterodimer to pattern proximal leg segments). Additionally, arachnid *exd* has a novel distal ring of expression in the presumptive patella. This distal expression domain is observed in both the single copy ortholog of *exd* in the harvestman *Phalangium opilio* (Sharma et al. 2012a), as well as *exd* paralogs of spiders and scorpions (Pechmann and Prpic 2009; Nolan et al. 2020), but its function is not known.

Here, we show that this distal ring domain of *exd* is required for patterning the patella-tibia boundary of the harvestman *Phalangium opilio.* Disruption of Notch signaling results in diminution of this distal *exd* domain, supporting the interpretation that it plays a role in segmentation. Armed with a developmental genetic definition of patellar identity, we survey exemplars of chelicerate diversity and demonstrate that a patella homolog is present in phylogenetically significant chelicerate groups, such as sea spiders, mites, and pseudoscorpions. These results suggest that a patella was present in the common ancestor of Chelicerata.

## Results

### A distal domain of exd localizes to the patella-tibia boundary during P. opilio embryonic appendage development

In early outgrowth of the limb bud (stage 9; stages following Gainett et al. [2022]), *Po-dac* is expressed in a medial territory proximal to the expression domain of *Po-Distal-less* (*Po-Dll*) (fig. 3*a*; supplementary fig. S1-S3). At this early stage, *Po-exd* is detected as two discontinuous domains: one occurs proximally in the presumptive coxal segment and body wall, whereas a second ectodermal ring is observed distal to the *Po-dac* domain. Weak, diffuse *Po-exd* expression is also observed distal to the ring domain, into the tip of the appendage during this early stage. As the appendages elongate in later stages of development, the medial *Po-dac* domain and the distal *Po-exd* domain expand and become heterogeneous in expression intensity along the PD axis (fig. 3*b*-3*d*; supplementary figure S3). By stage 10, the *Po-dac* domains exhibit two stronger rings of expression that correspond to the presumptive distal trochanter and the distal femur, with weaker expression signal between these domains (fig. 3*b*; supplementary figure S3). Subsequent refinement of these domains in stage 11 and stage 12 embryos yields stronger localization to the distal femur and trochanter territories and weaker intermediate expression (fig. 3*c*, 3*d*; supplementary figure S3). The restricted expression of *Po-dac* in the medial territory of the developing appendage mirrors *dac* expression domains across other arthropod lineages, particularly insects, wherein *dac* loss-of-function mutants exhibit deletion of the femur and tibia (Angelini and Kaufman 2004; Sewell et al. 2008; Angelini et al. 2009; Angelini et al. 2012).

**Figure 3.**
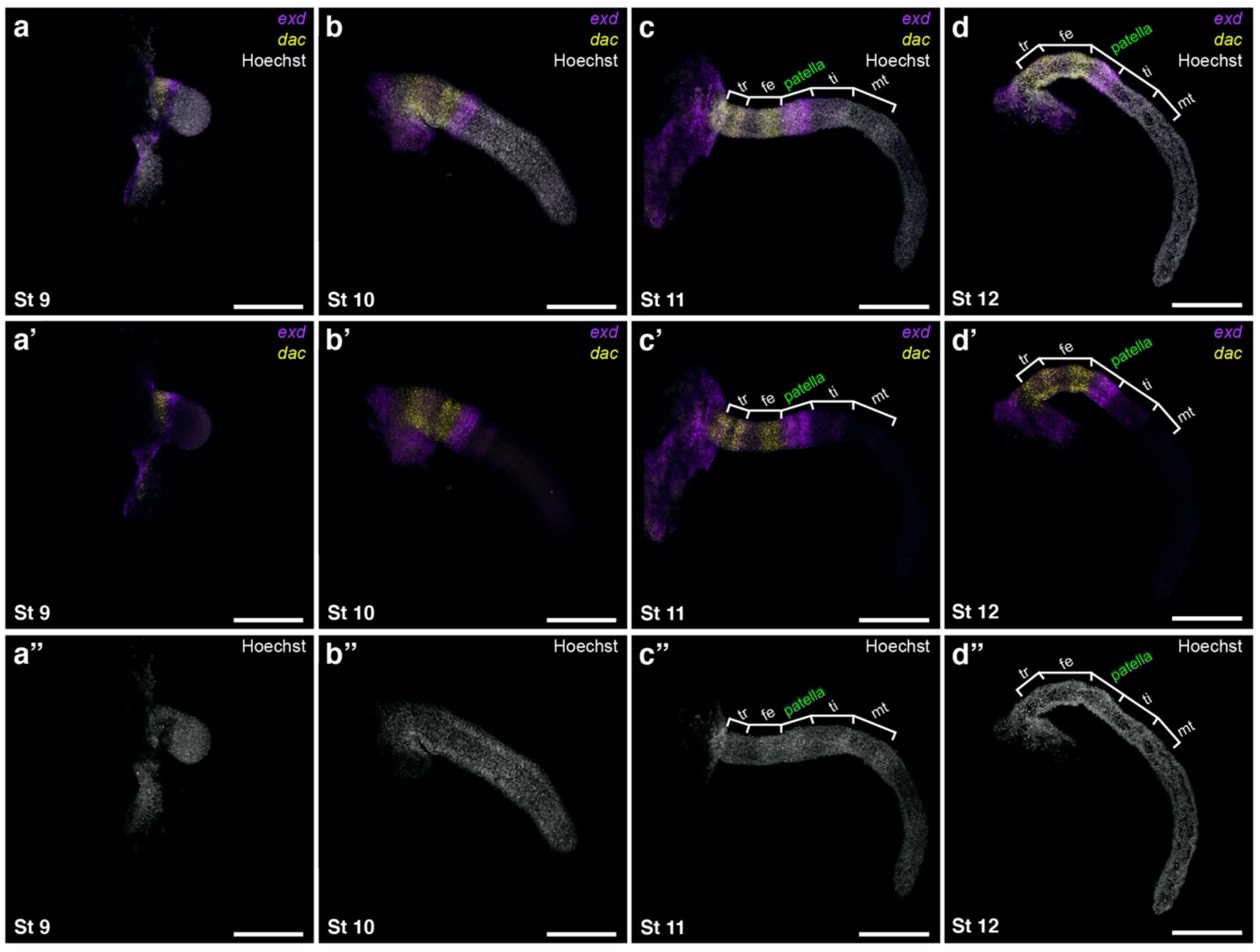
A distal ring domain of *Po-exd* is established early in *P. opilio* embryonic development and localizes to the patella-tibia segmental boundary. (**a-d**) Leg mounts of L2 in selected stages with merged visualization of Hoechst counterstaining (white), *Po-exd* (magenta), and *Po-dac* (yellow). (**a’**-**d’**) Multiplexed expression of *Po-exd* and *Po-dac*. (**a’’**-**d’’**) Isolated Hoechst counterstaining. Scale bars: 250 µm.

In contrast to mandibulate development (e.g., insect, amphipod, or millipede), a distal ring of *Po-exd* expression is detected early in the outgrowth of the limb bud (stage 9), with minimal overlap of expression with the more proximal *dac* domain (fig. 3*a*; supplementary figure S3). By stage 11, this domain encompasses nearly the entire patellar segment, whereas the weaker, more distal expression localizes to the presumptive tibial segment (fig. 3*b*, 3*c*; supplementary figure S3). In stage 12 embryos, the strong *Po-exd* domain concentrates at the patella-tibia boundary, whereas the more proximal patellar territory experiences a slight gradation of weakening expression and slight expansion of the *Po-dac* territory into the femur-patella boundary (fig. 3*d*; supplementary figure S3).

### The distal domain of Po-exd is necessary for establishing the distal boundary of the patella

To assess the function of *Po-exd* in developing appendages, we performed embryonic RNAi via microinjection of double-stranded RNA (dsRNA) against *Po-exd* at two different points during *P. opilio* development (fig. 4). Embryos were injected at either stage five (early germband; henceforth, “early knockdown”) or stage seven (initial formation of prosomal limb buds; henceforth, “late knockdown”). Early knockdown of *Po-exd* incurred an array of major developmental defects that spanned irregular development of the head and failure to form the ocularium (prosomal protrusion bearing the median eyes); defects of antero-posterior segmentation and truncation of posterior segments and segmental fusions (fig. 4*a*-4*e*; supplementary fig. S4). Segmental defects of the prosoma included asymmetric fusions of adjacent segments and irregular placement of appendages, which exhibited proximal truncations (fig. 4*d*, 4*e*). Embryos exhibiting this array of phenotypes did not survive to hatching. These loss-of-function phenotypes closely parallel known phenotypic spectra for *exd* loss-of-function experiments in mandibulate arthropods, suggesting that the function of *exd* is conserved across arthropods with respect to early development (Rauskolb et al. 1995; Mito et al. 2008; Bruce and Patel 2020). Abnormal morphogenesis, including aberrant appendage localization, without homeotic transformations, is consistent with previous studies demonstrating that Hox genes require nuclear Exd and Hth proteins as cofactors for proper specification of the body wall, while Hox transcription factors operate independently to specify appendage identity (Smith and Jockusch 2014). Likewise, appendage truncations are consistent with independent *exd* and *hth* specification of proximal appendage identity (Smith and Jockusch 2014).

**Figure 4.**
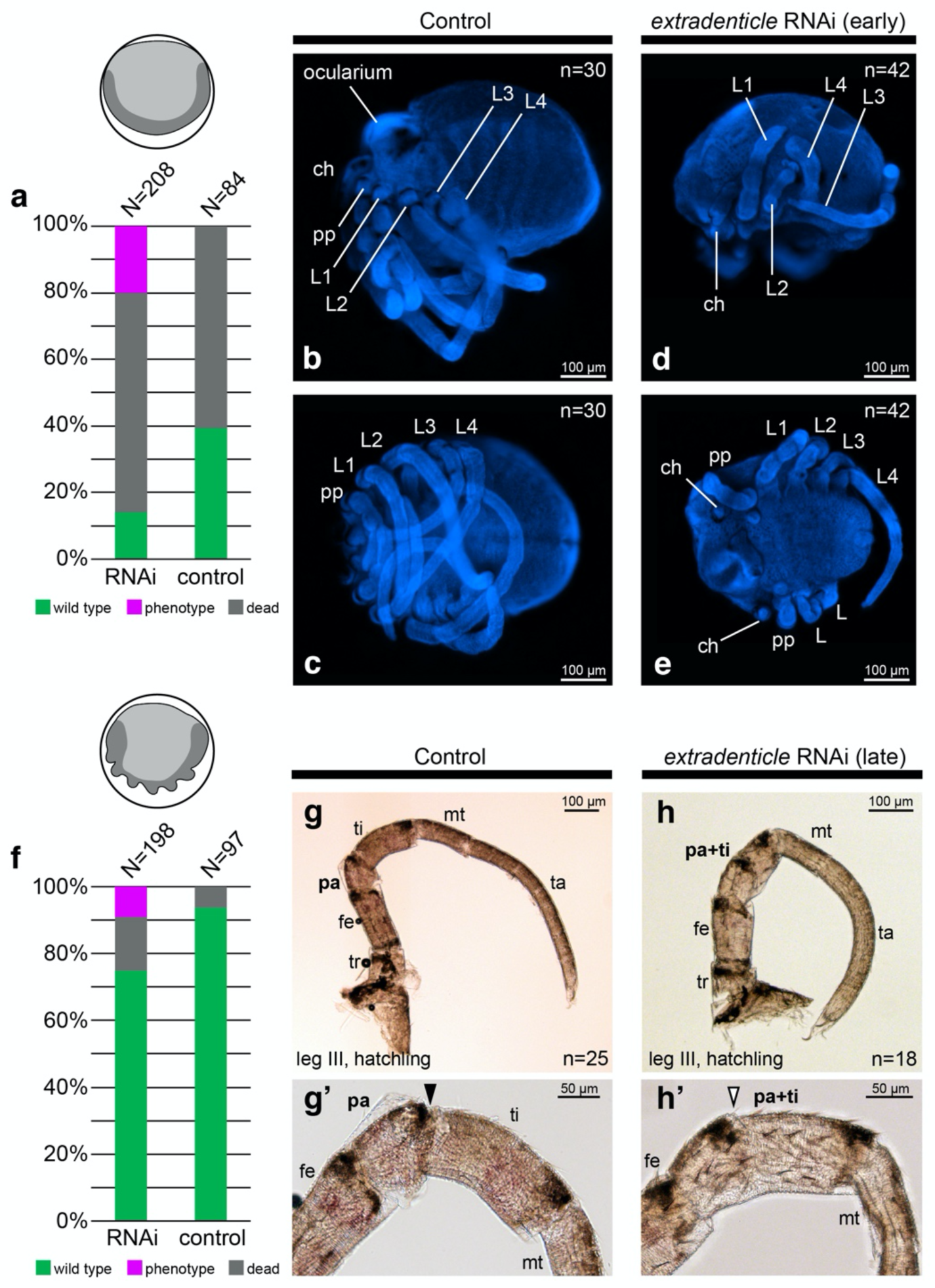
RNAi against *Po-exd* incurs a fusion at the patella-tibia joint in *P. opilio.* Icons indicate morphology of early and late RNAi embryos. Dark grey corresponds to the germband. (**a**) Distribution of outcomes following early *Po-exd* RNAi or negative control injections. (**b**-**e**) Early knockdown of *Po-exd.* (**b**) Negative control embryo, lateral view. (**c**) Same embryo as in (**b**), ventral view. (**d**) *Po-exd* RNAi embryo, lateral view. (**e**) *Po-exd* RNAi embryo, ventral view. Note posterior truncation and proximal leg defects in RNAi embryos. (**f**) Distribution of outcomes following late *Po-exd* RNAi or negative control injections. (**g**-**h’**) Late knockdown of *Po-exd.* (**g**) Leg three of negative control hatchling. (**g’**) Same hatchling as in (**g**), showing magnification of patella-tibia joint. (**h**) Leg three of *Po-exd* RNAi hatchling exhibiting fusion of patella-tibia joint. Note the location of melanized cuticle at dorso-distal boundary of femur, patella, and tibia. (**h’**) Same hatchling as in (**h**), showing magnification of fused patella-tibia joint. Black arrowhead: wild type patella-tibia joint. White arrowhead: fused patella-tibia joint. Abbreviations: ch, chelicera; pp, pedipalp; L1, leg one; L2, leg two; L3, leg three; L4, leg four; sp, spiracle; O2, second opisthosomal segment.

The effects of early *Po-exd* knockdown on the appendages precluded assessment of its role in specification of the patellar boundary. We therefore performed late knockdowns to interfere with *Po-exd* expression in stages where prosomal segmentation and body wall patterning had been established (Gainett et al. 2022) (fig. 4*f*-4*h*). Late knockdown of *Po-exd* resulted in embryos that were largely wild type in morphology and able to complete hatching but exhibited a fusion of the patella and tibia (fig. 4*h*). The interpretation of a fusion of segments, rather than a deletion, is substantiated by the retention of melanized patches at the distal and dorsal termini of the femur, patella, and tibia in hatchlings (fig. 4*h’*). These results suggest that the distal ring of *Po-exd* is required to form the distal segmental boundary of the patella.

### The distal domain of Po-exd is regulated by Notch signaling

The distribution of *Po-exd* transcripts during patella formation is reminiscent of the ring domains characteristic of leg segmentation genes in the Notch-Delta signaling pathway in both spider and insect model systems, such as *Delta, Serrate,* and *nubbin* (Rauskolb and Irvine 1999; Prpic and Damen 2009). To test whether the distal domain of *Po-exd* is under the control of the Notch signaling cascade, we performed late knockdown of *Po-Notch* (*Po-N*). The later delivery of dsRNA was selected to minimize early developmental defects that would impair appendage outgrowth.

*Po-N* RNAi embryos were broadly characterized by prominent segmental and neurogenic phenotypes (fig. 5; supplementary fig. S5, S6). As compared to the small, uniformly distributed rosettes of invaginating cells in the ventral neuroectoderm of wild type embryos, *Po-N* embryos instead demonstrate large, irregular pockets of cells, consistent with failed invagination of the neuroectoderm (fig. 5*b*; supplementary fig. S5*b*, S6). This outcome reflects a previously reported *Notch* RNAi phenotype in the spider *Cupiennius salei* (Stollewerk 2002), which resulted in failed invagination and yielded an accumulation of cells in the apical layer of the ventral ectoderm. While Notch signaling mediates cell interactions within these neural clusters, *Notch* is also required for lateral inhibition of the territory surrounding these clusters, limiting their size via activation of the *split* gene complex that represses neural identity (Chen et al. 2023). Consistent with this known activity, irregular development of the central nervous system could also be visualized via expression of *Po-dac*, which showed defects in developing segmental neuromeres (supplementary fig. S5*b*). The appendages of *Po-N* RNAi embryos displayed reductions in length and a wrinkled appearance compared to wild type appendages, consistent with previously reported *Notch* phenotypes in *C. salei* (Prpic and Damen 2009). Uniquely, *Po-N* RNAi also yielded consistent defects in the distalmost territory of most appendages, appearing as either early bifurcation or a central lacuna of tissue. *Po-N* RNAi embryos also exhibited abnormal development of the labrum, resulting in a smaller and circular structure, as opposed to the subtriangular wild type counterpart.

**Figure 5.**
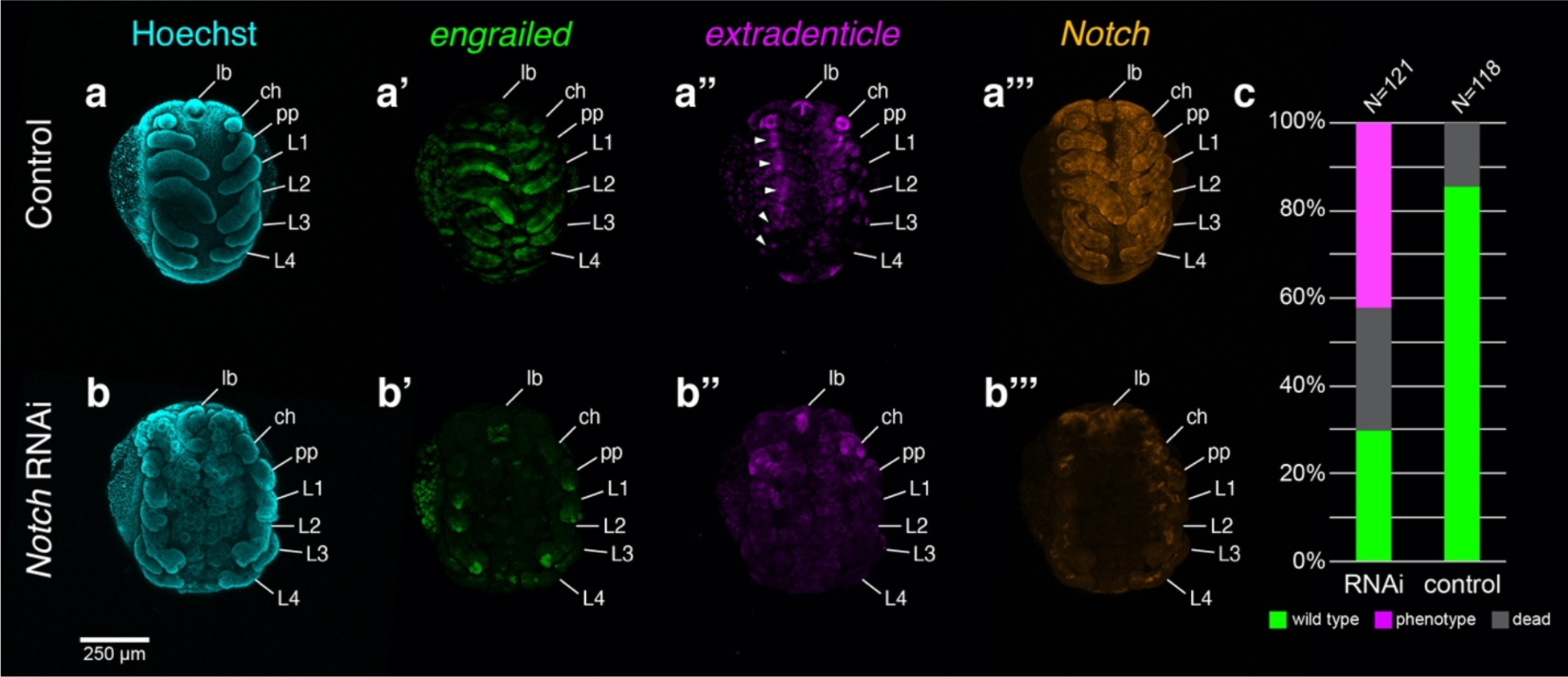
Knockdown of *Po-N* yields broad developmental defects and diminution of the *Po-exd* distal ring domain. (**a**) Stage 10 negative control embryo in ventral view with Hoechst counterstaining (cyan). (**a’-a’’’**) Same embryo as in (**a**) with expression of *Po-en* (green, **a’**), *Po-exd* (magenta, **a’’**), and *Po-N* (orange, **a’’’**). (**b**) Stage 10 *Po-N* RNAi embryo with Hoechst counterstaining. (**b’-b’’’**) Same embryo as in (**b**) with expression of *Po-en* (**b’**), *Po-exd* (**b’’**), and *Po-N* (**b’’’**). (**c**) Distribution of outcomes following *Po-N* RNAi or negative control injection. White arrowheads: distal ring domains of *Po-exd* in wild type embryo. Abbreviations: lb, labrum; ch, chelicera; pp, pedipalp; L1, leg one; L2, leg two; L3, leg three; L4, leg four.

To validate the incidence of segmental defects in *Po-N* RNAi embryos, we assayed expression of the segment polarity gene *engrailed* (*en*) (fig. 5; supplementary fig. S6). In wildtype embryos, *Po-en* was expressed in broad stripes in the posterior compartment of each segment. Expression was also detected in the posterior portion of each appendage along the length of the PD axis. Weak phenotypes in *Po-N* RNAi experiments retained *en* expression in each body segment, but expression was more diffuse and remained localized near the ventral midline (supplementary fig. S6*b*). In the appendages, the strongest *en* expression was found in small territories at the distal terminus, yet subsets of appendages still show weak expression throughout the PD axis. In severe phenotypes, *Po-en* was not detected in the ventral ectoderm, and only the distal tip expression is detectable in appendages supplementary fig. S6*c*). These results corroborate previous segmentation defects and aberrant deployment of segment polarity genes following abrogation of *Notch* signaling in arachnid and mandibulate embryos (Stollewerk et al. 2003; Pueyo et al. 2008; Eriksson et al. 2013).

To test the regulatory relationship of *Po-N* and the distal ring of *Po-exd,* we assayed *Po-N* RNAi embryos for expression of limb patterning genes. In *Po-N* RNAi embryos, expression of *Po-dac* in the medial territory and *Po-exd* in the proximal leg were both retained, suggesting that the establishment of these expression domains is not regulated by Notch signaling (supplementary fig. S5*b*, S7). By contrast, abrogation of *Po-N* resulted in marked diminution or abrogation of the distal *Po-exd* ring, consistent with the interpretation that this expression domain is regulated by the Notch segmentation cascade (fig. 5*b’’*; supplementary fig. S5*b*’’, S7*b*). These results closely reflect a previous experiment in the spider *C. salei,* wherein RNAi against the spider *Notch* homolog resulted in abrogation of the distal ring of *exd-1*, but had no effect on the expression of *dac, Dll,* or the proximal domain of *exd* (Prpic and Damen 2009). It was previously suggested that the distal *exd-1* domain could be uniquely operating downstream of *Notch* signaling, but since that work, the function of *exd-1* has not been investigated to date in any spider.

Taken together, these results support the interpretation that a distal, arachnid-specific expression domain of *exd* has acquired a novel segmentation function responsible for patterning the patella-tibia boundary.

### Late knockdown of Po-dac suggests an ancestral role for dac homologs in medial segmentation

In insects, *dac* is known to be required for leg morphogenesis and elongation of the medial appendage territory (Mardon et al. 1994). One possible interpretation of the spider *dac-2* segmentation phenotype is that it reflects an indirect effect of *dac* RNAi (i.e., limb outgrowth, rather than segmentation), and that this function was already present in single-copy *dac* homologs across Panarthropoda. A previous investigation of *dac* function in *P. opilio* yielded a canonical limb gap phenotype, with deletions of the femur and patella in the leg and palp as well as the proximal segment of the chelicera, precluding a test of this hypothesis (Sharma et al. 2013). We reasoned that *dac* may continue to play a role in the elongation and outgrowth of these segments after their specification, a function that could be examined by interfering with *dac* expression later in development.

To test this reasoning, we performed early and late knockdown of *Po-dac*, following the same experimental strategy as for *Po-exd* (fig. 6). Early knockdown of *Po-dac* recapitulated the result of a previous experiment (fig. 6*c*, 6*d*). Late knockdown of *Po-dac* incurred a milder phenotypic spectrum, characterized by shortening and/or fusion of medial segments of the palps and legs, but no defects of proximal (coxa and trochanter) or distal (metatarsus and tarsus) segments (fig. 6*e*-6*g*). Fusions were interpretable based on pigmentation patterns of medial podomeres. The most severe condition observed in late knockdown embryos consisted of fusion of the trochanter through the tibia (fig. 6*g*).

**Figure 6.**
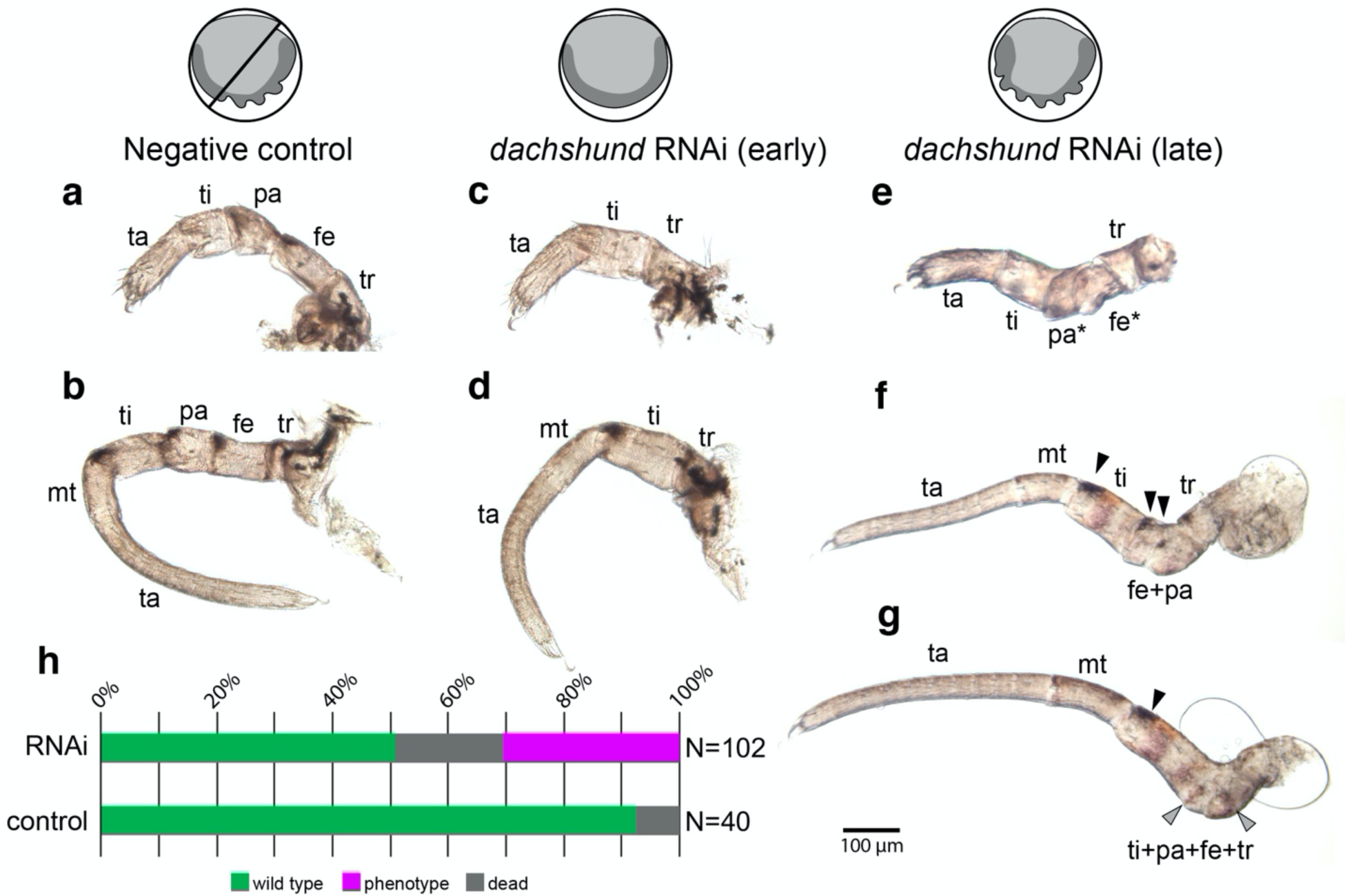
Late RNAi against *Po-dac* results in segmental fusions of medial podomeres. Icons indicate early and late RNAi embryos. Dark grey corresponds to the germband. (**a**) Pedipalp of negative control hatchling. (**b**) Leg three of negative control hatchling. (**c**) Pedipalp of early knockdown hatchling. (**d**) Leg three of early knockdown hatchling. (**e**) Pedipalp of late knockdown hatchling. (**f**) Leg three of late knockdown hatchling. (**g**) Leg four of late knockdown hatchling. (**h**) Distribution of outcomes following late *Po-dac* RNAi or negative control injection. Black arrowheads: melanized cuticle patches at dorso-distal boundary of femur, patella, and tibia. Grey arrowheads: traces of melanized patches in knockdown phenotypes. Asterisks indicate segments with aberrant morphology.

To assess the regulatory relationship between *Po-exd* and *Po-dac* in the patella-tibia boundary, we assayed a subset of late knockdown *Po-dac* RNAi embryos for expression of *Po-exd* (fig. 7). Late knockdown *Po-dac* embryos exhibited a visibly shorter and thickened medial palp and leg territory, consistent with the phenotype described for hatchlings from this experiment. Despite modest reduction of *Po-dac* expression, the spatial arrangement of *Po-dac* and the distal *Po-exd* domain was not altered, and the distal ring of *Po-exd* in the palps and legs was not visibly affected (fig. 7*b*, 7*c*). These results support the interpretation that *Po-dac* is not required for the maintenance of the distal *Po-exd* domain.

**Figure 7.**
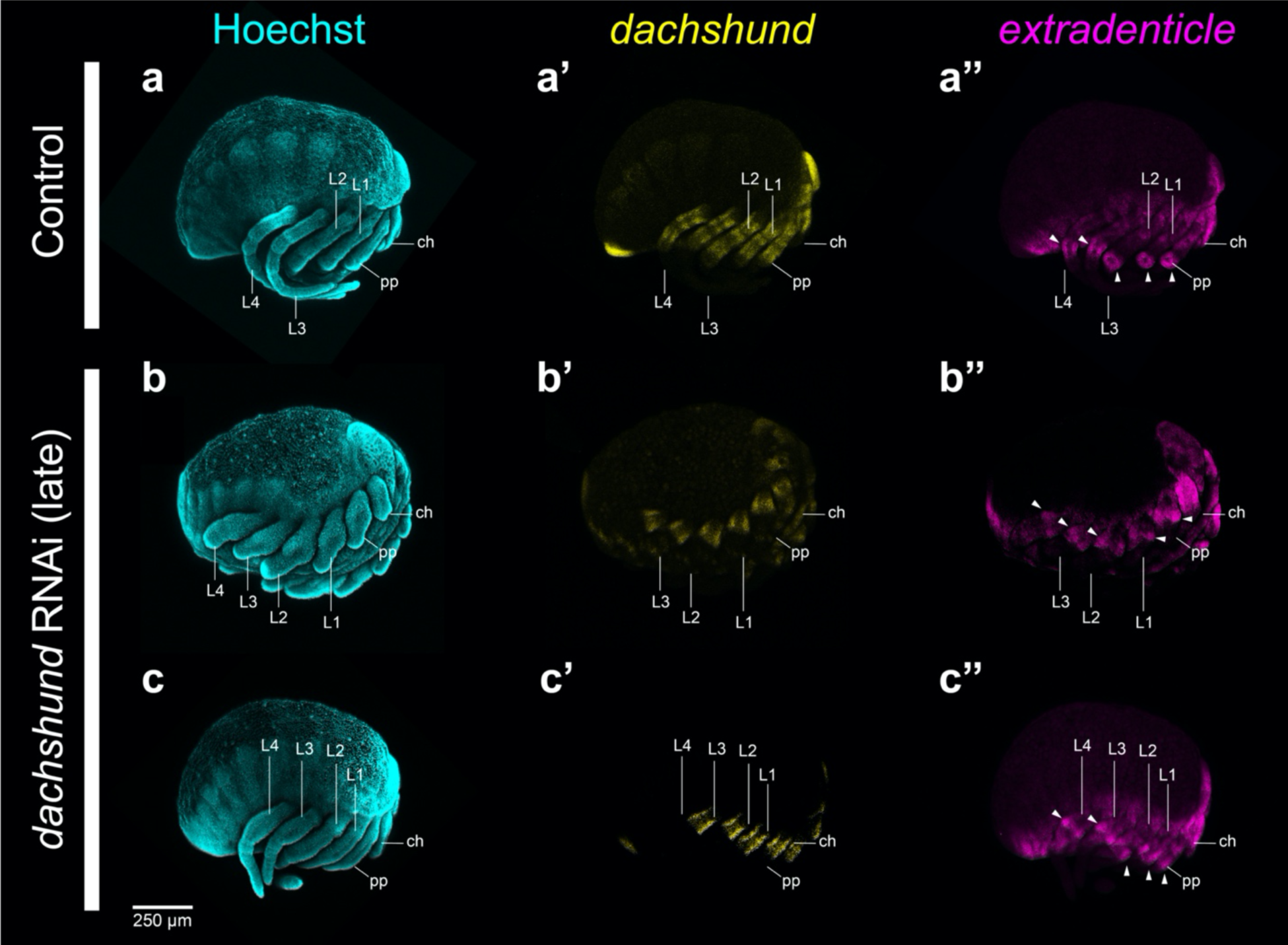
Late RNAi against *Po-dac* does not diminish distal expression of *Po-exd.* (**a**) Negative control embryo in lateral view, with Hoechst counterstaining (cyan). (**a’**-**a’’**) Same embryo as in (**a**) with single-channel expression of *Po-dac* (yellow, **a’**) and *Po-exd* (magenta, **a’’**). (**b**-**c’’**) *Po-dac* RNAi embryos in lateral view with Hoechst counterstaining (**b**, **c**), *Po-dac* expression (**b’**, **c’**), and *Po-exd* expression (**b’’**, **c’’**). Note the more severe medial appendage defects in (**b**) coincident with pronounced developmental delay. White arrowheads: distal ring domains of *Po-exd* expression in all embryos examined. Abbreviations: ch, chelicera; pp, pedipalp; L1, leg one; L2, leg two; L3, leg three; L4, leg four.

### Limb patterning gene expression dynamics in the sea spider Pycnogonum litorale

In contrast to arachnids, most sea spiders undergo pronounced indirect development, characterized by the hatching of a protonymphon larva with three larval limb pairs only, which correspond to the chelicera, the pedipalp, and the oviger (supplementary fig. S8*a*) (Vilpoux and Waloszek 2003; Brenneis et al. 2017). During postembryonic development, body segments bearing leg pairs form sequentially at the posterior terminus. Each of the four legs undergoes a similar sequence of developmental stages. Therefore, we primarily focused on leg 1 in instars II to IV.

The primordium of leg 1 is first discernible in late instar II (supplementary fig. S8*a*). Distally, it displays *Pl-Dll* expression, which is proximally bordered by a narrow ring-like *Pl-dac* territory. One to two single *Pl-dac-*positive cells are located in the distal limb bud tip. The entire primordial bud expresses *Pl-exd* at low levels. A ring-like domain with stronger signal intensity partially overlaps with the *Pl-dac* ring and extends slightly further distally (supplementary fig. S8*a*). In early instar III, leg 1 resembles a proper limb bud that projects posteriorly (fig. 8*a*; supplementary S8*b*). *Pl-Dll* is still strongly expressed in the distal tip, showing a gradual decrease of intensity toward the medial limb bud portion. The ring-like *Pl-dac* territory is more pronounced than in the preceding instar, its distal boundary overlapping slightly with the *Pl-Dll* domain. In the limb bud’s tip, the single *Pl-dac-*positive cells persist. *Pl-exd* expression reaches from the bud’s proximal base up to half of its length, being strongest along its distalmost portion that extends beyond the distal border of the *Pl-dac* territory (fig. 8*a*).

**Figure 8.**
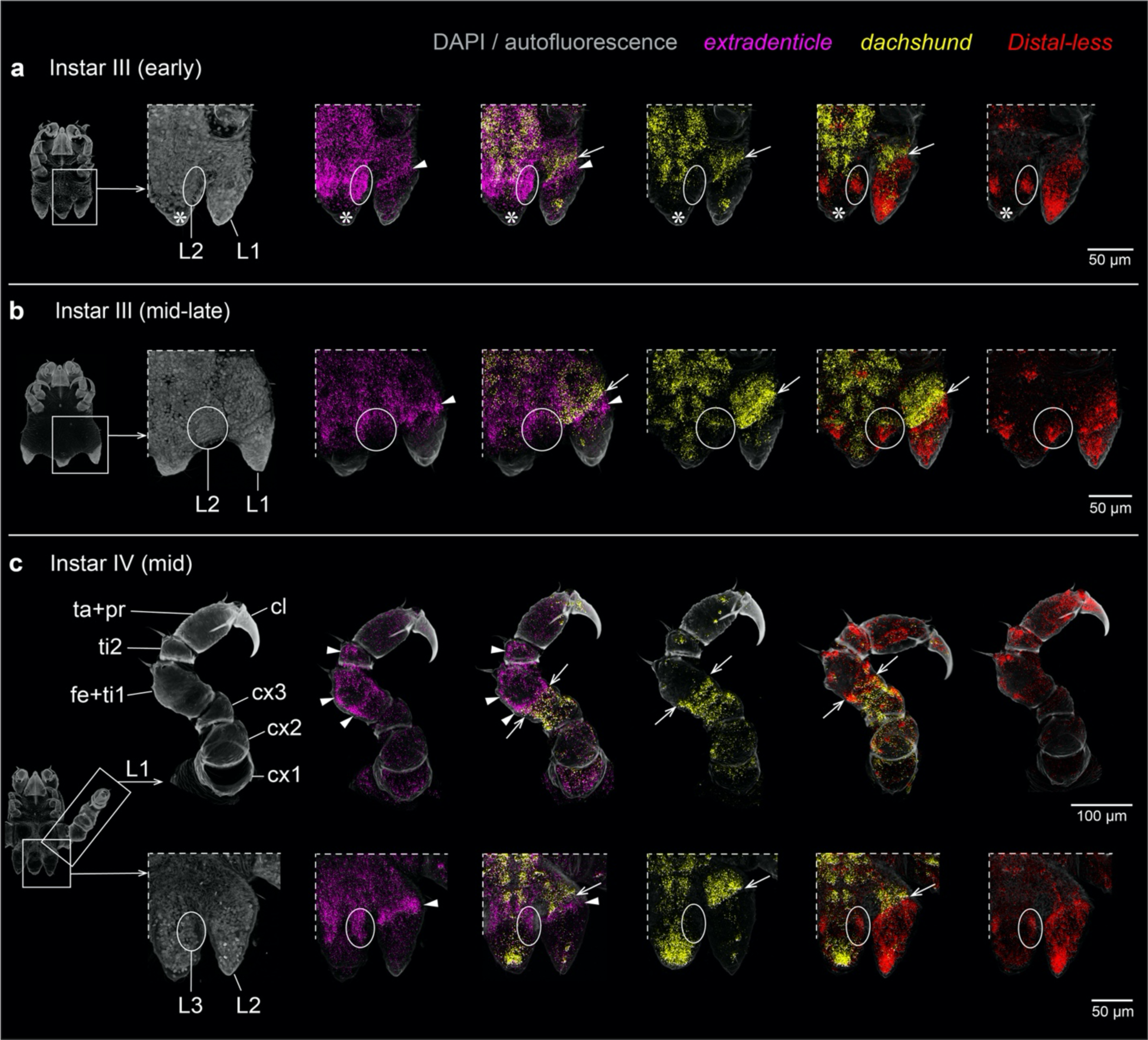
*Pl-exd* is expressed in a ring-like domain distal to *Pl-dac* expression in the developing legs of the sea spider *P. litorale*. All images apart from upper row in (**c**) show a ventral detail of the posterior body pole as indicated to the left of each row. Arrows: distal boundary of *Pl-dac* domain. Arrowheads: *Pl-exd* expression distal to the *Pl-dac* domain. Ovals and circles: limb bud primordia hidden under the cuticle. Asterisks indicate region of damaged tissue at the tip of the posterior body pole. (**a**) Detail of leg 1 bud and leg 2 primordium in early instars III. (**b**) Detail of leg 1 bud and leg 2 primordium in mid- to late-stage instars III. (**c**) Upper row: dissected leg one of mid-stage instars IV. Lower row: detail of leg 2 bud and leg 3 primordium in mid-stage instar IV. Abbreviations: cx, coxa; fe+ti1, femur-tibia 1 precursor; L, leg; mc, main claw; ta+pro, tarsus-propodus precursor; ti2, tibia 2.

In advanced stages of instar III, the tissues of leg 1 have considerably expanded along the PD axis, but remain confined under the limb bud’s rigid cuticle. This leads to a telescope-like folding and partial coiling of the leg tissue along the PD axis (fig. 8*b*; supplementary fig. S8*c*). Spatial relationships of the gene expression domains remain unchanged, with *Pl-exd* featuring a ring-like territory distal to the *Pl-dac* ring. Furthermore, the distal *Pl-Dll* domain includes scattered clusters with higher expression levels (fig. 8*b*).

With the molt toward instar IV, leg 1 becomes functional and is comprised of six podomeres (fig. 8*c*) (Sánchez and López-González 2010; Brenneis et al. 2017). At this point, observed gene expression domains can be unequivocally assigned to discrete podomeres. Directly after the molt, the expression signal for all three genes was observed to be relatively weak, presumably due to the considerable stretching of the largely single-layered ectodermal cell layer, concomitant with the rapid leg extension (supplementary fig. S8*d*). In mid- and late-stage instars IV, signal strength was observed to increase again (fig. 8*c*; supplementary fig. S8*e*). *Pl-Dll* is strongly expressed in the main claw and the tarsus-propodus precursor. In the more proximal podomeres, scattered domains of stronger expression are primarily located in tissue underlying cuticular sensilla, suggesting a potential role of *Pl-Dll* in sensory cells. *Pl-dac* expression demarcates a medial domain that includes coxa 3 and the proximal portion of the femur-tibia 1 precursor, its distal boundary marking the border of the two future podomeres (fig. 8*c*). More distally, *Pl-dac* is upregulated only in scattered single cells, which may represent sensory cells. *Pl-exd* displays low expression levels along the entire PD axis, but the proximal coxa 1 as well as the distal two thirds of the femur-tibia 1 precursor and the entire tibia 2 show elevated expression (fig. 8*c*). In the femur-tibia 1 precursor, the proximal expression boundary overlaps slightly with the distal boundary of the *Pl-dac* domain.

In the early stages of leg 2 and leg 3 development included in our experimental series, spatial relationships of gene expression correspond well to leg 1, with the additional observation that *dac* is the last of the three genes surveyed to be expressed in the earliest recognizable limb bud primordia (fig. 8; supplementary fig. S8*b*-S8*e*). Taken together, the consistent expression of *Pl-exd* distal to the medial *Pl-dac* territory during *P. litorale* leg development closely mirrors the situation in *P. opilio.* Beyond this, the specific *Pl-dac* and *Pl-exd* expression domains in the segmented leg of instar IV provides the first molecular developmental arguments for the homology of these pycnogonid leg podomeres with those of other chelicerates. This includes the identification of the sea spider tibia 1 as the homolog of the patella.

### Limb patterning gene expression dynamics in non-model arachnids

We next examined the spatial relationships of *exd* and *dac* homologs in a subset of arachnid orders wherein the incidence of a patellar homolog has been historically disputed (fig. 9). In the developmental transcriptome of the pseudoscorpion *Pselaphochernes parvus*, we detected two copies of *exd* and *dac*, consistent with the recent placement of Pseudoscorpiones within Arachnopulmonata and the inference of a shared genome duplication (Ontano et al. 2021). In embryos of *P. parvus*, we detected expression domains of *dac* and *exd* homologs that largely reflected the patterns previously described for other arachnopulmonates (Prpic and Damen 2003; Nolan et al. 2020). Expression of *Pp-dac-1* was observed in the femur and proximal patella of the palp and legs, whereas *Pp-exd-1* was observed as three domains: (a) a proximal domain corresponding to the body wall, coxa, trochanter, and proximal femur; (b) a median domain intercalating the *Pp-dac-1* expression domains at the femoro-patellar boundary; and (c) a distal domain corresponding to the distal patella (fig. 9*a*). By contrast, the expression domains of *Pp-dac-2* and *Pp-exd-2* were more weakly detected in later developmental stages and comprised two overlapping rings at the femoro-patellar boundary (fig. 9*b*, 9*c*).

**Figure 9.**
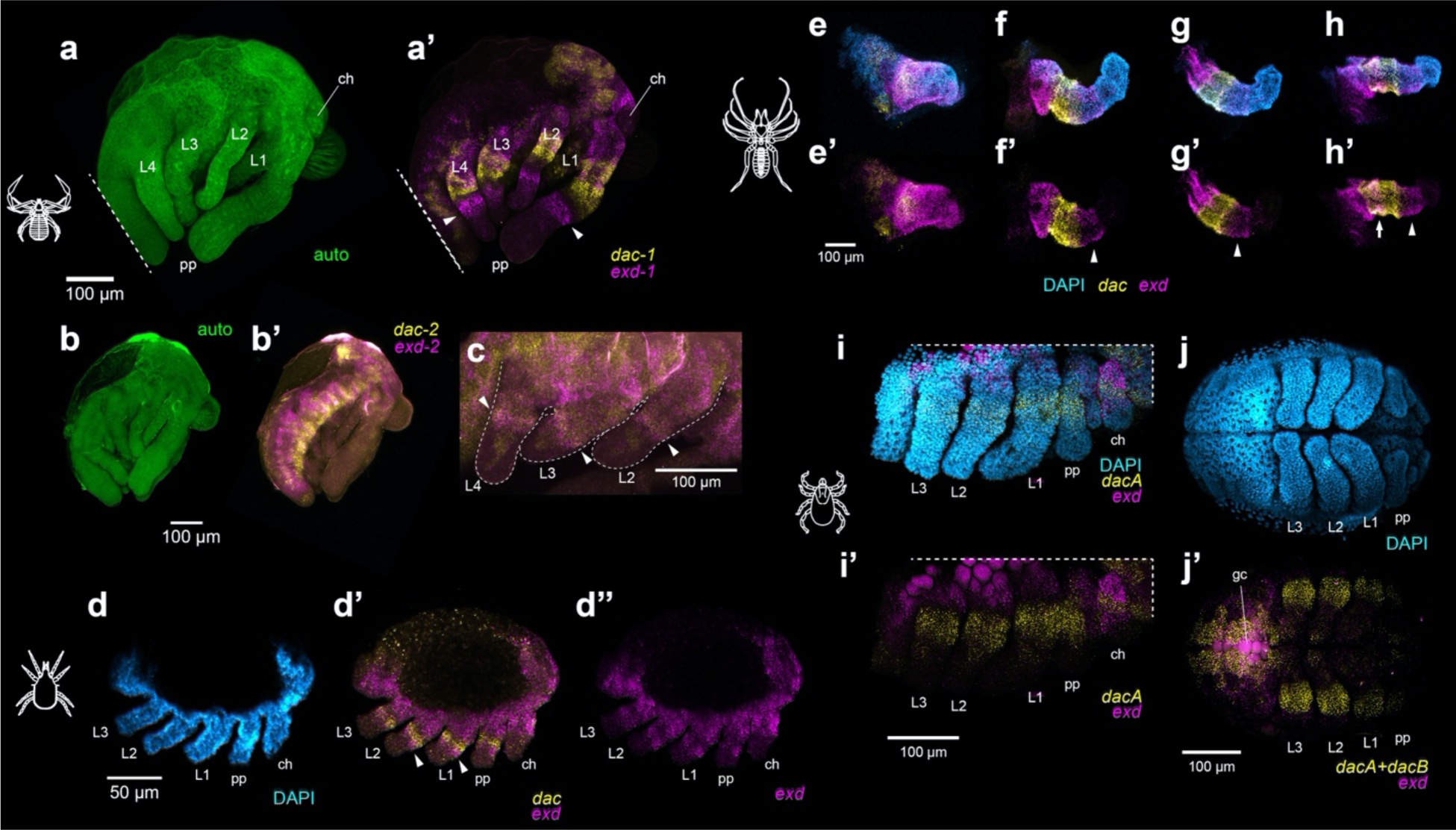
Surveys of *exd* and *dac* expression across arachnid orders with disputed patellar homologs. (**a**) Embryo of the pseudoscorpion *Pselaphochernes parvus* visualized via cuticular autofluorescence (green). (**a’**) Same embryo as in (**a**) with multiplexed expression of *Pp-exd-1* (magenta) and *Pp-dac-1* (yellow). (**b**) Autofluorescent visualization of *P. parvus* embryo. (**b’**) Same embryo as in (**b**) with expression of *Pp-exd-2* and *Pp-dac-2.* (**c**) Magnified view of *Pp-exd-2* and *Pp-dac-2* expression in the legs of an older *P. parvus* embryo. (**d**) Limb bud stage embryo of the acariform mite *Archegozetes longisetosus* with Hoechst nuclear counterstaining (cyan). (**d’**, **d’’**) Same embryo as in (**d**) with multiplexed expression of *Al-exd* (magenta) and *Al-dac* (yellow) (**d’**), or single-channel expression of *Al-exd* (**d’’**). (**e**-**h**) Limb mounts of the solifuge *Titanopuga salinarum* at leg elongation stage with nuclear counterstaining (cyan), and expression of *Ts-exd* (magenta) and *Ts-dac* (yellow). (**e**, **e’**) Chelicera. (**f**, **f’**) Pedipalp. (**g**, **g’**) Leg one. (**h**, **h’**) Leg three. Note additional domain of overlapping *dac* and *exd* in the basifemur of leg three (unique to legs three and four; white arrow). White arrowheads indicate an *exd* boundary distal of *dac.* Abbreviations: ch, chelicera; pp, pedipalp; L1, leg one; L2, leg two; L3, leg three; L4, leg four. (**i**) Stage 9 embryo of the tick *Ixodes scapularis* with Hoechst nuclear counterstaining (cyan), *Is-dacA* expression (yellow), and *Is-exd* expression (magenta). (**e’**) Same embryo as in (**e**) without nuclear counterstaining. (**j**) Stage 10 embryo of *I. scapularis* with nuclear counterstaining. (**j’**) Same embryo as in (**j**) showing multiplexed expression of *Is-dacA* and *Is-dacB* (yellow), and *Is-exd* (magenta). Note autofluorescence of yolk and germ cells (gc) in magenta channel in (**i’**, **j’**).

In the acariform mite *Archegozetes longisetosus*, the single-copy homologs of *dac* and *exd* were expressed comparably to their *P. opilio* counterparts. A ring of *Al-exd* expression in juxtaposition with, and distal to, the *Al-dac* domain was observed in the palp and legs of the hexapodous larva at the limb bud stage (fig. 9*d*).

In the appendages of the solifuge *Titanopuga salinarum*, expression of the single-copy *Ts-exd* and *Ts-dac* homologs was largely consistent with expression patterns observed in other surveyed taxa (fig. 9*e*-9*h*). The chelicerae exhibit strong expression of *Ts-exd* in both podomeres (fig. 9*e*, 9*e’*). A domain of *Ts-dac* was detected in the proximal territory of the chelicera, as has previously been described for the mite *A. longisetosus* (Barnett and Thomas 2013). In the pedipalps and first two pairs of walking legs, *Ts-dac* was expressed in the presumptive femoral segment, whereas *Ts-exd* was expressed in both a proximal territory encompassing the coxa and trochanter, and a distal ring domain adjacent to the medial *Ts-dac* expression (fig. 9*f*-9*h*). This distal expression suggests that the “post-femur” of some authors can be homologized to the patella (Millot 1949; Punzo 1998), consistent with the nomenclature of Shultz (1989). The third and fourth walking legs also exhibited the conserved proximal and distal domains of *Ts-exd*, and medial domain of *Ts-dac* (fig. 9*h*, 9*h’*). In contrast to the pedipalp or the first two leg pairs, an additional domain comprised of overlapping *Ts-dac* and *Ts-exd* expression was observed in the presumptive basifemur (*sensu* Shultz, 1989). Presence of the non-overlapping distal *Ts-exd* domain again supports retention of a patellar homolog and refutes previous interpretation of a subdivided trochanter and a subdivided femur condition in the third and fourth solifuge leg (Millot 1949; Punzo 1998).

In the genome assembly of the tick *Ixodes scapularis*, we discovered two copies of *dac* and one copy of *exd*. Gene tree analysis of the tick *dac* paralogs suggested independent origins with respect to the arachnopulmonate copies. When multiplexed in a single channel, the two *dac* copies (*Is-dacA*, *Is-dacB*) exhibited comparable dynamics with respect to the single-copy harvestman homolog, spanning the trochanter, basifemur, and distifemur (fig. 9*j*). Surprisingly, *Is-exd* expression was detected only in the body wall, coxa, and trochanter; we did not detect *Is-exd* expression as a ring distal to the *dac* paralogs in any stages surveyed (fig. 9*i*, 9*j*).

## Discussion

### A phylogenetically consistent mechanism for patellar origin

Developing mechanistic connections between genotype and phenotype is a fundamental goal of evolutionary developmental biology. Crucially, in comparative development contexts, the putative genetic mechanism underlying a novel trait must be compatible with the phylogenetic distribution of that trait. Here, we investigated the fit of a putative developmental mechanism for patellar origin using a phylogenetic test and showed that the phylogenetic distribution of *dac-2*, which is restricted to Arachnopulmonata, is inconsistent with the interpretation that the origin of this gene copy underlies the patterning of the patellar segment across Chelicerata. Paralleling this case, expression surveys of other paralogous genes in spider models have prompted inferences of neofunctionalization that exhibit similar mismatch of phylogenetic distribution between gene and trait. In one case, expression of *hth-1* reflects patterns found in arthropods broadly, whereas *hth-2* is expressed as a series of distal ring domains that vary across spider lineages (Turetzek et al. 2017). The authors took these expression differences to mean neofunctionalization of the duplicated *hth-2* copy, in addition to the inference that changes in the number of *hth-2* expression rings across spiders were mechanistically meaningful with regard to differences in appendage morphology across spider taxa. However, it was later shown that a similar division of expression domains is observed across both copies of *hth* in scorpions and whip spiders, while broader taxonomic sampling of *hth* has subsequently implicated its origin as a result of the arachnopulmonate whole genome duplication. (Sharma et al. 2012a; Gainett and Sharma 2020; Nolan et al. 2020). Thus, while the functional significance of their expression dynamics remain unknown, the incidence of the *hth-2* paralog is demonstrably not spider-specific, nor do the expression levels of *hth-2* paralogs correlate with specific leg phenotypes (Sharma 2023). As a second example, expression of one paralog of the paired box gene *Pax2* in the lateral eyes of spiders was taken to suggest subfunctionalization of ancestral pleiotropic functions in brain, appendage, and eye development, as well as a key role in differentiating the lateral eyes from the median eyes (Janeschik et al. 2022). However, a recent work refuted a role for *Pax2* as an eye marker specific to lateral eyes; the single copy homolog of *Pax2* is expressed in both the median eyes and vestigial lateral eyes of harvestmen, whereas both orthologs of *Pax2* in the scorpion are also expressed in median and lateral eye primordia (Gainett et al. 2024). These patterns suggest that the dynamics of *Pax2-1* in the developing eyes of spiders are a taxon-specific phenomenon and do not reflect the ancestral condition of *Pax2* homologs across Chelicerata.

These previous interpretations of neofunctionalization based on spider expression patterns have substantiated the perception that the morphological evolution of chelicerates has been largely shaped by whole genome duplication and neofunctionalization of paralogs, despite a dearth of functional studies that link new genes to new phenotypes in arachnopulmonates (Sharma 2023). Our work suggests instead that novel traits can be established by the rewiring of existing gene regulatory networks comprised of ancient genes. In support of this interpretation, we have demonstrated that the acquisition of a novel expression domain by the conserved appendage patterning gene *extradenticle* is essential for establishment of the patellar segment. While divergence of paralog expression domains across arachnopulmonates is a compelling phenomenon, understanding their role in the evolution of novel traits requires functional studies to substantiate hypotheses of neofunctionalization. As shown here in the case of *dac-2,* functional data must also be paired with broader taxonomic sampling and comparative gene expression, toward applying a phylogenetic test of putative genotype-phenotype connections. More generally, surveys outside the focal study taxon are essential to polarizing developmental phenomena. The increasing availability of genomic resources and functional techniques for diverse model species spanning animal diversity, as demonstrated herein through the first gene expression surveys for multiple chelicerate orders, foretells a more robust framework for future investigations of comparative developmental mechanisms.

### The developmental dynamics of arachnopulmonate dachshund are consistent with subfunctionalization

The coincidence of the phenotype incurred by knockdown of *Po-exd* and *dac-2* in the spider *P. tepidariorum* is remarkable and invites reinvestigation of *dac* in chelicerate appendage evolution. This coincident loss-of-function phenotype could be interpreted as a case of developmental system drift, wherein the divergence of genetic mechanisms across taxa does not affect a trait’s expression. Alternatively, the function of *dac-2* in spiders may reflect the subdivision of an older role for *dac* in establishing the segmental boundaries of medial leg segments, albeit indirectly in a Notch-independent manner, given retention of medial *dac* expression following abrogation of *Po-N* (fig. 5*b’*; supplementary fig. S7). Apropos, *dac* loss-of-function phenotypes in other arthropod models have suggested a role for *dac* in segmentation. As examples, loss-of-function *dac* mutants in *D. melanogaster* exhibit a fusion of the a5-arista joint in the antenna (Dong et al. 2001). Similarly, weak knockdown of *dac* results in fusion of the femur-tibia joint in the hemipteran *Oncopeltus fasciatus* (Angelini and Kaufman 2004), and larval RNAi against *dac* in the beetle *Tribolium castaneum* incurs fusions of the first three tarsal articles (Angelini et al. 2012). These phenotypes are consistent with the observation that *dac* null mutants exhibit increased cell death in the presumptive medial territory of leg imaginal discs of *D. melanogaster* (Mardon et al. 1994).

The loss of segment boundaries in late *Po-dac* RNAi embryos is consistent with phenotypic spectra of *dac* homologs across the arthropod tree of life and suggest a conserved role for *dac* in the morphogenesis, axis subdivision, and outgrowth of the medial appendage territory. With respect to *dac-2*, these results suggest that the previously described function of the spider *dac* duplicate copy does not reflect neofunctionalization (Turetzek et al. 2016), as much as a subdivision of the ancestral function between the two arachnopulmonate-specific daughter copies. Consistent with this interpretation, the phenotypic spectrum of *Po-dac* encompasses the phenotype exhibited by spider *dac-2* RNAi hatchlings. In addition, the irregular outgrowth at the fused patella-tibia boundary in *P. tepidariorum dac-2* RNAi hatchlings resemble morphogenetic defects described in insect *dac* RNAi experiments (Angelini and Kaufman 2004).

As a test of this interpretation, future experiments should reexamine *Notch* loss-of-function phenotypes to assess whether RNAi against *Notch* results in loss of the *dac-2* distal domain, as shown for *exd* (consistent with developmental system drift) or in retention of *dac-2* expression in the patella (consistent with subfunctionalization of the ancestral *dac* domains). Functional data for *dac-1* are also sorely needed to validate interpretation of either sub- or neofunctionalization. However, we add the caveat that inherent limits on the effectiveness of RNAi in *P. tepidariorum* may hinder functional investigations of leg patterning in that species. As an example, we trialed maternal RNAi against *exd-1* and *exd-2* in this study, following previous approaches to RNAi in *P. tepidariorum* (Akiyama-Oda and Oda 2006; Khadjeh et al. 2012; Setton and Sharma 2018). We observed no morphological defects in those experiments, with all embryos and hatchlings exhibiting wild type morphology (data not shown). Similarly, our efforts to evince the role of *dac-1* in several aspects of spider morphogenesis (e.g., leg patterning; neurogenesis; eye development) have repeatedly met with the same result in *P. tepidariorum* (data not shown).

### A patella was present in the common ancestor of Chelicerata

Together with existing expression data in spider and scorpion models (Prpic and Damen 2009; Nolan et al. 2020), the demonstration of a distal *exd* domain in harvestmen, acariform mites, solifuges, and pseudoscorpions underscores widespread conservation of patellar homologs across terrestrial arachnids. Retention of the distal ring in Pycnogonida likewise supports the presence of the patella in the common ancestor of Chelicerata (fig. 10), as previously advocated in some of the early morphological works (Snodgrass 1958; Dencker 1974; Schram and Hedgpeth 1978; Shultz 1989; Sabroux et al. 2023). As such, it seems that historical confusion and terminological mismatches in the arthropod literature may largely reflect an overreliance on morphometrics and perceived differences in podomere function, rather than strict interpretations of homology (reviewed in Shultz [1989]). Instead, broad congruence in both the number of appendage podomeres and patterns of muscle insertion sites across chelicerate orders supports a more unified nomenclature, with recognition of taxon-specific gains, subdivisions, or loss of podomeres (Shultz 1989).

**Figure 10.**
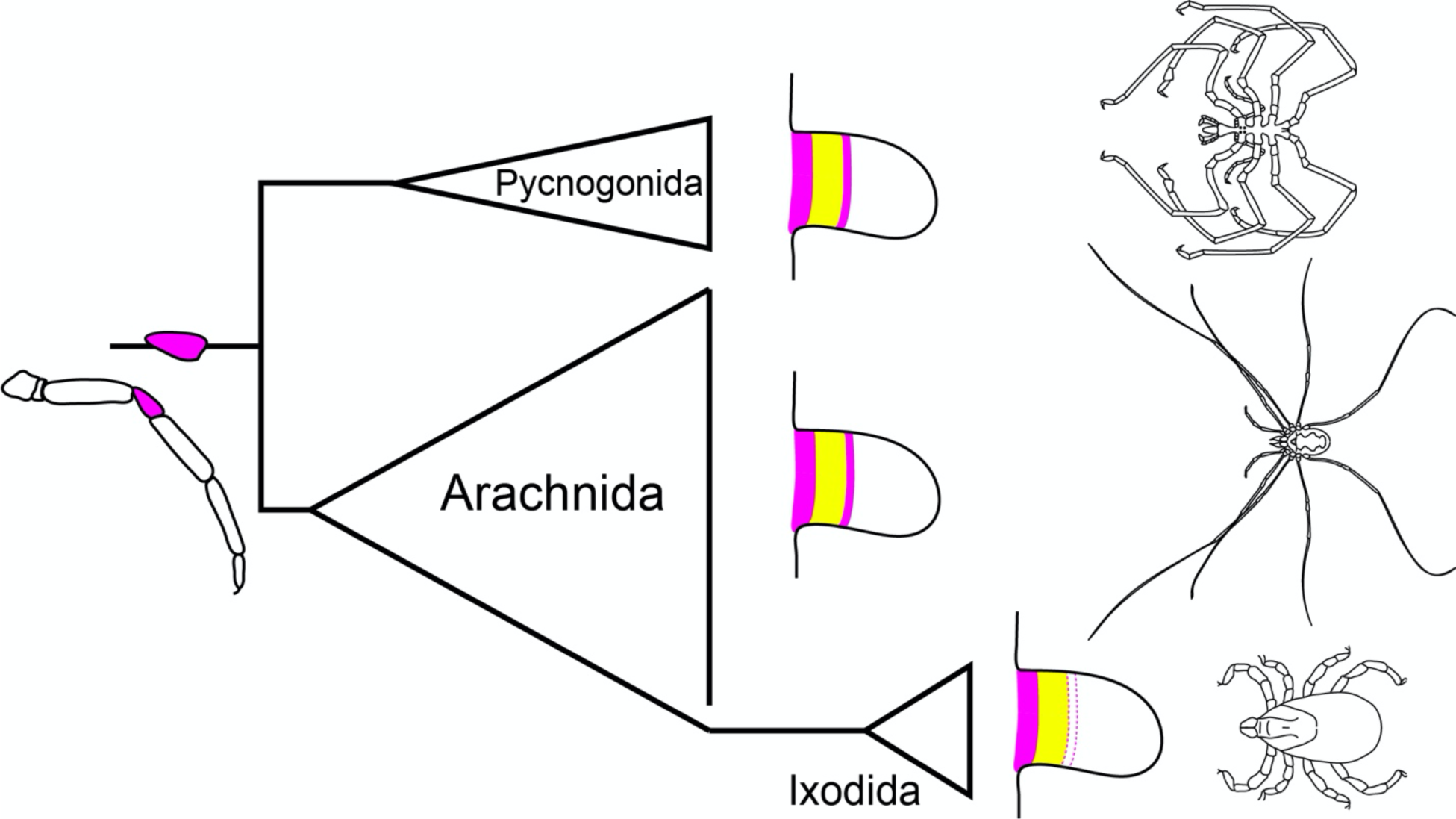
A distal ring domain of *exd,* abutting a conserved medial domain of *dac*, during embryonic appendage formation is responsible for the origin of the chelicerate patella. Presence of the distal ring domain in Pycnogonida and many chelicerate orders supports its presence in the chelicerate common ancestor. Lineage-specific loss of distal *exd* expression in ixodid ticks suggests loss of a patellar homolog.

The clear functional link between the distal ring domain of *exd* expression and establishment of the patellar segmental boundary highlights the utility of evolutionary developmental biology in resolving long-standing and often contentious interpretations of homology in anatomical structures. As other examples within Arthropoda, the function of retinal determination genes in harvestmen revealed vestiges of the lateral eyes and an additional pair of median eyes, whereas external morphology had long suggested retention of only a single pair of median eyes in this group (Gainett et al. 2024). Similarly, surveys of Hox gene expression have advanced the understanding of the “arthropod head problem”, a question of positional homology of head structures in this phylum. Such surveys famously established the homology of sea spider chelifores, arachnid chelicerae, and mandibulate (first) antennae as deutocerebral structures (Jager et al. 2006).

Despite improved resolution of patellar homology and evolutionary origin, the trait cannot be universally applied to all chelicerate taxa. The absence of distal *exd* expression in the parasitiform tick *I. scapularis* impels two alternative interpretations. First, it is possible that ixodid ticks have lost a patellar homolog, justifying the previous use of genu for the fifth podomere identity. Alternatively, Ixodida may have acquired either a novel genetic mechanism for patellar patterning or repurposed an existing appendage patterning gene to this end, exemplifying developmental system drift. The presence of two *dac* copies in the *I. scapularis* genome is particularly intriguing in this regard and invites interrogation of divergent paralog function following tandem duplication. The recent advent of gene editing tools in *I. scapularis* offers a promising vehicle for functional investigation of *dac* function in this system (Sharma et al. 2022). Notably, two other chelicerate orders have occasionally been suggested to lack patellar homologs. The pedipalps and walking legs of Palpigradi (microwhip scorpions) are thought to possess a genu in the identical PD axis position of the patella in other groups (van der Hammen 1982), whereas Ricinulei (hooded tick spider) pedipalps place the femur in that position, by nature of possessing two trochanters (Pittard and Mitchell 1972). The small size, difficulty of collection, and dearth of functional resources for these organisms presently prevents analysis of segmental homologies. Yet, recent phylogenetic affinities of these groups may yield more parsimonious hypotheses. The most comprehensive analysis of higher-level chelicerate relationships consistently recovered a sister group relationship of palpigrades and the clade containing both Acariformes and Solifugae with support (Ballesteros et al. 2022). As such, it is likely that palpigrades retain a patellar homolog, rather than genu, given its presence in both sister taxa. But in the same analyses, Ricinulei are recovered as sister group to Parasitiformes, and without resolution of segmental affinity in *I. scapularis,* suggesting that Ricinulei may also lack a patellar homolog.

Beyond the homology of the patella, chelicerate podomeres remain rife with unresolved homology disputes and questionable ancestral states (supplemental table S1, S2). The most prominent example is the condition of the femur, which is retained as a single podomere in most extant lineages (Arachnopulmonata, Opiliones, Palpigradi, Xiphosura), but is subdivided into basifemur and distifemur in groups such as Parasitiformes and Acariformes. Likewise, presence of an additional podomere in walking legs III and IV in Solifugae and Ricinulei is often described as either an additional trochanter or femur; the identification of a novel overlapping domain of *dac* and *exd* in legs III and IV of a solifuge embryo in this study (compare fig. 9*g* to fig. 9*h*) provide the first clues of how this additional podomere may be patterned. Reconstruction of the ancestral condition is further confounded by the additional proximal segmentation in Pycnogonida, with many taxonomists inferring the presence of three coxae and a single femur. This scheme therefore suggests an undivided femur as the likely ancestral chelicerate condition. Here, we have demonstrated strong expression of *exd* in the distal compartment of the still undivided femur-tibia 1 precursor podomere of the *P. litorale* instar IV, homologizing the proximal tibia 1 of sea spiders with the patella. This placement of the patella as the fifth podomere along the PD axis is reminiscent of the placement of the patella in Solifugae or non-oribatid Acariformes, groups possessing a subdivision of the femur. Application of this podomere terminology therefore aligns the three coxae and single femur of pycnogonids with coxa, trochanter, basifemur, and telofemur, potentially supporting the subdivision of femora as the ancestral state in Chelicerata. Descriptions of fossilized chelicerate lineages reinforce this interpretation. Unlike extant horseshoe crabs that possess five post-coxal segments, extinct synziphosurines, such as *Offacolus* and *Dibasterium,* retain six post-coxal segments (Sutton et al. 2002; Briggs et al. 2012). Many eurypterid taxa also possess additional segmentation in their third and fourth walking legs, represented as either a bipartite femur or trochanter (Selden 1981). Recent total evidence analysis placing Eurypterida and synziphosurines as part of a clade within the larger Merostomata (Ballesteros et al. 2022) further suggests secondary fusion of proximal segments in extant horseshoe crabs.

Patterns of muscle attachments also lend credence to the inference that the ancestral femur was subdivided (see Shultz [1989]). In *Limulus* (Xiphosura), Araneae, and Amblypygi, two muscles arise in the anterior trochanter, span the trochanter-femur joint diagonally, and insert in the proximoposterior femur. Yet, these muscles lack tendon insertions, obscuring their function. Both have been homologized with the depressor muscle spanning the basifemur-telofemur joint of Solifugae. Likewise, Amblypygi and Schizomida both possess a muscle arising in the anterior of the trochanter that inserts distally into the dorsal midline of the femur, without means of a tendon. This muscle has been homologized with the basifemur-telofemur levator in Solifugae. These unusual patterns are suggestive of a former mobile joint that has secondarily fused in groups like extant tetrapulmonates. Certain members of the extinct Trigonotarbida (putatively sister to Tetrapulmonata; Ballesteros et al. 2022) and extinct lineages of Araneae, Amblypygi, and Uropygi also possess appendages that have been reconstructed with an additional podomere between trochanter and femur (Shear et al. 1987).

Given the comparative framework, new study systems, and genomic resources established in this study, future investigations are well poised to address the ancestral condition of proximal segmentation in Chelicerata. Such investigations should prioritize identification of transcription factors that establish basi- and telofemoral identities in model taxa, with the goal of surveying the same mechanisms in groups like Pycnogonida, Solifugae, and Acariformes to polarize the ancestral condition across the chelicerate tree of life.

## Materials and Methods

### RNA Sequencing

Brooding females and nymphal stages of the pseudoscorpion *Pselaphochernes parvus* were collected from compost bins in Madison, Wisconsin (43°04′43.8″N, 89°23′12.1″W) in July 2022 and June 2023 (voucher specimens lodged in Western Australian Museum). A subset of eggs and juveniles were fixed in TRIzol TriReagent (Thermofisher) for RNA extraction. For the solifuge *Titanopuga salinarum,* three juveniles were collected in Córdoba, Argentina (64°48′S, 30°02′W) in December 2023 and brain tissue was dissected in phosphate buffered saline (PBS) and fixed in RNA*later,* with subsequent transfer to TRIzol. Total RNA was extracted from these samples and purified mRNA was sequenced on an Illumina NovaSeq platform at the UW-Madison BioTechnology Center, following our previous strategy for library preparation (Ontano et al. 2021). Transcriptomic assembly was performed using Trinity v. 2.15 (Grabherr et al. 2011); for the solifuge, the reads obtained in this study were combined with previous developmental transcriptomes to generate a new assembly (Gainett et al. 2023). A developmental transcriptome of the sea spider *P. litorale* was previously generated from pooled embryos and postembryonic instars reared in a laboratory culture at Greifswald University, Germany (Brenneis et al. 2023). All developmental stages were fixed and stored in RNAlater for RNA extraction.

### Bioinformatics and Phylogenetic Analysis

Homologs of *extradenticle, dachshund, Notch,* and *engrailed* were identified in the genome of *Phalangium opilio* (Gainett et al. 2021) and sequence identities confirmed by alignment against transcripts from previous studies in this species (Sharma et al. 2012a; Sharma et al. 2012b), as well as using SMART-BLAST. Homologs of *extradenticle* and *dachshund* were additionally extracted from the transcriptomes of the sea spider *Pycnogonum litorale* (Brenneis et al. 2023), the pseudoscorpion *P. parvus* (this study), and the solifuge *Titanopuga salinarum* (Gainett et al. 2023); this study); and from the genomes of the mite *Archegozetes longisetosus* (Brückner et al. 2022) and the tick *Ixodes scapularis* (Nuss et al. 2023). For *dac* and *exd,* peptide translations of nucleotide sequences were added to a previous alignment of panarthropod homologs (Nolan et al. 2020; Ontano et al. 2021) and multiple sequence alignment was performed *de novo* with CLUSTAL Omega (Sievers and Higgins 2018). Gene trees were inferred using IQ-TREE v. 1.6.12 (Nguyen et al. 2015) with an LG+I+G substitution model and nodal support was estimated using 1000 ultrafast bootstrap replicates. Alignments are provided in supplementary files S1 and S2. Gene trees are provided in supplementary file S3 and S4.

### Embryo Collection, Fixation, and In Situ Hybridization

Embryos were fixed and assayed for fluorescent detection of gene expression following established or minimally modified protocols, as detailed previously for *P. opilio* (Gainett et al. 2021) and *A. longisetosus* (Barnett and Thomas 2013). Adult females of *T. salinarum* were collected from the same locality as the juveniles used in RNA sequencing. Females were housed and resulting embryos fixed and assayed as in Gainett et al. (2023).

For *P. parvus,* embryos were dissected out of the broodsac with fine forceps and transferred to 2 mL Eppendorf tubes. Embryos were fixed in a solution of 3.2% paraformaldehyde in 1× PBS for 20 minutes, followed by washes in 1× PBST (0.1% Tween-20) and gradual dehydration into 100% methanol.

For *I. scapularis,* pathogen-free, unfed adults were acquired from Oklahoma State University Centralized Tick Rearing Facility and were maintained in an environmental chamber at 20°C and 95% relative humidity (RH) at the University of Nevada-Reno. Adult ticks were fed on New Zealand White rabbits (Reyes et al. 2020) until fully engorged. Gravid females were collected and placed into individual transparent containers for egg-laying at 20°C and 95% RH. Oviposition began 6-14 days after collection and eggs were harvested daily into 1.5 mL microcentrifuge tubes. To fix, eggs were transferred to a 70µm cell strainer (EASYstrainer, Greiner Bio-One) and washed for three minutes in 8% sodium hypochlorite while agitating with a paintbrush. Eggs were subsequently washed three times with deionized (DI) water. Using a paintbrush, the eggs were then transferred into PCR tubes with 100µL of DI water and incubated at 90°C for three minutes followed by snap cooling at -20°C for an additional three minutes before thawing at ambient temperature. DI water was then removed and replaced with a 1:1 ratio of 4% paraformaldehyde and heptane. Eggs were allowed to fix at ambient temperature on a vortex mixer for one hour or overnight. The paraformaldehyde layer was removed and replaced with an equal volume of 100% methanol and tubes were vigorously shaken for one to two minutes. Following fixation, embryos were stored at 4°C until ready for use. All procedures involving animal subjects were approved by the Institutional Animal Care and Use Committee (IACUC) at the University of Nevada-Reno (IACUC #21-01-1118-1).

For *P. litorale,* postembryonic instars were reared in the in-house laboratory culture at the Animal Facility at University of Vienna. Instars II to IV were collected from their host, the hydrozoan *Clava multicornis.* After relaxation for 1-2 minutes in freshly carbonated artificial seawater (ASW; 32‰), specimens were fixed in 4% paraformaldehyde in ASW for 1 hour at ambient temperature, washed two times in ASW, gradually transferred into 1x PBS, followed by gradual dehydration into 100% methanol. Samples were stored at -20°C until further use. Prior to HCR-FISH, the instars were gradually rehydrated into 1x PBS and exposed to 5-10 brief pulses in a bath ultrasonicator to enhance cuticle permeability.

Probe design for hybridization chain reaction (HCR) consisted of 12-30 probe pairs, depending upon the length of the available template sequence. Probes were designed either with the HCR Probe Maker tool (Kuehn et al. 2022), or via submission of CDS to Molecular Instruments. Input CDS templates for proprietary Molecular Instrument probe design and probe sequences obtained from the HCR Probe Maker tool for all six chelicerate species are provided in supplemental file S5 (supplementary tables S3-S19).

### Cloning of Orthologs, dsRNA Synthesis, and RNAi

Fragments of *Po-exd* and *Po-N* were amplified using standard PCR protocols and cloned using a TOPO TA Cloning Kit using One Shot Top10 chemically competent *Escherichia coli* (ThermoFisher) following the manufacturer’s protocol, and their PCR product identities were verified via Sanger sequencing with M13 universal primers. A plasmid containing a sequence of *Po-dac* was available from a previous study (Sharma et al. 2013). All gene-specific primer sequences are provided in supplemental file S5 (supplementary table S20). Double-stranded RNA (dsRNA) was synthesized following the manufacturer’s protocol using a MEGAscript T7 kit (Ambion/Life Technologies) from amplified PCR product. The quality of dsRNA was assessed and concentrations adjusted using a NanoDrop ONE to 3.7-4 µg/µL. dsRNA was mixed with vital dyes for visualization of injections. Microinjection under halocarbon-700 oil (Sigma-Aldrich) was performed as previously described (Sharma et al. 2013). Subsets of developing embryos were fixed for HCR and assayed for selected genes.

### Imaging

Brightfield microscopy was performed using a Nikon SMZ fluorescence stereomicroscope mounted with a DSFi2 digital color camera driven by Nikon Elements software. SEM was performed using a Quanta FEI 200 scanning electron microscope. Confocal laser scanning microscopy was performed using a Zeiss LSM 780 microscope driven by Zen software. For *P. litorale*, confocal laser scanning microscopy was performed with a Leica SP5 microscope, driven by LAS-AF software. Beyond the documentation of gene expression (594nm and 633nm laser lines) and Dapi counterstain (405nm laser line), cuticular autofluorescence was separately recorded with the 488nm laser line. Using the software Amira 3D (version 2021.1; ThermoFisher Scientific), the cuticular signal in the 488nm channel was semi-automatically segmented (grey-value based thresholding) and the voxels included in the resulting material were set to grey value 0 in all other channels via the “Arithmetic” function, resulting in the separation of cuticular autofluorescence from gene expression signals.

## Supporting information

Electronic supplementary material

## Acknowledgements

Microscopy was performed at the Newcomb Imaging Center, Department of Botany, University of Wisconsin-Madison, with the aid of Sarah Swanson. The LABRE team and Gabriel Boaglio provided support during collecting trips in Argentina. Hugh G. Steiner and Pola O. Blaszczyk assisted with maintenance of *P. opilio.* Hypatia Coop in Madison (WI) kindly assisted with collection of pseudoscorpion embryos. Marion Wanninger and Max Hämmerle assisted with the maintenance of *P. litorale.* G.B. thanks Andreas Wanninger for providing research infrastructure. Discussions with Armin Moczek refined some of the experiments addressing *dac* subfunctionalization. This work was funded by the National Science Foundation grant no. IOS-2016141 grant to P.P.S; and National Institutes of Health NIAID grant nos. R21AI128393, R212200536 and R012200185 to M.G.-N.

## Author Contributions

B.C.K. and P.P.S. designed the objectives of the study. B.C.K., E.M.L., S.M.N., G.M.H., G.G., E.V.W.S., and P.P.S. performed functional experiments. B.C.K., E.M.L., S.M.N., G.G., and P.P.S. generated data for *P. opilio.* N.H.P. contributed tools and reagents for analyses, and sponsored the Whitman Fellowship for P.P.S. for arachnid tool development. G.B. generated data for *P. litorale.* A.A.B. and I.J. generated data for *A. longisetosus.* B.C.K. and I.A.H. generated data for *I. scapularis.* B.C.K., C.S., and D.E.V. field collected females and fixed resulting embryos of *T. salinarum.* B.C.K., G.G., E.V.W.S., and P.P.S. collected embryos and females of *P. parvus.* B.C.K. and P.P.S. performed RNA extraction and transcriptome assembly. B.C.K. generated expression data for *T. salinarum* and *P. parvus.* B.C.K., G.G., E.V.W.S., I.A.H., A.A.B., and G.B. performed bioinformatics. B.C.K., G.B., and P.P.S. wrote the first draft of the manuscript. B.C.K., M.S.H., G.B., and P.P.S. created figures and tables. P.P.S. and M.G.-N. procured funding.

## Data Availability

All image data are published in the manuscript or as supplemental material. All genomic resources are publicly available on NCBI.

