## Supplementary material for "A novel expression domain of *extradenticle* underlies the evolutionary developmental origin of the chelicerate patella": Electronic supplementary material

**This PDF file includes:**

Supplementary Figures S1 to S8
Supplementary Tables S1-S20
Supplementary Methods
Supplementary References

### Supplementary Figures

**Figure S1.** Wild type expression of proximodistal patterning genes in representative stages of appendage formation in *P. opilio.* All embryos in ventral view with anterior toward the right. Each row represents a single embryo with single-channel visualization of Hoechst nuclear counterstaining (cyan), and expression of *Po-dac* (yellow), *Po-exd* (magenta), and *Po-Dll* (red).

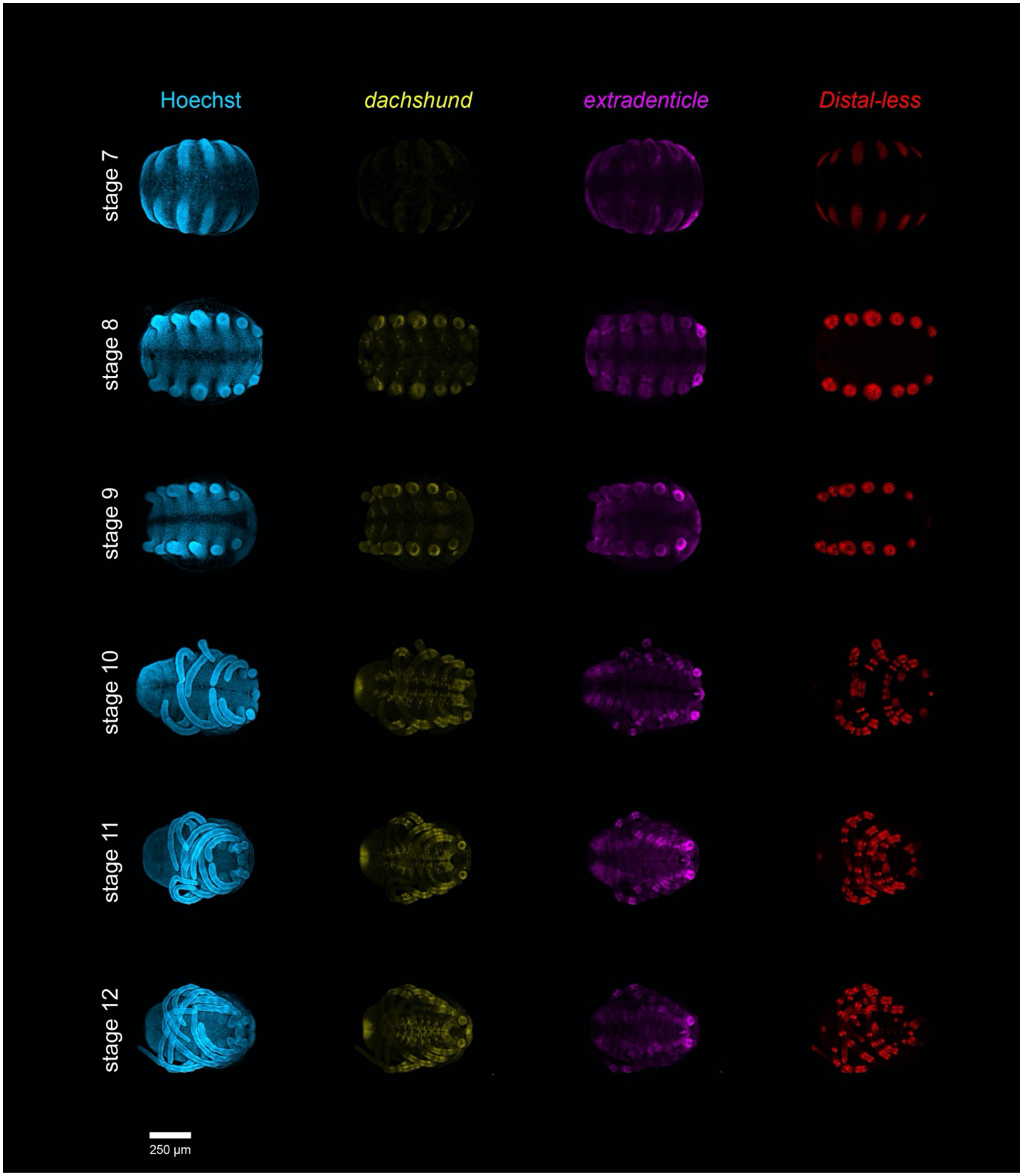

**Figure S2.** Wild type expression of proximodistal patterning genes in representative stages of appendage formation in *P. opilio.* All embryos in lateral view with anterior toward the right. Each row represents a single embryo with single-channel visualization of Hoechst nuclear counterstaining (cyan), and expression of *Po-dac* (yellow), *Po-exd* (magenta), and *Po-Dll* (red).

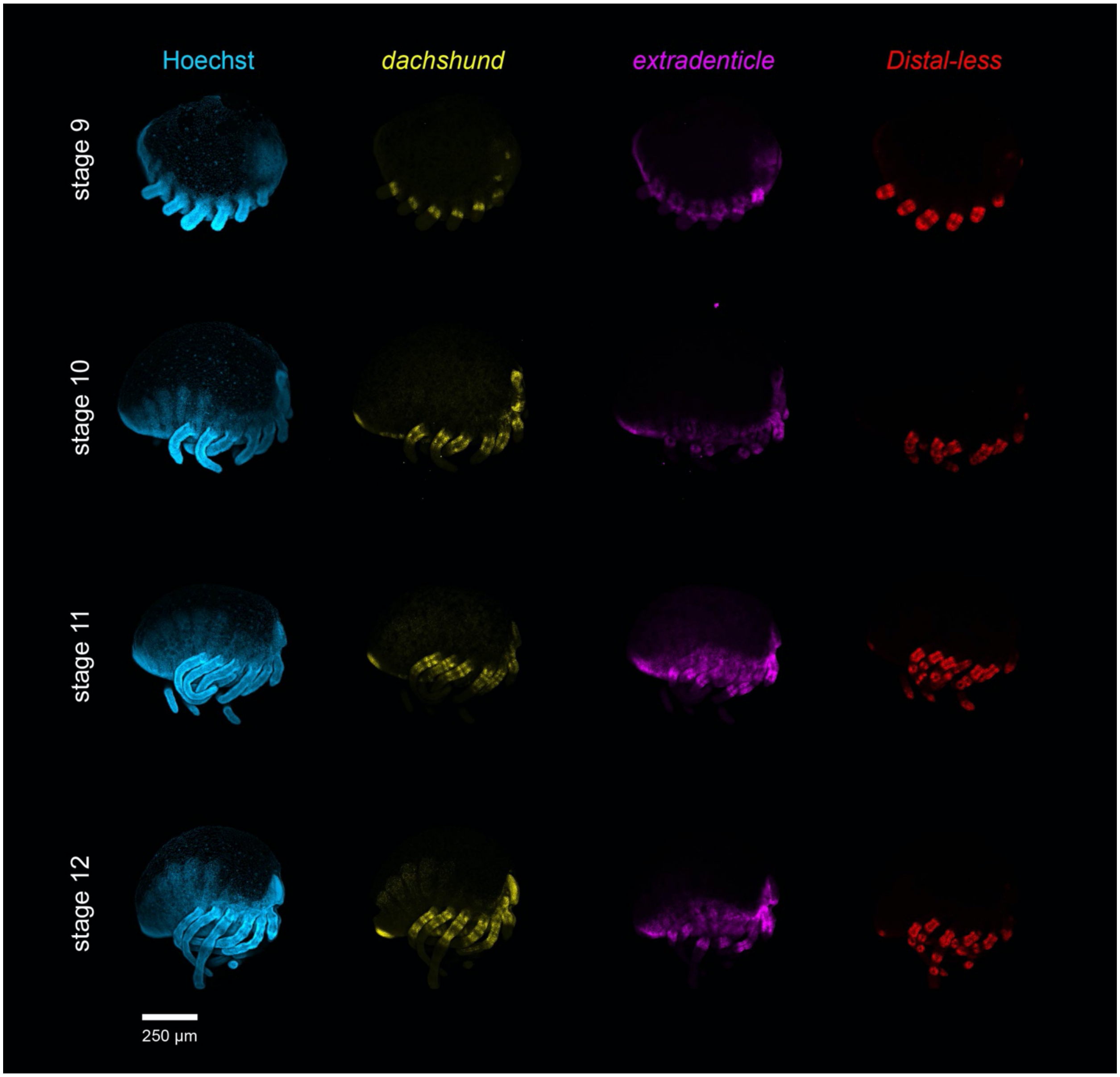

**Figure S3.** Wild type expression of proximodistal patterning genes in dissected appendages during representative stages of appendage formation in *P. opilio.* Appendages represent those in Figure 3 with demultiplexed single-channel visualization of Hoechst nuclear counterstaining (cyan), and expression of *Po-dac* (yellow), *Po-exd* (magenta), and *Po-Dll* (red). All appendages are in lateral view with distal to the right.

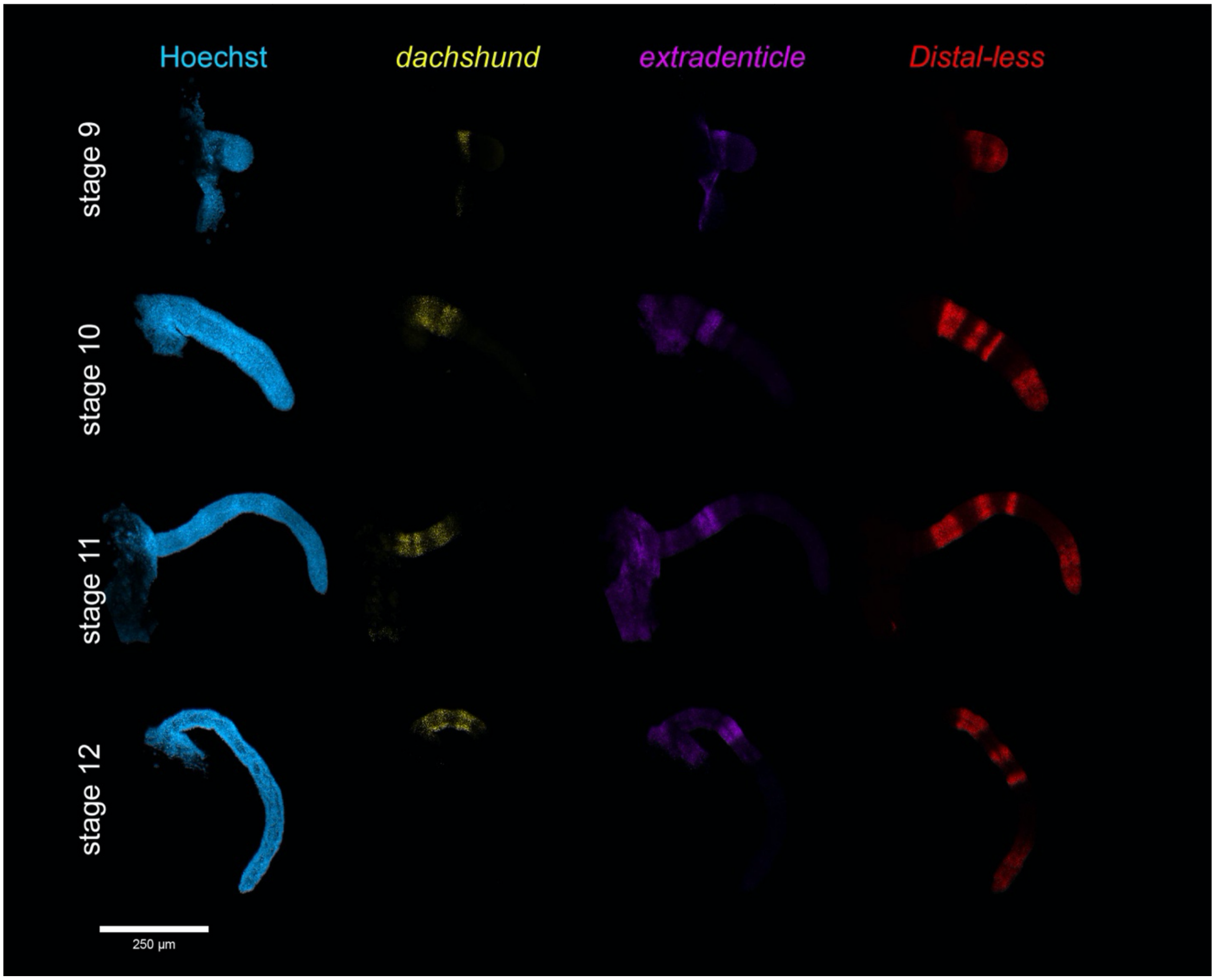

**Figure S4.** Validation of on-target *Po-exd* RNAi-mediated knockdown. Each row corresponds to an individual negative control (**a-a''**) or *Po-exd* RNAi embryo (**b-b''**). Embryos presented in ventral view with anterior toward the top. (**a, b**) Hoechst nuclear counterstaining (cyan). (**a', b'**) Expression of *Po-exd* (magenta). (**a'', b''**) Expression of *Po-Dll* (red).

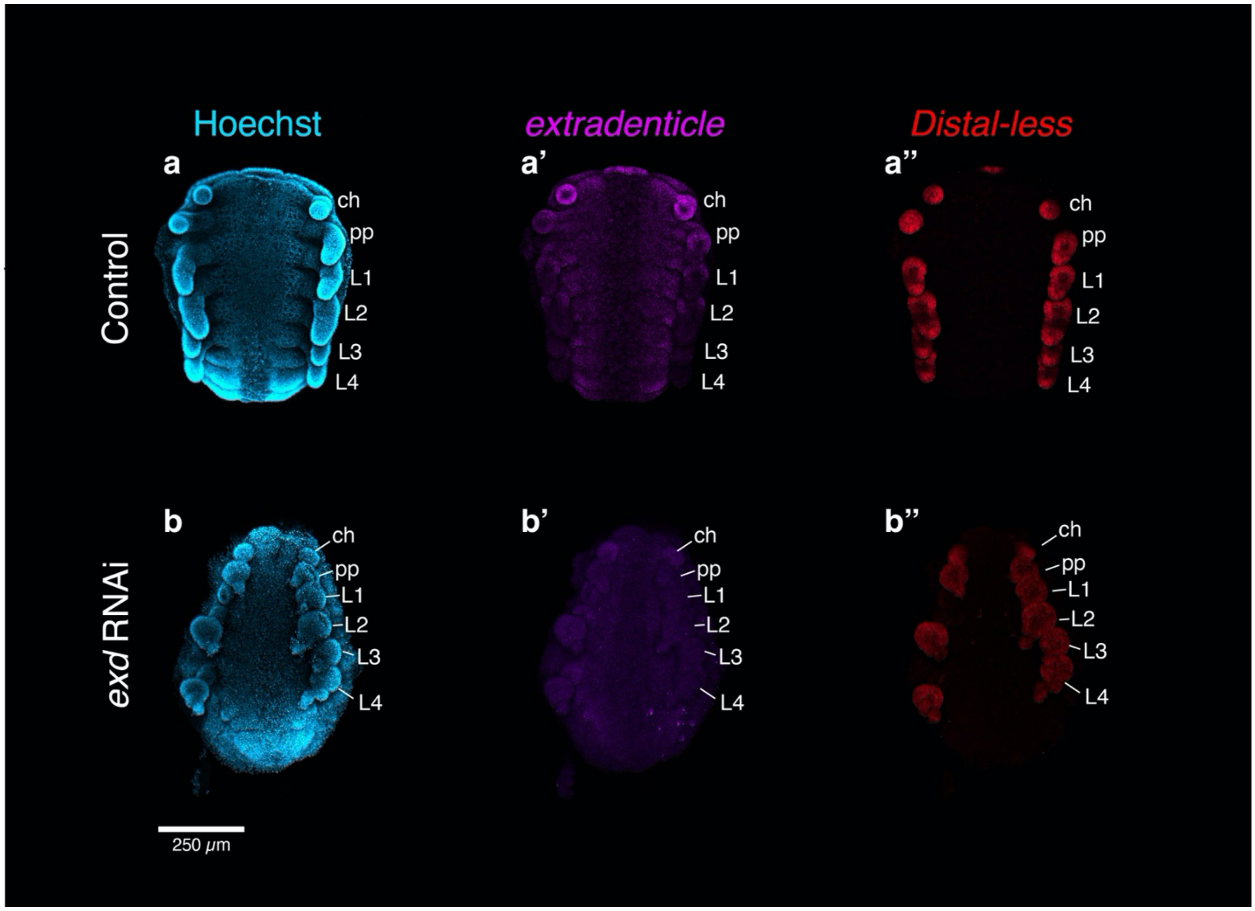

**Figure S5.** Knockdown of *Po-N* yields additional defects in central nervous system development, visualized via expression of *Po-dac.* Each row corresponds to an individual negative control (**a-a'''**) or *Po-N* RNAi embryo (**b-b'''**). Embryos presented in ventral view with anterior toward the top. (**A, B**) Hoechst nuclear counterstaining (cyan). (**a', b'**) Expression of *Po-dac* (yellow). (**a'', b''**) Expression of *Po-exd* (magenta). (**a''', b'''**) Expression of *Po-N* (orange). Abbreviations: lb, labrum; ch, chelicera; pp, pedipalp; L1, leg one; L2, leg two; L3, leg three; L4, leg four.

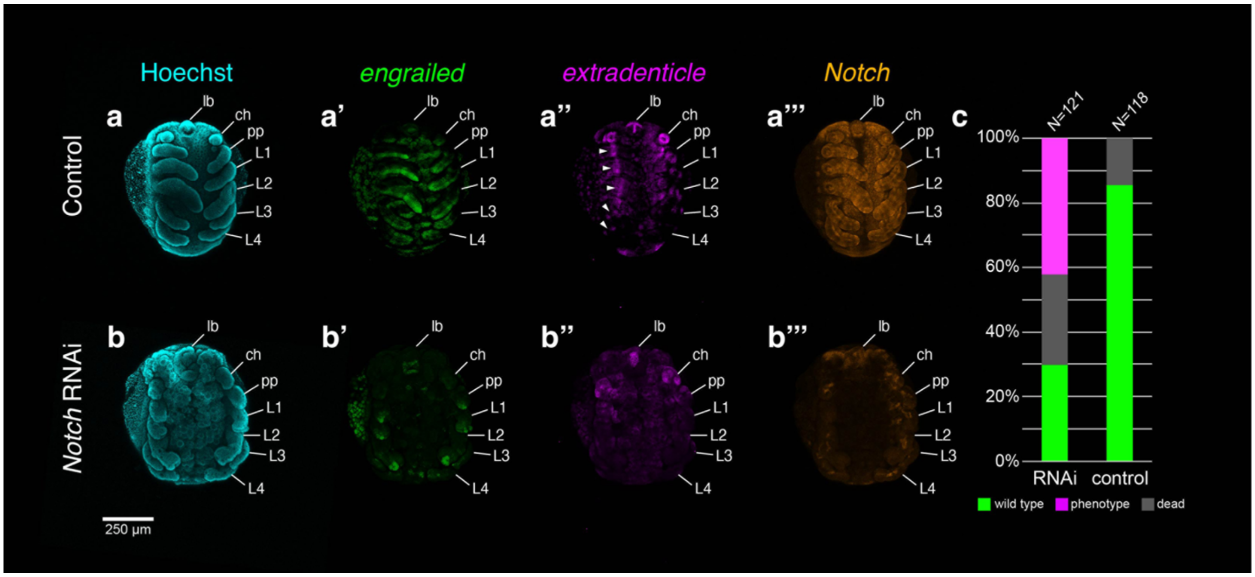

**Figure S6.** Range of observed segmental defects, visualized via expression of *Po-en* (green), following *Po-N* RNAi treatments. (**a**) Negative control embryo with wild type morphology and expression of *Po-en* in iterative stripes in the posterior compartment of all body segments and appendages. (**b**) Weak phenotype following *Po-N* RNAi. Embryo retains truncated, abnormal segmental stripes of *Po-en* in the body, but restricted expression in the appendages to distal termini. (**c**) Strong phenotype following *Po-N* RNAi. *Po-en* expression is detected only in the distal termini of appendages.

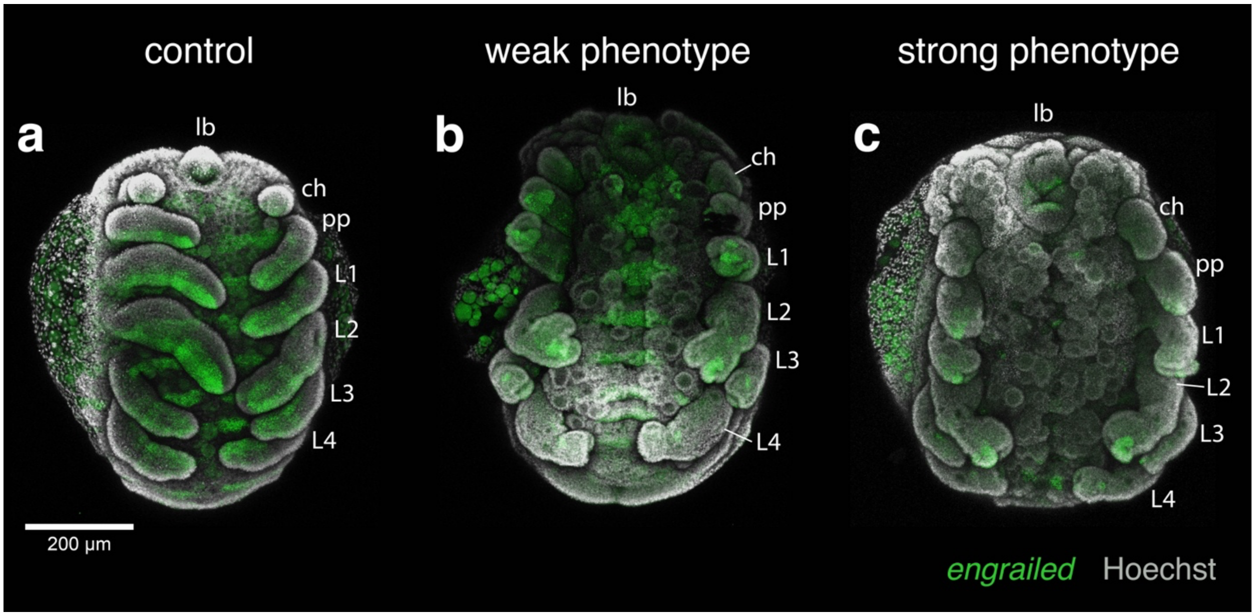

**Figure S7.** Dissected appendages of *Po-N* RNAi embryos demonstrate prominent reductions in *Po-exd* expression, but not *Po-dac.* Rows correspond to individual appendages from negative control (**a-a'''**) or *Po-N* RNAi embryos (**b-b''', c-c''**). All appendages in lateral view with distal to the right. (**a-c**) Hoechst nuclear counterstaining (gray) with merged expression of *Po-dac* (yellow) and *Po-exd* (magenta). (**a'-c'**) Single-channel visualization of Hoechst counterstaining. (**a''-b''**) Single-channel expression of *Po-dac.* (**a'''-c''**) Single-channel expression of *Po-exd.* Embryo in (**c-c''**) was not surveyed for expression of *Po-dac.*

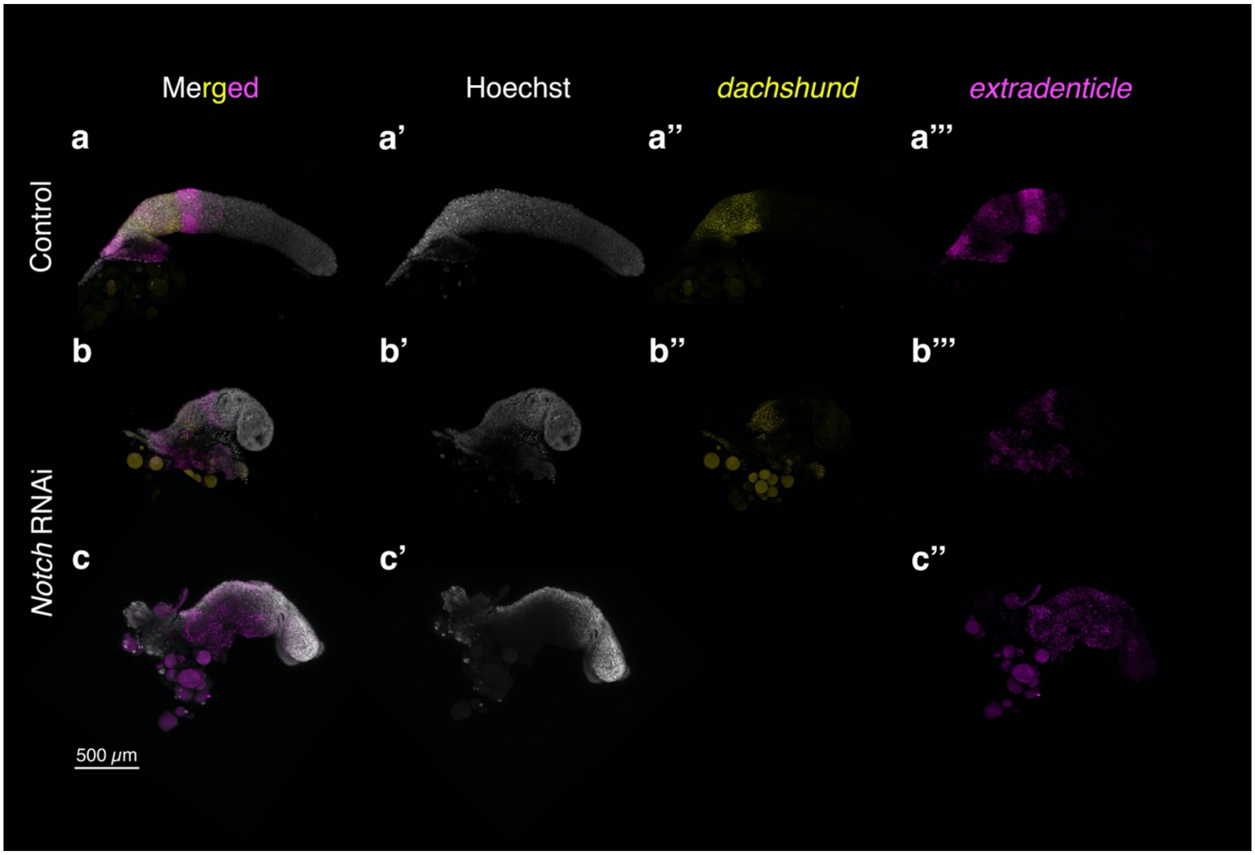

**Figure S8.** Overview of *Pl-exd, Pl-dac,* and *Pl-Dll* expression during postembryonic development of *P. litorale.* All images show ventral views. One leg of instar IV (**d, e**) has been virtually removed (dashed line). Arrows: distal boundary of *Pl-dac* domain. Arrowheads: *Pl-exd* expression distal to the *Pl-dac* domain. Asterisks indicate regions of damaged tissue. (**a**) Late instar II. (**b**) Early instar III. (**c**) Late instar III. (**d**) Early instar IV. (**e**) Late instar IV. Abbreviations: ch, chelicera; L, leg; ov, ovigeral larval limb; pp, pedipalpal larval limb.

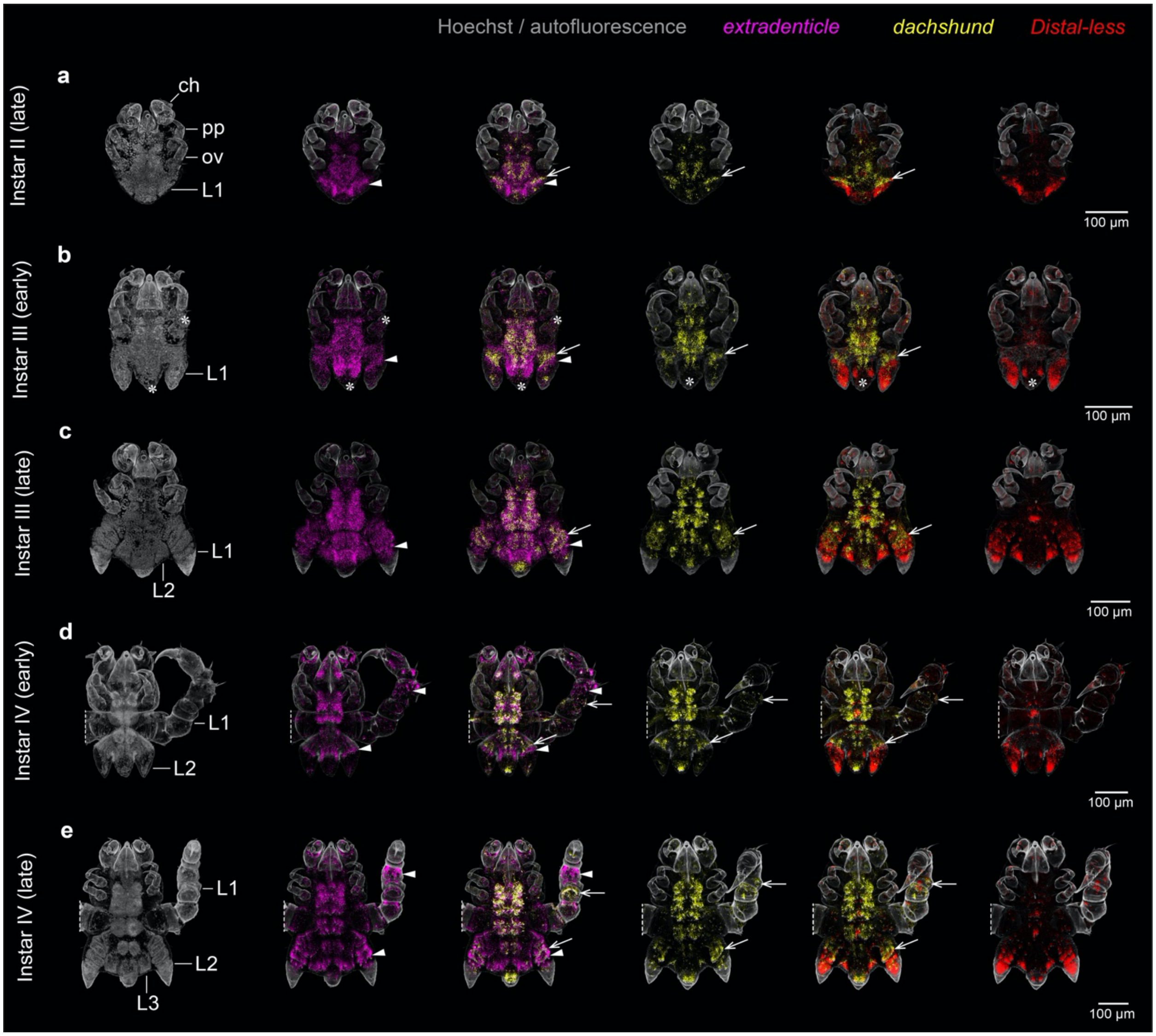

**Table S1.** Select alternative terminologies for pedipalp podomeres across chelicerate orders. Abbreviations: Cx – coxa; Tr – trochanter; Fe – femur; Tita – tibiotarsus; Pa – patella; Ti – tibia; Ta – tarsus; Mt – metatarsus; Bta – basitarsus; Dta – distitarsus; Gn – genu.

| Order | P-I | P-II | P-III | P-IV | P-V | P-VI | Terminus | Reference |
|---|---|---|---|---|---|---|---|---|
| Scorpiones | Cx | Tr | Pre-Fe | Fe | Tita | Movable Finger (Chela) | —— | Millot & Vachon (1949a) |
| Scorpiones | Cx | Tr | Fe | Pa | Ti | Ta | —— | Hjelle (1990) |
| Pseudoscorpiones | Cx | Tr | Fe | Ti | Chela | Movable Finger | —— | Chamberlin (1931) |
| Pseudoscorpiones | Cx | Tr | Fe | Ti | Mt | Ta | —— | Savory (1964) |
| Pseudoscorpiones | Cx | Tr | Pre-Fe | Fe | Tita | Pretarsus | —— | Weygoldt (1969) |
| Pseudoscorpiones | Cx | Tr | Fe | Pa | Ti (Chela) | Ta (Movable Finger) | —— | Shultz (1989); Harvey (1992) |
| Araneae | Cx | Tr | Fe | Pa | Ti | Ta | Claws | Foelix (2011) |
| Amblypygi | Cx | Tr | Fe | Ti | Bta | Dta | Claws | Millot (1949a); Weygoldt (2000) |
| Amblypygi | Cx | Tr | Fe | Pa | Ti | Ta | Post-Tarsus | Savory (1964) |
| Uropygi | Cx | Tr | Fe | Ti | Bta | Ta | —— | Graveley (1915); Savory (1964); Millot (1949d) |
| Uropygi | Cx | Tr | Fe | Pa | Ti | Ta | Apotele | Shultz (1993) |
| Schizomida | Cx | Tr | Fe | Ti | Bta | Ta | Epitarsus | Millot (1949d); Savory (1964) |
| Schizomida | Cx | Tr | Fe | Pa | Ti | Ta | Spur | Rowland (1975) |
| Ricinulei | Cx | Tr 1 | Tr 2 | Fe | Ti | Ta | Spur (Éperon) | Millot (1949c); Pittard & Mitchell (1972) |
| Palpigradi | Cx | Tr | Fe | Ti | Bta | Ta | Pseudonychium | Millot (1949b) |
| Palpigradi | Cx | Tr | Fe | Ti | Mt | Ta | —— | Börner (1901) |
| Palpigradi | Cx | Tr | Fe | Gn | Ti | Ta | Apotele | van der Hammen (1982) |
| Solifugae | Cx | Tr | Fe | Ti | Bta | Ta | Adhesive Organ | Millot & Vachon (1949b) |
| Solifugae | Cx | Tr | Fe | Ti | Mt | Ta | Adhesive Organ | Punzo (1998) |
| Opiliones | Cx | Tr | Fe | Pa | Ti | Ta | Claws | Shultz & Pinto-da-Rocha (2007) |
| Parasitiformes | Cx | Tr | Fe | Gn | Ti | Ta | Apotele | van der Hammen (1966); |

| | | | | | | | | |
|---|---|---|---|---|---|---|---|---|
| | | | | | | | | Krantz & Walter (2009) |

**Table S2.** Select alternative terminologies for walking leg podomeres across chelicerate orders. Abbreviations: Cx – coxa; Fe – femur; Ti – tibia; Ta – tarsus; Pr – propodus; Tr – trochanter; Pa – patella; Tita – tibiotarsus; Bta – basitarsus; Tta – telotarsus; Bfe – basifemur; Tfe – telofemur; Mt – metatarsus; Gn – genu; Bti – basitibia; Tti – telotibia; Ata – acrotarsus; Btr – basitrochanter; Ttr – telotrochanter.

| Order | P-I | P-II | P-III | P-IV | P-V | P-VI | P-VII | P-VIII | P-IX | Terminus | Reference |
|---|---|---|---|---|---|---|---|---|---|---|---|
| Pantopoda (Pycnogonida) | Cx 1 | Cx 2 | Cx 3 | Fe | Ti 1 | Ti 2 | Ta | Pr | —— | Main Claw | Sars (1891); Helfer & Schlottke; King (1973) |
| Pantopoda | Cx 1 | Cx 2 | Cx 3 | Fe | Ti 1 | Ti 2 | Ta 1 | Ta 2 | —— | Main Claw | Meinert (1899) |
| Pantopoda | Cx | Tr 1 | Tr 2 | Fe | Pa | Ti | Ta 1 | Ta 2 | —— | Pretarsus | Snodgrass (1958); Schram & Hedgpeth (1978) |
| Xiphosura (L1-4) | Cx | Tr | Fe | Pa | Tita | Pre-tarsus (movable finger) | —— | —— | —— | —— | van der Hammen (1989); Shultz (1989) |
| Xiphosura (L5) | Cx | Tr | Fe | Pa | Ti | Ta | Pre-tarsus | —— | —— | —— | Shultz (1989) |
| Scorpiones | Cx | Tr | Fe | Ti | Ta 1 | Ta 2 | Ta 3 | —— | —— | —— | Kraepelin (1891) |
| Scorpiones | Cx | Trans-Cx | Pre-Fe | Fe | Ti | Bta | Ta | —— | —— | —— | Millot & Vachon (1949a) |
| Scorpiones | Cx | Tr | Fe | Pa | Ti | Bta | Tta | —— | —— | —— | Hjelle (1990) |
| Pseudoscorpiones | Cx | Tr | Bfe | Tfe | Ti | Mt | Ta | —— | —— | Arolium & Claws | Chamberlin (1931) |
| Pseudoscorpiones | Cx | Tr | Bfe | Tfe | Ti | Mt | Ta | —— | —— | Pretarsus | Savory (1964) |
| Pseudoscorpiones | Cx | Tr | Fe | Pa | Ti | Mt | Ta | —— | —— | Arolium & Claws | Shultz (1989); Harvey (1992) |
| Pseudoscorpiones (Monosphyronida) | Cx | Tr | Fe | Pa | Ti | Mio-Ta | —— | —— | —— | Arolium & Claws | Chamberlin (1931) |
| Pseudoscorpiones (Feaelloidea & Elassommatina) | Cx | Tr | Fe | Pa | Ti | Ta | —— | —— | —— | Arolium & Claws | Harvey (1992) |
| Araneae | Cx | Tr | Fe | Pa | Ti | Mt | Ta | —— | —— | —— | Foelix (2011) |
| Amblypygi | Cx | Tr | Fe | Pa | Ti | Bta | Tta | —— | —— | Apotele | Millot (1949a); Weygoldt (2000) |
| Uropygi | Cx | Tr | Fe | Pa | Ti | Bta | Tta | —— | —— | —— | Millot (1949d) |
| Uropygi | Cx | Tr | Fe | Pa | Ti | Mt | Ta | —— | —— | —— | Savory (1964) |
| Schizomida | Cx | Tr | Fe | Pa | Ti | Bta | Ta | —— | —— | —— | Millot (1949d); Rowland (1975) |
| Schizomida | Cx | Tr | Fe | Pa | Ti | Mt | Ta | —— | —— | —— | Savory (1964) |
| Ricinulei (L1-2) | Cx | Tr | Fe | Pa | Ti | Bta | Ta | —— | —— | Claws | Millot (1949c) |

| | | | | | | | | | | | |
|---|---|---|---|---|---|---|---|---|---|---|---|
| Ricinulei (L1-2) | Cx | Tr | Fe | Pa | Ti | Mt | Ta | —— | —— | Claws | Pittard & Mitchell (1972) |
| Ricinulei (L3-4) | Cx | Tr 1 | Tr 2 | Fe | Pa | Bta | Ta | —— | —— | Claws | Millot (1949c) |
| Ricinulei (L3-4) | Cx | Tr 1 | Tr 2 | Fe | Pa | Mt | Ta | —— | —— | Claws | Pittard & Mitchell (1972) |
| Palpigradi | Cx | Tr | Fe | Pa | Ti | Mt | Ta | | —— | —— | Börner (1901) |
| Palpigradi | Cx | Tr | Fe | Pa | Ti | Bta | Ta | —— | —— | Pretarsus | Shultz (1989) |
| Palpigradi | Cx | Tr | Fe 1 | Fe 2 | Gn | Ti | Ta | —— | —— | Apotele | van der Hammen (1982) |
| Solifugae (L1-2) | Cx | Tr | Pre-Fe | Post-Fe | Ti | Bta | Ta | —— | —— | —— | Millot & Vachon (1949b); Punzo (1998) |
| Solifugae (L1-L2) | Cx | Tr | Fe | Pa | Ti | Bta | Ta | —— | —— | —— | Shultz (1989) |
| Solifugae (L3-4) | Cx | Tr 1 | Tr 2 | Pre-Fe | Post-Fe | Ti | Bta | Ta | —— | —— | Millot & Vachon (1949b); Punzo (1998) |
| Solifugae (L3-4) | Cx | Tr | Bfe | Tfe | Pa | Ti | Bta | Ta | —— | —— | Shultz (1989) |
| Opiliones | Cx | Tr | Fe | Pa | Ti | Mt | Ta | —— | —— | Claws | Shultz & Pinto-da-Rocha (2007) |
| Parasitiformes (Opilioacarida, L1) | Cx | Tr | Bfe | Tfe | Gn | Bti | Tti | Bta | Tta | Apotele | van der Hammen (1966) |
| Parasitiformes (Opilioacarida, L2) | Cx | Tr | Fe | Gn | Ti | Bta | Tta | Ata | —— | Apotele | van der Hammen (1966) |
| Parasitiformes (Opilioacarida, L3-L4) | Cx | Tr 1 | Tr 2 | Fe | Gn | Ti | Bta | Tta | Ata | Apotele | van der Hammen (1966) |
| Parasitiformes (Opilioacarida, L1) | Cx | Tr | Bfe | Tfe | Gn | Ti | Bta | Tta | —— | —— | Krantz & Walter (2009) |
| Parasitiformes (Opilioacarida, L2) | Cx | Tr | Fe | Gn | Ti | Bta | Tta | —— | —— | —— | Krantz & Walter (2009) |
| Parasitiformes (Opilioacarida, L3-4) | Cx | Btr | Ttr | Fe | Gn | Ti | Bta | Tta | —— | —— | Krantz & Walter (2009) |
| Parasitiformes (Holothyrida) | Cx | Tr | Bfe | Tfe | Gn | Ti | Ta | —— | —— | Ambulacrum & Claws | Krantz & Walter (2009) |
| Parasitiformes (Ixodida) | Cx | Tr | Bfe | Tfe | Gn | Ti | Bta | Tta | —— | Claws | Krantz & Walter (2009) |
| Parasitiformes (Mesostigmata) | Cx | Tr | Bfe | Tfe | Gn | Ti | Bta | Tta | —— | —— | Krantz & Walter (2009) |
| Acariformes (Trombidiformes) | Cx | L-1 | L-2 | L-3 | L-4 | L-5 | L-6 | —— | —— | —— | Cook (1974) |
| Acariformes (Trombidiformes, Endeostigmata, Parasitengona) | Cx | Tr | Bfe | Tfe | Gn | Ti | Ta | —— | —— | Apotele | Evans (1992) |
| Acariformes (Oribatida) | Cx | Tr | Fe | Gn | Ti | Ta | —— | —— | —— | Apotele | Evans (1992); Krantz & Walter (2009) |

### Supplementary Methods

### Hybridization Chain Reaction Probe Design

Probes for *Archegozetes longisetosus exd* (17 probe pairs) and *Pycnogonum litorale Dll* (19 probe pairs) were designed by Molecular Instruments using the following CDS templates:

>*Alon_exd*
ATGGACGATAATCAACACAATCATTCACTATCAATAATGCATGCGGTCTCTCAACAAATGTCGGGTCATGTAGTGC
CACAACCGCCTCACGGATACGGGCTGTCCGCTGGTCACGACCCGACCGGTACCACCAATTCCGATGGAGAAACAC
GAAAACATGACATCTCAGAGATATTACAGCAAATCATGAATATTACTGACCAAAGTCTCGATGAGGCTCAGGCCA
GGTCTGTATTGACAACAAATAACAGGAAACATACTTTGAATTGTCATCGCATGAAGCCAGCACTATTCAGCGTTTT
ATGTGAAATCAAGGAAAAGACAGTATTGAGCTTAAGAAACACTCAAGAAGAAGAACCACCAGATCCACAACTGA
TGCGGTTAGACAATATGCTAATAGCGGAAGGGGTTGCAGGACCCGAAAAAGGTGGCGGTCAGTCTGCGGCCGCTA
ATGCGGCGGCAGCTGCGGCTGCGGGTCAGTCGGGAGGTGAAAATGCAATCGAGCACTCAGACTATAGGGCCAAAC
TCGCACAAATTAGAACAATATACCATCAAGAACTTGAGAAATATGAACAGGCCTGTAACGAGTTCACAACACATG
TTATGAATCTATTACGAGAGCAAAGTCGGACGCGTCCAATTACGCCAAAAGAAATCGAAAGAATGGTTCAAATTA
TACACAAAAAGTTTAATTCAATTCAAGTACAGCTCAAACAAAGCACCTGTGAAGCAGTTATGATTCTTAGATCAAG
GTTTTTGGATGCCAGGAGAAAACGAAGAAATTTCAGCAAGCAAGCAACTGAAATATTAAATGAATATTTTTATTCA
CACCTGAGTAACCCATACCCGAGTGAAGAAGCAAAAGAAGAATTAGCGCGAAAGTGTAGTATTACAGTATCTCAA
GTATCAAATTGGTTCGGAAATAAGAGAATTAGGTATAAGAAAAATATTGGAAAAGCTCAAGAAGAAGCAAATCTT
TATGCAGCTAAAAAAGCAGCCGGTTCGTCCCCATACGGTTTAACGCCATCATCTCAGGGCTCGGGAAATATAATGA
GTCCACCGCCACCACCTGGTAGTTCACAGGATGGATGGTCTATGAATGGTGACTATGGAAGTCAGGCCCATAGGC
ACGGAGGCTATTCCTCAGATGGTTCCTCGGGAGGCATGTATGATCCCGGGATGCATCAGGTATCACCTCTTTGA

>*Plit_Dll*
AGTTTCAAAAAATTATTTTTTATATGTTTACGGTGAAGCGTTGAATTTTGTTTTTTTTTAATTTAAAAAAAAAAATCTT
ATTTTATTATTTGACTTTGTTCGGTGTAAACTAACGGCCGTCACCACGTTGAATTTTACGGATTTGTCGTGTCCATA
CCCGATAGCCACTAGTATGGCGGGTAATTCGGATCACGGTTCAATCGAACAAGAACATTTGTCCAAACAGTCCGCA
TTCATGGAGATTCAACAAGCTGGCGGACTGGGACCTCATCACGGGACTACGACGTATCCTATCAGGTCAAGTTATC
AGAGTCAACCGCCTCAACATCACGATAGTGCGTTCGCCATGTCTCATGCTCAACAAGCTGCGGCGGCGGCCGCCG
GTCGTTCAGCCTTAGGACCGTATCATTTCCCGATGAATGCGCATAATACTCCCGCTGGTTATCCAACTGCTCATCCT
TATCTCGGAGCGTATCAGACATCGTGTCCTTCACCTCCAAGAGACGATAAATCGCACATGGAAGAAACATTACGA
GTAAATGGAAAAGGTAAAAAAATGCGTAAACCTAGAACAATATACTCGAGTTTACAGCTACAACAGTTGAATAGA
AGATTTCAAAGAACCCAGTATTTAGCTTTACCCGAAAGAGCCGAATTAGCAGCTTCATTAGGACTTACACAAACAC
AGGTAAAAATATGGTTTCAAAATAGACGGTCGAAATACAAAAAGCTGATGAAGAGTAATCCAACACCTGGTGGAC
CGATTCCACACTTAGGTGGACCGCCATCGCAGCACAGTGTCAATAACGGACCTCTCCAGCATCCGAATCCGGCTAC
AACTCCACAAACTCCACCGTCGCAACAAAATCATCCGTCACATAGTCCACATAGTGATCATCATCATCATCATCCA
TTAGGTAATAACGGTCCTCACGGGCTTGGATTGTTACCTGTTTCGTGCGGACCCGGTATAAGTCCGTCTTCGATAAG
TTCTCCGATGGACTCTTGGAGTATGAACAGTGTTGCCGCCAAAGCCGCTGCTGCAGCCGCGGCAGTTAACGTCACC
AATTCTTACATGAGTCAATATCCTTGGTATCATCAGCCTGATCCGAGTATTAATCAACAATTATTAACGTAG

**Table S3.** Probe pairs designed for *Phalangium opilio exd* HCR in situ hybridization (B2 initiator).

| Pair | Initiator | Spacer | Hybridization | Hybridization | Spacer | Initiator |
|---|---|---|---|---|---|---|
| 1 | CCTCGTAAATCC TCATCA | AA | TATACCCGGGTCGTACATT CCTGCC | TTACGGATGCAATTCCGGT GGCTGA | AA | ATCATCCAGTAA ACCGCC |
| 2 | CCTCGTAAATCC TCATCA | AA | TCCTCCATCTGAGAAATGA CGTGCC | GGCTGAGGCGGCATCCCG TCAGAAT | AA | ATCATCCAGTAA ACCGCC |
| 3 | CCTCGTAAATCC TCATCA | AA | CGGCGTTGTCCATAGTGGA CGTAAC | GCGCATTCCCTTGCGTTTG GACAAG | AA | ATCATCCAGTAA ACCGCC |
| 4 | CCTCGTAAATCC TCATCA | AA | GACTTGACCATTCGAAATG GCGAAC | AGGAACGCTCGTGTGATA GTTATCG | AA | ATCATCCAGTAA ACCGCC |
| 5 | CCTCGTAAATCC TCATCA | AA | GAACTGACCATTGGTGGA CACGGAT | CTGCCATCCGTGGATACCG TTGGAG | AA | ATCATCCAGTAA ACCGCC |
| 6 | CCTCGTAAATCC TCATCA | AA | TCGTTAGACCAGCCGAAG CGGCTTT | GCGTCGTGGGAAGGCTGT ACTCTGG | AA | ATCATCCAGTAA ACCGCC |

| | | | | | | |
|---|---|---|---|---|---|---|
| 7 | CCTCGTAAATCCTCATCA | AA | AGCTTTACCAATATTTTTCTTGTAT | GGCCGCGTATAAATTGGCTTCTTCC | AA | ATCATCCAGTAAACCGCC |
| 8 | CCTCGTAAATCCTCATCA | AA | ACCTGGGAGACTGTTATGCCACACT | ATACGTTTGTTACCGAACCAATTAG | AA | ATCATCCAGTAAACCGCC |
| 9 | CCTCGTAAATCCTCATCA | AA | TAGGGTAAGGGTTACTTAAATGTGA | TGGCTAATTCCTCCTTGGCTTCTTC | AA | ATCATCCAGTAAACCGCC |
| 10 | CCTCGTAAATCCTCATCA | AA | TTGCTTGCTAAAGTTCCGCCTTTTC | GAAGTACTCGTTGAGAATTTCCGTC | AA | ATCATCCAGTAAACCGCC |
| 11 | CCTCGTAAATCCTCATCA | AA | ATCATGACGGCTTCGCACGTGCTCT | CTCGCGTCTAGAAACCTGGATCGCA | AA | ATCATCCAGTAAACCGCC |
| 12 | CCTCGTAAATCCTCATCA | AA | TCTTGTGGATGATCTGAACCATTCG | TGAGCTGCACTTGGATGGAGTTGAA | AA | ATCATCCAGTAAACCGCC |
| 13 | CCTCGTAAATCCTCATCA | AA | CCTGCTCTGTTCCCGGAGCAGGTTC | GATCTCCTTGGGCGTGATGGGACGC | AA | ATCATCCAGTAAACCGCC |
| 14 | CCTCGTAAATCCTCATCA | AA | TGTTCGTATTTTTCGAGCTCCTGGT | ACGTGCGTCGTGAACTCGTTGCACG | AA | ATCATCCAGTAAACCGCC |
| 15 | CCTCGTAAATCCTCATCA | AA | CTCTGTAGTCGGAGTGTTCAATGGC | AGATTTGCCTAATCTGCGCCAGTTT | AA | ATCATCCAGTAAACCGCC |
| 16 | CCTCGTAAATCCTCATCA | AA | TGCCGCCGAAGCGTTGGCCGCCGCG | TTCGGGTTGACCGGGCCCGCCCGCC | AA | ATCATCCAGTAAACCGCC |
| 17 | CCTCGTAAATCCTCATCA | AA | ACTCCTTCCGCTATTAACATATTGT | GGTCCGCTTCCCTTTTCTGGACCGG | AA | ATCATCCAGTAAACCGCC |
| 18 | CCTCGTAAATCCTCATCA | AA | CTTCTTCTTGTGTGTTTCTCAAACT | GCCTCATTAACTGAGGGTCTGGCGG | AA | ATCATCCAGTAAACCGCC |

**Table S4.** Probe pairs designed for *Phalangium opilio dac* HCR in situ hybridization (B1 initiator).

| Pair | Initiator | Spacer | Hybridization | Hybridization | Spacer | Initiator |
|---|---|---|---|---|---|---|
| 1 | GAGGAGGGCAGCAAACGG | AA | TGGGTTGTCGCTTAAGTCGCCTTGG | CTCGTTATTCGCCGTACTCGTTACG | TA | GAAGAGTCTTCCTTTACG |
| 2 | GAGGAGGGCAGCAAACGG | AA | TCATTGTCCTCGTCATCAGCGCACT | CGTTCATCCTCTTCTTCATCTTCCG | TA | GAAGAGTCTTCCTTTACG |
| 3 | GAGGAGGGCAGCAAACGG | AA | CGCCCGTGGCGTTACCATTTCCGGC | TGTAATTGTGTTCGCTGCCGCTGTG | TA | GAAGAGTCTTCCTTTACG |
| 4 | GAGGAGGGCAGCAAACGG | AA | TCGGGTACTGTTCTGCGATAGGTTA | CAAGCCATTACCCGGACCGCCAACG | TA | GAAGAGTCTTCCTTTACG |
| 5 | GAGGAGGGCAGCAAACGG | AA | CCTGCAAGAAGCCAGAAGTAGCCAT | ACCGGACTGTGACCGTTGGCCGAGG | TA | GAAGAGTCTTCCTTTACG |
| 6 | GAGGAGGGCAGCAAACGG | AA | TGTCAAAACCGTAAAATTTATCCTT | AGAACGTGGGGTCCTTCAGTCGGTG | TA | GAAGAGTCTTCCTTTACG |
| 7 | GAGGAGGGCAGCAAACGG | AA | GACCCCGTCCGTATGTCCTTGTGAC | GTCCGCTAGCTCTTCTCTTGATGAG | TA | GAAGAGTCTTCCTTTACG |
| 8 | GAGGAGGGCAGCAAACGG | AA | GAGGATGGGTTTATGTTGTTGGAAC | TCTTTAATGACTGAACCGTCCGTCC | TA | GAAGAGTCTTCCTTTACG |
| 9 | GAGGAGGGCAGCAAACGG | AA | AGACGAGGAAGATGAGGATGATGAT | AGCCGCCGCCTGGGCGGCCGCCGAT | TA | GAAGAGTCTTCCTTTACG |
| 10 | GAGGAGGGCAGCAAACGG | AA | CATGAAAGGCAGCGGGTTGATGTGA | CCCGTGGGCGTTGGGGTGGTTCAGC | TA | GAAGAGTCTTCCTTTACG |
| 11 | GAGGAGGGCAGCAAACGG | AA | AGCCGTTCGGTAACAAACCGTGCTT | CCGCTTGGGCCGCCGCGTGGTGACT | TA | GAAGAGTCTTCCTTTACG |
| 12 | GAGGAGGGCAGCAAACGG | AA | TCGGCTTTTGGAAATGTGTCCGTTT | CAATCGTTCACCTGTTGAATAGAAA | TA | GAAGAGTCTTCCTTTACG |
| 13 | GAGGAGGGCAGCAAACGG | AA | CCTTTCTTGAGAAGGCCGGCGTGTG | TATCCACCGTACTCTCCGTCAATTC | TA | GAAGAGTCTTCCTTTACG |
| 14 | GAGGAGGGCAGCAAACGG | AA | ACGCTCTCTTTGGTGGTCGGCCCGG | TGGAGGAACTGGTCAGTCCCACCAT | TA | GAAGAGTCTTCCTTTACG |
| 15 | GAGGAGGGCAGCAAACGG | AA | TGTGCAATCTTTATACAAGGAGTCG | GCCAGAAGAGAAGACACACCTCGCT | TA | GAAGAGTCTTCCTTTACG |
| 16 | GAGGAGGGCAGCAAACGG | AA | TTGACTCCGGGTTGTATGGCTCCTA | TCTTTGCAGCTGAGTAATTTGCAAC | TA | GAAGAGTCTTCCTTTACG |
| 17 | GAGGAGGGCAGCAAACGG | AA | TACAAACGATGGGAGTTATGTCCAA | CTCTGAGAATTCGAACTTGTTCGAC | TA | GAAGAGTCTTCCTTTACG |
| 18 | GAGGAGGGCAGCAAACGG | AA | TCCTCCGACCAGGTGTTTCAAAAAC | TTTGAGTTTAGTGTAGACCGTGTGC | TA | GAAGAGTCTTCCTTTACG |
| 19 | GAGGAGGGCAGCAAACGG | AA | TATTCTCCACCGACCAAGAAAGCGG | TCGAAAGCTTGGGGTAGACAAAGTA | TA | GAAGAGTCTTCCTTTACG |
| 20 | GAGGAGGGCAGCAAACGG | AA | GGCATTCGTTGCCGACGCCGTCACC | CTTTAGCTCCGCGATAATCTATCAA | TA | GAAGAGTCTTCCTTTACG |

**Table S5.** Probe pairs designed for *Phalangium opilio Notch* HCR in situ hybridization (B3 initiator).

| Pair | Initiator | Spacer | Hybridization | Hybridization | Spacer | Initiator |
|---|---|---|---|---|---|---|
| 1 | GTCCCTGCCTCTATATCT | TT | ATGGGCGTTTTGAGATTGCTGATTT | TCAAATGAAAACCGCTTCTTGGGGT | TT | CCACTCAACTTTAACCCG |
| 2 | GTCCCTGCCTCTATATCT | TT | GTCCGATTGAGCGGAATGGGGTGAA | CAAAGGGCTAGAGATTCCTTCCGAC | TT | CCACTCAACTTTAACCCG |
| 3 | GTCCCTGCCTCTATATCT | TT | AACCACCGTGGCTATGTTGGGAAGG | GTAAATAATGTTGAGGGGTAGCTTC | TT | CCACTCAACTTTAACCCG |
| 4 | GTCCCTGCCTCTATATCT | TT | CCACCACTCAAGAGCATTTGAAGGT | TGTCCACCCATATTGTTTCCGCCAC | TT | CCACTCAACTTTAACCCG |
| 5 | GTCCCTGCCTCTATATCT | TT | GAAGTTGGCAATGATGGTCGTTGTT | GCCCTCATAACGGCCATGTGAGTGG | TT | CCACTCAACTTTAACCCG |
| 6 | GTCCCTGCCTCTATATCT | TT | GTTTTGACGAGGGTGATGCGCTTTC | TCCGTTACTCATACCACCAGGCATT | TT | CCACTCAACTTTAACCCG |
| 7 | GTCCCTGCCTCTATATCT | TT | CACCCATCCCGGTTAAATGGTCCAT | CCCCTAAATACATTTGGCCTCCGTT | TT | CCACTCAACTTTAACCCG |
| 8 | GTCCCTGCCTCTATATCT | TT | AGGTCCAAGTTGGCCAAATTGGGGT | CCCATTTGCTGTTGGCAAGCGTTAA | TT | CCACTCAACTTTAACCCG |
| 9 | GTCCCTGCCTCTATATCT | TT | TAGAGAATTAACTGGAGAGAGCGTC | ATCGACGCAACCCATACCAGGGGAT | TT | CCACTCAACTTTAACCCG |
| 10 | GTCCCTGCCTCTATATCT | TT | CACCGCCTTCATTGACCGGCATCAT | TGGATGGTTTCCTCCTGGTACTACC | TT | CCACTCAACTTTAACCCG |
| 11 | GTCCCTGCCTCTATATCT | TT | TTAGACATCTGCTGACTCGGTGGCA | GGGGAACCGATAGTAGCGTTAGACA | TT | CCACTCAACTTTAACCCG |
| 12 | GTCCCTGCCTCTATATCT | TT | ATCCCGATTTGCGTAATTTTCAAGA | CGGTAACCGATCCATATGATCAGTT | TT | CCACTCAACTTTAACCCG |
| 13 | GTCCCTGCCTCTATATCT | TT | CTCCGTGGCTAAGTAGAACCCTGAC | CTTTATTGTCTTGAGCATCTCTGTT | TT | CCACTCAACTTTAACCCG |
| 14 | GTCCCTGCCTCTATATCT | TT | ATGAGATCTTCCACCATGCCCTCAA | GCCGCATTGATGTCAGCTTCAGCAT | TT | CCACTCAACTTTAACCCG |
| 15 | GTCCCTGCCTCTATATCT | TT | CTGAAAAACGCCTTGGGCATCGGCA | GTTGGTGGCACGATTTCGAAGCAAA | TT | CCACTCAACTTTAACCCG |
| 16 | GTCCCTGCCTCTATATCT | TT | AAGATTCTCCAGTTCGCTCGGTAGT | TGGCATAACGCGCGGCTAAGTGAAG | TT | CCACTCAACTTTAACCCG |
| 17 | GTCCCTGCCTCTATATCT | TT | ACCGCCTCGGAAAGACGCCAACATT | TTCTTCGTCTTCGCCAGTGTCCAAA | TT | CCACTCAACTTTAACCCG |
| 18 | GTCCCTGCCTCTATATCT | TT | TCCACGTTGCCATGATCGAGATCAC | GGAGTAAAACCGGCTGGTCCTTGTA | TT | CCACTCAACTTTAACCCG |
| 19 | GTCCCTGCCTCTATATCT | TT | GTTCCTAACATCGGCGGCGTCCAAA | AGGAGGAGTCAAGGCCAAAATGTCT | TT | CCACTCAACTTTAACCCG |
| 20 | GTCCCTGCCTCTATATCT | TT | TCGTCATAGTCGGCGGTAGTCATGA | TGTTGAGTCCATGGTCGAGGATCGT | TT | CCACTCAACTTTAACCCG |

**Table S6.** Probe pairs designed for *Phalangium opilio engrailed* HCR in situ hybridization (B1 initiator).

| Pair | Initiator | Spacer | Hybridization | Hybridization | Spacer | Initiator |
|---|---|---|---|---|---|---|
| 1 | GAGGAGGGCAGCAAACGG | AA | ACCTTCGCCATCCATGGCGACGGTA | CTATGATTTGTCCATTGAATCATCT | AA | GAAGAGTCTTCCTTTACG |
| 2 | GAGGAGGGCAGCAAACGG | AA | TCTGAAACCAAATTTTAATTTGGGA | TGGCCTTCTTGATTTTGGCCCGTTT | AA | GAAGAGTCTTCCTTTACG |
| 3 | GAGGAGGGCAGCAAACGG | AA | GAACTCTTGTTTTAGCCGGGCCAAC | TTTTTCCGTCAAGTATCTGTTTTCC | AA | GAAGAGTCTTCCTTTACG |
| 4 | GAGGAGGGCAGCAAACGG | AA | TTCTCATCAGGTTTCTTTTCCTTCT | TCGGCGGTGAACGCTGTCCTGGGTC | AA | GAAGAGTCTTCCTTTACG |
| 5 | GAGGAGGGCAGCAAACGG | AA | CAGATGAGGGTCTGTCGGAGTATCG | TCATGCGTCGGGACCGGGGACCTGG | AA | GAAGAGTCTTCCTTTACG |
| 6 | GAGGAGGGCAGCAAACGG | AA | GGGTCCCTTTTTGCCCGTTTCGCCG | GCAATAGACCCAGGCGGGCCATAAA | AA | GAAGAGTCTTCCTTTACG |
| 7 | GAGGAGGGCAGCAAACGG | AA | GGCGACTGGCTGGTTTCGGGCGTGG | TCTTGGGGCGCCAAGGAGGGCAATT | AA | GAAGAGTCTTCCTTTACG |
| 8 | GAGGAGGGCAGCAAACGG | AA | AGTTAACGGAACTACAGTTCGAACT | CGCCGACGCTCCGAGGCCTAACGGA | AA | GAAGAGTCTTCCTTTACG |
| 9 | GAGGAGGGCAGCAAACGG | AA | ATGATTGCGATGGTGTTCGCGATTG | ACTTTCGGGTTGTTGAACGCGCCGG | AA | GAAGAGTCTTCCTTTACG |
| 10 | GAGGAGGGCAGCAAACGG | AA | ACCGCCGAGAAAAAGTCCGTCGGAG | TTGCTGGCGGGTGTTGGCGGACCGG | AA | GAAGAGTCTTCCTTTACG |
| 11 | GAGGAGGGCAGCAAACGG | AA | TATGAACGAGTCCGTTATGAGTCCT | GGGGCACGGCCGCGTGATGTAGGTG | AA | GAAGAGTCTTCCTTTACG |

| | | | | | | |
|---|---|---|---|---|---|---|
| 12 | GAGGAGGGCAGCAAACGG | AA | TTTAGGAGTTTTGCCGAATTCCGGC | GACTTCGGCCGCGATGGGCGACCCG | AA | GAAGAGTCTTCCTTTACG |
| 13 | GAGGAGGGCAGCAAACGG | AA | GAGTGCCTGGAAAAGGGCGCCCCAA | AAAATCTTCTCGATGGAAAACTTGA | AA | GAAGAGTCTTCCTTTACG |
| 14 | GAGGAGGGCAGCAAACGG | AA | AGTTCTCTACGGCGCCGGGGAAGGA | CGGCTGGTATGACCGGAGGCGCCCG | AA | GAAGAGTCTTCCTTTACG |
| 15 | GAGGAGGGCAGCAAACGG | AA | ATCGTCGACGCGGGAACTGTCCGCG | ATTTCGCTGAGGAGGCGGCGTCGTT | AA | GAAGAGTCTTCCTTTACG |
| 16 | GAGGAGGGCAGCAAACGG | AA | CGCGCCGGGCTGCTGATCGACGTTC | GACGACGTACAGTCCAAGTCGTCGG | AA | GAAGAGTCTTCCTTTACG |
| 17 | GAGGAGGGCAGCAAACGG | AA | CCATAGCGCCCGCTTTTCGAGACCG | GCTTGGCCTCTACCATCGTATCTAA | AA | GAAGAGTCTTCCTTTACG |
| 18 | GAGGAGGGCAGCAAACGG | AA | CGACTACGGTGGTCTTACCCGAAGA | GGGGTCCGATCTTTCTCCCCGCCGC | AA | GAAGAGTCTTCCTTTACG |
| 19 | GAGGAGGGCAGCAAACGG | AA | TTCTCTCTCGTAATAAGTAAAACTC | ACTCGAACCTCGAGTAAGATTGAAA | AA | GAAGAGTCTTCCTTTACG |

**Table S7.** Probe pairs designed for *Phalangium opilio Distal-less* HCR in situ hybridization (B3 initiator).

| Pair | Initiator | Spacer | Hybridization | Hybridization | Spacer | Initiator |
|---|---|---|---|---|---|---|
| 1 | GTCCCTGCCTCTATATCT | TT | TCGAAGAACTCCAGCGATTGATTAT | TTTTTCCGATAGGTTTTCGGGATGA | TT | CCACTCAACTTTAACCCG |
| 2 | GTCCCTGCCTCTATATCT | TT | CCCGTGTGATGGTTGCGCATTTGTC | CATATGCACGATGTTCACGTGTTCA | TT | CCACTCAACTTTAACCCG |
| 3 | GTCCCTGCCTCTATATCT | TT | TCTGGGAACGGTAATAGCTCGTTCC | CCCGGTTTGGTTTACTTCTTGTCGC | TT | CCACTCAACTTTAACCCG |
| 4 | GTCCCTGCCTCTATATCT | TT | ATTCGTGAGTGATTGGTTTTTGTTT | GAACTCGAAGTAACGCGAAGAGGGT | TT | CCACTCAACTTTAACCCG |
| 5 | GTCCCTGCCTCTATATCT | TT | TTCACGGGGTTATGTTAAAATTTGG | AATATAAACCACGAGACGTGACGTA | TT | CCACTCAACTTTAACCCG |
| 6 | GTCCCTGCCTCTATATCT | TT | CAAGAATACTGTGGCATGTAACCGT | GTTAACGAAGGGTCCGCTTGGTGAT | TT | CCACTCAACTTTAACCCG |
| 7 | GTCCCTGCCTCTATATCT | TT | GATTCATGTCCCAAGAACTAATAGG | TATGCATGTTCATAGCGGCTTTCGC | TT | CCACTCAACTTTAACCCG |
| 8 | GTCCCTGCCTCTATATCT | TT | CCCGATTGGGTCATCATGCCCGATT | GCGTTGGACATGCTATTTACGGGCG | TT | CCACTCAACTTTAACCCG |
| 9 | GTCCCTGCCTCTATATCT | TT | TCGTTGCCGGGTTTGGTTGCTGTAA | AATGTCCTTCCGACGGAGTTTGAGG | TT | CCACTCAACTTTAACCCG |
| 10 | GTCCCTGCCTCTATATCT | TT | CTTGTTGTTGAGCTTTCAACATTTT | GACCGCCCGGGCCCGAATTCGGAGG | TT | CCACTCAACTTTAACCCG |
| 11 | GTCCCTGCCTCTATATCT | TT | GATTTTCACCTGCGTTTGCGTTAGA | GTACTTGGAACGGCGATTTTGGAAC | TT | CCACTCAACTTTAACCCG |
| 12 | GTCCCTGCCTCTATATCT | TT | GGTAACGCCAAGTACTGCGTTCTTT | AGAGACGCGGCTAATTCCGCTCTTT | TT | CCACTCAACTTTAACCCG |
| 13 | GTCCCTGCCTCTATATCT | TT | GACTCGAGTATATCGTTCTGGGCTT | ACCTTCTGTTTAATTGCTGTAATTG | TT | CCACTCAACTTTAACCCG |
| 14 | GTCCCTGCCTCTATATCT | TT | GTCTCTTCTAAACCAGATTTGTCGT | TTTTTGCCCTTTCCATTGACTCTCA | TT | CCACTCAACTTTAACCCG |
| 15 | GTCCCTGCCTCTATATCT | TT | TGGCGAGGTAATGGCTAGGATGAGT | CACACGAACCGACGTTGGTCGGATA | TT | CCACTCAACTTTAACCCG |
| 16 | GTCCCTGCCTCTATATCT | TT | GGAATTGTTCATGGAAAAGGGGTAG | CGAATTATGCAAACCAGAGTTTAAC | TT | CCACTCAACTTTAACCCG |
| 17 | GTCCCTGCCTCTATATCT | TT | GAATTTCCATAAAGGCCGACTTGTT | CATTCAGTCCGGCCGCCGCCTGCTG | TT | CCACTCAACTTTAACCCG |
| 18 | GTCCCTGCCTCTATATCT | TT | GGTGGCGGCTTTGGCGGAACAACGT | AGCCGAGAACTCGCTAAAATTCAAC | TT | CCACTCAACTTTAACCCG |

**Table S8.** Probe pairs designed for *Pselaphochernes parvus exd-1* HCR in situ hybridization (B2 initiator).

| Pair | Initiator | Spacer | Hybridization | Hybridization | Spacer | Initiator |
|---|---|---|---|---|---|---|
| 1 | CCTCGTAAATCCTCATCA | AA | CACCCAAAGCTCTTGATTAAAATAA | GGGGACACTCACGACAAATGCGAGT | AA | ATCATCCAGTAAACCGCC |
| 2 | CCTCGTAAATCCTCATCA | AA | CGCCTATCCTCCATAGGCGAGGGAG | TTTTTACAAGAAGTCGAACCCTAAG | AA | ATCATCCAGTAAACCGCC |
| 3 | CCTCGTAAATCCTCATCA | AA | CTGGCACCCATGGAGGAATACGAGT | CAGATTTTGTAGTCCTGGGATGGCA | AA | ATCATCCAGTAAACCGCC |

| | | | | | | |
|---|---|---|---|---|---|---|
| 4 | CCTCGTAAATCCTCATCA | AA | GAGTGGAAACCATGTTATAGGGAGA | GGGGCGGACTAATCATGGGTCCCTG | AA | ATCATCCAGTAAACCGCC |
| 5 | CCTCGTAAATCCTCATCA | AA | CTTCTTCCTGAGCTTTGCCGATATT | CCGCTTTCTTGGCGGCGTACAGGTT | AA | ATCATCCAGTAAACCGCC |
| 6 | CCTCGTAAATCCTCATCA | AA | CCAATTGGAGACCTGAGATACGGTG | CTTGTACCGGATTCTCTTGTTGCCG | AA | ATCATCCAGTAAACCGCC |
| 7 | CCTCGTAAATCCTCATCA | AA | CCTCTTCACTGGGGTACGGATTGCT | AACACTTCCTAGCCAGCTCTTCCTT | AA | ATCATCCAGTAAACCGCC |
| 8 | CCTCGTAAATCCTCATCA | AA | CTCGGTCGCCTGTTTGCTGAAGTTT | GTGCGAGTAGAAGTACTCGTTGAGG | AA | ATCATCCAGTAAACCGCC |
| 9 | CCTCGTAAATCCTCATCA | AA | ATCTGAGGATCATAACGGCTTCACA | GTTTCCGCCTTGCGTCGAGGAACCT | AA | ATCATCCAGTAAACCGCC |
| 10 | CCTCGTAAATCCTCATCA | AA | GTTGAATTTGCGGTGGATGATCTGA | GGACTGCTTGAGTTGGACTTGGATG | AA | ATCATCCAGTAAACCGCC |
| 11 | CCTCGTAAATCCTCATCA | AA | ACTCATTGCAGGCCTGCTCGTACTT | TCAGCAGGTTCATGACGTGGGTGGT | AA | ATCATCCAGTAAACCGCC |
| 12 | CCTCGTAAATCCTCATCA | AA | TCGGGAGGCTCCTCATCTTGTGTGT | ATGTTGTCCAGCCGCATAAGCTGAG | AA | ATCATCCAGTAAACCGCC |
| 13 | CCTCGTAAATCCTCATCA | AA | GAAGAGTGCAGGTTTCATGCGATGG | TTTTTCTTTGATTTCACATAGAACA | AA | ATCATCCAGTAAACCGCC |
| 14 | CCTCGTAAATCCTCATCA | AA | TCATCCAGACTTTGGTCCGTTATAT | TTGAGCGTATGTTTTCTTGCTTGAG | AA | ATCATCCAGTAAACCGCC |
| 15 | CCTCGTAAATCCTCATCA | AA | GTTCAGAAGGTTGTTGTCCATTAGG | CTGAAATGTCATGTTTTCTTACATC | AA | ATCATCCAGTAAACCGCC |
| 16 | CCTCGTAAATCCTCATCA | AA | TGCATGCTGAGGCACGACATGACCC | AACTTGATGTGGTTGAGGCATGCCA | AA | ATCATCCAGTAAACCGCC |
| 17 | CCTCGTAAATCCTCATCA | AA | GGATGAAGCATTCTTTGTTGCTCGT | ATGCTAACGGCAGGGTGCTGAGTAA | AA | ATCATCCAGTAAACCGCC |
| 18 | CCTCGTAAATCCTCATCA | AA | GGAAGGACATCAGTGATGAAAAACA | TGCTTTAACATCACCCTCGGTGAGC | AA | ATCATCCAGTAAACCGCC |
| 19 | CCTCGTAAATCCTCATCA | AA | AATAAACATCCGCGAGTACAAAAAC | GCTGGGCAGGTCGTCAGTTGCAAGC | AA | ATCATCCAGTAAACCGCC |
| 20 | CCTCGTAAATCCTCATCA | AA | AAACGTTTGACGGACAACAAGCAAC | TAATCACCGCAATAAGAGTATTTTG | AA | ATCATCCAGTAAACCGCC |

**Table S9.** Probe pairs designed for *Pselaphochernes parvus exd-2* HCR in situ hybridization (B2 initiator).

| Pair | Initiator | Spacer | Hybridization | Hybridization | Spacer | Initiator |
|---|---|---|---|---|---|---|
| 1 | GTCCCTGCCTCTATATCT | TT | TCTGGATATGAGGTTCCCTAAGAAG | AATGGTGCTTATATAATGTTTTGTC | TT | CCACTCAACTTTAACCCG |
| 2 | GTCCCTGCCTCTATATCT | TT | ACTTACACTGTACAGAGCGTTTTTC | AACCGATAGAATGATTGTGGTAACA | TT | CCACTCAACTTTAACCCG |
| 3 | GTCCCTGCCTCTATATCT | TT | AATTTCCGCCTGATGGTAGAGTAGG | GGGGATCTCCACAAATAGAAAAATA | TT | CCACTCAACTTTAACCCG |
| 4 | GTCCCTGCCTCTATATCT | TT | TATATACAGTAGATACCAAATGTCA | GAAAGAGGTGCTCTGCAGTTAGTTG | TT | CCACTCAACTTTAACCCG |
| 5 | GTCCCTGCCTCTATATCT | TT | TCTTGAGCTTTGCCAATGTTCTTTT | TTTTTGGCGGCGTAGAGGTTGGCCT | TT | CCACTCAACTTTAACCCG |
| 6 | GTCCCTGCCTCTATATCT | TT | TGGAGACCTGAGAGACAGTGATGCC | ATCTTATCCTCTTGTTTCCGAACCA | TT | CCACTCAACTTTAACCCG |
| 7 | GTCCCTGCCTCTATATCT | TT | CTCGCTAGGGTAAGGGTTACTCAAG | CTTCCTCGCCAATTCCTCCTTGGCC | TT | CCACTCAACTTTAACCCG |
| 8 | GTCCCTGCCTCTATATCT | TT | GTCGCCTGCTTGCTGAAGTTGCGCC | GAGTAGAAGTATTCGTTCAGGATCT | TT | CCACTCAACTTTAACCCG |
| 9 | GTCCCTGCCTCTATATCT | TT | TGAGGATCATGACCGCTTCACACGT | TCCTCCTGGCGTCCAAGAACCGGGA | TT | CCACTCAACTTTAACCCG |
| 10 | GTCCCTGCCTCTATATCT | TT | GAACTTGCGATGGATGATCTGGACC | CTGCTTGAGTTGAACCTGGATGGAG | TT | CCACTCAACTTTAACCCG |
| 11 | GTCCCTGCCTCTATATCT | TT | CTCGTCCGGCTCTGCTCCCGTAGTA | CGCTCGATCTCCTTGGGCGTGATGG | TT | CCACTCAACTTTAACCCG |
| 12 | GTCCCTGCCTCTATATCT | TT | AGGCCTGCTCGTATTTCTCGAGTTC | TCATGACGTGGGTGGTGAACTCGTT | TT | CCACTCAACTTTAACCCG |
| 13 | GTCCCTGCCTCTATATCT | TT | TTTGGCGCGGTAGTCCGAATGCTCG | GTGATATATAGTACGGATCTGCGCC | TT | CCACTCAACTTTAACCCG |
| 14 | GTCCCTGCCTCTATATCT | TT | GCCGCCGAAGAAACATTCGCTGCAG | GCATTCTCGGCTTGGCCTCCCGACG | TT | CCACTCAACTTTAACCCG |
| 15 | GTCCCTGCCTCTATATCT | TT | CGACGCCCTCGGCTACCAGCATGTT | CGGCACCACCTCCCTTTTCGGGTCC | TT | CCACTCAACTTTAACCCG |
| 16 | GTCCCTGCCTCTATATCT | TT | GAAGAGTGCAGGCTTCATCCGATGG | TTTTTCTTTGATCTCACAGAGAACA | TT | CCACTCAACTTTAACCCG |
| 17 | GTCCCTGCCTCTATATCT | TT | TCATCCAAGCTCTGGTCGGTAATAT | TTGAGGGTATGTTTCCTTGCTTGAG | TT | CCACTCAACTTTAACCCG |

| | | | | | | |
|---|---|---|---|---|---|---|
| 18 | GTCCCTGCCTCTATATCT | TT | ACGACGGGGAGGTAACCGCCAGGGT | GGGGTCGCAGTCTCGGGGCTTTGGT | TT | CCACTCAACTTTAACCCG |
| 19 | GTCCCTGCCTCTATATCT | TT | TTTCCATCGCGGGTCACATCCATAG | GCATACCGGGAGGATGTAACATATT | TT | CCACTCAACTTTAACCCG |
| 20 | GTCCCTGCCTCTATATCT | TT | TTTCTCAACGCATATAGAAGCCACG | CGCACGCTGGACCGCGGCGACGTAC | TT | CCACTCAACTTTAACCCG |

**Table S10.** Probe pairs designed for *Pselaphochernes parvus dac-1* HCR in situ hybridization (B3 initiator).

| Pair | Initiator | Spacer | Hybridization | Hybridization | Spacer | Initiator |
|---|---|---|---|---|---|---|
| 1 | GTCCCTGCCTCTATATCT | TT | GTTTAATTGGAGGTGGTTTGACATG | TTGGAATCCTACTTAGATGAAATGA | TT | CCACTCAACTTTAACCCG |
| 2 | GTCCCTGCCTCTATATCT | TT | ACAGGGTTCTCTTATCATCATCGCG | TAGATAACTTTCACTAATGATGATG | TT | CCACTCAACTTTAACCCG |
| 3 | GTCCCTGCCTCTATATCT | TT | TTTCTTTCTCAAAACTTAGCGACTA | CCCCGTATCAAAATATCCCAAGTCA | TT | CCACTCAACTTTAACCCG |
| 4 | GTCCCTGCCTCTATATCT | TT | CAAAGCGATGCCTCAGAATGTTTTG | GCGAGCCTTGTCTTCTAACGAACGA | TT | CCACTCAACTTTAACCCG |
| 5 | GTCCCTGCCTCTATATCT | TT | ACCAACACGGGATCTTTTATAAATG | ACCTTCCAACACTTGATGGACCGTA | TT | CCACTCAACTTTAACCCG |
| 6 | GTCCCTGCCTCTATATCT | TT | GTTTATATTGGATGTGTTAATGAGG | TCAACGTTTGTTTTTCTCTCGTCTG | TT | CCACTCAACTTTAACCCG |
| 7 | GTCCCTGCCTCTATATCT | TT | GCCAGAGTGGGTTTTAAAATGATGG | GAGAATTTTGGGGTGTATAAAAATG | TT | CCACTCAACTTTAACCCG |
| 8 | GTCCCTGCCTCTATATCT | TT | CATACAAGACTTGGGAAACAGGTTT | GAACATCATGTGGTGCCTATGTATT | TT | CCACTCAACTTTAACCCG |
| 9 | GTCCCTGCCTCTATATCT | TT | TAGATGTTCAGGCTGACAAAGGACA | TATGGTAGGACTTGATATGAAGATA | TT | CCACTCAACTTTAACCCG |
| 10 | GTCCCTGCCTCTATATCT | TT | CCTAGTGTTTCTCTCTCGATCCAAT | ATCTTGTATTTGGTGCTCTTTTTCA | TT | CCACTCAACTTTAACCCG |
| 11 | GTCCCTGCCTCTATATCT | TT | AAGGTCTCCGCTGAGGTGTTGCGTA | TGGGCCAAGGAATCATTGAGGAGTC | TT | CCACTCAACTTTAACCCG |
| 12 | GTCCCTGCCTCTATATCT | TT | TCTCGTTCTCCAGTTGATCCTGGAC | CGTCTTCGTACTGGGAGCGCTTCTT | TT | CCACTCAACTTTAACCCG |
| 13 | GTCCCTGCCTCTATATCT | TT | CCTTTTCTGGTACACGATTCTTGCT | ACGACGTGATCGCTTTTCTCTCCGG | TT | CCACTCAACTTTAACCCG |
| 14 | GTCCCTGCCTCTATATCT | TT | CTCTCCCGAACTTCTCGTTCTCTTA | TGTTCGTCCATGAGCTGCTTCTCGA | TT | CCACTCAACTTTAACCCG |
| 15 | GTCCCTGCCTCTATATCT | TT | GGTTGACCTGTCGTTCTTGCTGTCG | TCTCCATCTTGAGCTCGGCTTTTTC | TT | CCACTCAACTTTAACCCG |
| 16 | GTCCCTGCCTCTATATCT | TT | TGTTCTCTGTCTTCGTCGTCTTCAT | CTCACGTCGGGGTTGTCGCTCAGCT | TT | CCACTCAACTTTAACCCG |
| 17 | GTCCCTGCCTCTATATCT | TT | GAACCAACCATAGGTGCCCATTGAT | AGAGATTGAGGACGGGACTGTGGCC | TT | CCACTCAACTTTAACCCG |
| 18 | GTCCCTGCCTCTATATCT | TT | TAGCCCGTAGAGACGTTCTTTCTGT | AGCGGTGTCTTTCTGTTTCTGACTC | TT | CCACTCAACTTTAACCCG |
| 19 | GTCCCTGCCTCTATATCT | TT | GTAAGGAGATCGGCAGACCGAGCTG | TGGTTTTCCTCCTTTAATCTGGCTG | TT | CCACTCAACTTTAACCCG |
| 20 | GTCCCTGCCTCTATATCT | TT | GGACTCCGTCCAGGCGGGACTTCTT | CGTTCTCATAGCTTCCATAGTCAGG | TT | CCACTCAACTTTAACCCG |
| 21 | GTCCCTGCCTCTATATCT | TT | CAGTCCTTGTACAGGACATCGAAGT | CTTCCAGGCCTGGAACTTGCCGTAG | TT | CCACTCAACTTTAACCCG |
| 22 | GTCCCTGCCTCTATATCT | TT | ACGATAGGAGTGATGTCCAATCGCT | AGGATTCGCACCTGCTCGACGTTGC | TT | CCACTCAACTTTAACCCG |
| 23 | GTCCCTGCCTCTATATCT | TT | CGACCAGATGTTTCAGGAAGAGTTC | GTTTGGTATAGACAGTGTGAAGACC | TT | CCACTCAACTTTAACCCG |

**Table S11.** Probe pairs designed for *Pselaphochernes parvus dac-2* HCR in situ hybridization (B3 initiator).

| Pair | Initiator | Spacer | Hybridization | Hybridization | Spacer | Initiator |
|---|---|---|---|---|---|---|
| 1 | GTCCCTGCCTCTATATCT | TT | ATAGAATTGAAATTAAGAGAGTGAG | GGGGTCATCACATATAAACAGGTAA | TT | CCACTCAACTTTAACCCG |
| 2 | GTCCCTGCCTCTATATCT | TT | CATCAAGGGCGCTATAACAATTTGA | TCTACAGAAGAGGCTAAGCTATCAC | TT | CCACTCAACTTTAACCCG |
| 3 | GTCCCTGCCTCTATATCT | TT | TGCGAAAGGAAGATTTTCCGCGCAG | CCCCACTCCTGGAGAAATGAAGAAC | TT | CCACTCAACTTTAACCCG |
| 4 | GTCCCTGCCTCTATATCT | TT | GGCTCGAAGGACTGGGGTTTATGCT | GGGGAATTCTTGATATGTTTTAGAA | TT | CCACTCAACTTTAACCCG |

| | | | | | | |
|---|---|---|---|---|---|---|
| 5 | GTCCCTGCCTCTATATCT | TT | GACTCCACTTTCACCATGCGATCAC | ATGGGGCATTCTTGAACTTTATGTT | TT | CCACTCAACTTTAACCCG |
| 6 | GTCCCTGCCTCTATATCT | TT | ACCACACGTTGAGGAGTCTCATGGT | CCAGTTCTTGGGTTAATGATTCTGC | TT | CCACTCAACTTTAACCCG |
| 7 | GTCCCTGCCTCTATATCT | TT | GTACTGCGCTCTCCTCTTCATTTCA | TGCCGAGCTGTTCCTCAGGGCATCT | TT | CCACTCAACTTTAACCCG |
| 8 | GTCCCTGCCTCTATATCT | TT | GTTTTCTTTTCTCGCTTGAATCTCT | TCCAGTTGGTCTTGCACTCGCCGCC | TT | CCACTCAACTTTAACCCG |
| 9 | GTCCCTGCCTCTATATCT | TT | CCAGAAGTTGTTTCTCGATGTTTTC | GGTAGAGGATGCGTATTCTTTGTTC | TT | CCACTCAACTTTAACCCG |
| 10 | GTCCCTGCCTCTATATCT | TT | CTTTAGCTCGGCTTTATCAATGGAG | CAGTTCCCGTTCTCGCATCATCTCC | TT | CCACTCAACTTTAACCCG |
| 11 | GTCCCTGCCTCTATATCT | TT | TTGGCGGATAGGTAAGGTGCAAGAA | AGACCCAAGCCAACATCTCCGGGAG | TT | CCACTCAACTTTAACCCG |
| 12 | GTCCCTGCCTCTATATCT | TT | GACATCGGGGTTGTCGCTCAGTTCT | TAAACGATCGGAGTTCGCCGTGGAA | TT | CCACTCAACTTTAACCCG |
| 13 | GTCCCTGCCTCTATATCT | TT | GAGTCGTTGTAGTCTTCTTCCGGTG | GGCTCTCGATCATCTTCATCATCTT | TT | CCACTCAACTTTAACCCG |
| 14 | GTCCCTGCCTCTATATCT | TT | GCTGCTGTTTTGGGACAAGTTGAGA | GGTGTAGGCTGTGTCTTGATCTCGC | TT | CCACTCAACTTTAACCCG |
| 15 | GTCCCTGCCTCTATATCT | TT | TTAAGGTATGCTTGATGTTGCTCCT | GGGCTGTGACCGTTGGTCAGGGTTC | TT | CCACTCAACTTTAACCCG |
| 16 | GTCCCTGCCTCTATATCT | TT | GCGCTATGTCTACCAGGTCAGATGA | TCCTTTCGTAGCCGTAATGCCGTTC | TT | CCACTCAACTTTAACCCG |
| 17 | GTCCCTGCCTCTATATCT | TT | AGAGTGCTTGATGACGGAGTGTTGA | TGAGGTCTGGAGGCTCAGTTCGGGA | TT | CCACTCAACTTTAACCCG |
| 18 | GTCCCTGCCTCTATATCT | TT | TTCCTCCTGTCCCTTCTTCAGCCAA | GGAACCCGTTGGCCACCATTTTGTC | TT | CCACTCAACTTTAACCCG |
| 19 | GTCCCTGCCTCTATATCT | TT | CCCAGTTCTGTGATATGTCCGTTCT | TCACGTTTGTACCAGGACGGACGCG | TT | CCACTCAACTTTAACCCG |
| 20 | GTCCCTGCCTCTATATCT | TT | CTAGACGGGATTTCTTCAGCAGCTG | ACGAAGTCGTCCCAAAGTCCCCTGC | TT | CCACTCAACTTTAACCCG |
| 21 | GTCCCTGCCTCTATATCT | TT | CCTTGTAGAGGCTGTCAAAGTCTTT | GGCGGCCAGGCCTGGCAGTTGTACA | TT | CCACTCAACTTTAACCCG |
| 22 | GTCCCTGCCTCTATATCT | TT | CTGACTTGTTCCACGTTACAGACAA | TGGATGGCGCCCAGTCCACGAAGGA | TT | CCACTCAACTTTAACCCG |
| 23 | GTCCCTGCCTCTATATCT | TT | TGTAGACTGTGTGGAGGCCTCCCAC | GCATGATATCGAGCCGTTTCAGTTT | TT | CCACTCAACTTTAACCCG |
| 24 | GTCCCTGCCTCTATATCT | TT | AGGTAAACAAAGTAGATAATCGCCG | ATGTTTCAGGAAGAGTTCAAAAGCT | TT | CCACTCAACTTTAACCCG |

**Table S12.** Probe pairs designed for *Titanopuga salinarum exd* HCR in situ hybridization (B3 initiator).

| Pair | Initiator | Spacer | Hybridization | Hybridization | Spacer | Initiator |
|---|---|---|---|---|---|---|
| 1 | GTCCCTGCCTCTATATCT | TT | TCCTTTATGCTCCGACACGAACATA | GTGCCTTAACGCAGTCGCTTGTCTC | TT | CCACTCAACTTTAACCCG |
| 2 | GTCCCTGCCTCTATATCT | TT | GAAATTTGACCACCGGTACCAGGCG | AGTCTTCCCAGTGCCTCTCGTCCTG | TT | CCACTCAACTTTAACCCG |
| 3 | GTCCCTGCCTCTATATCT | TT | CTCCCATTGACGAGTAGGTATCGCC | GAATGCCCAACTGAGATTGGACGTT | TT | CCACTCAACTTTAACCCG |
| 4 | GTCCCTGCCTCTATATCT | TT | CTGCGCTCCTGGTGGCGGTGGAGGT | GTTCATACTCATGTTGTACATGCCG | TT | CCACTCAACTTTAACCCG |
| 5 | GTCCCTGCCTCTATATCT | TT | TGGGTGGTTGGCACCATATTGTAGG | CTAATCATTTGACCCGGTTGCTGAC | TT | CCACTCAACTTTAACCCG |
| 6 | GTCCCTGCCTCTATATCT | TT | TCCTGAGCTTTGCCGATGTTCTTCT | TTTTTGGCTGCGTAGAGATTGGCTT | TT | CCACTCAACTTTAACCCG |
| 7 | GTCCCTGCCTCTATATCT | TT | TGGACACCTGCGAAACCGTGATACC | ACCTTATCCGTTTGTTCCCAAACCA | TT | CCACTCAACTTTAACCCG |
| 8 | GTCCCTGCCTCTATATCT | TT | TTCACTAGGATAGGGATTACTAAGA | TTTCCTCGCCAGTTCCTCTTTCGCC | TT | CCACTCAACTTTAACCCG |
| 9 | GTCCCTGCCTCTATATCT | TT | GTCGCCTGTTTACTGAAGTTTCTCC | GAATAGAAGTACTCGTTGAGGATCT | TT | CCACTCAACTTTAACCCG |
| 10 | GTCCCTGCCTCTATATCT | TT | TGAGGATCATGACGGCTTCGCAAGT | TCCGCCTGGCGTCCAAGAACCTGGA | TT | CCACTCAACTTTAACCCG |
| 11 | GTCCCTGCCTCTATATCT | TT | GAACTTCTTGTGAATAATTTGTACC | CTGCTTCAGCTGGACTTGTATAGAG | TT | CCACTCAACTTTAACCCG |
| 12 | GTCCCTGCCTCTATATCT | TT | CGCGTCCGACTTTGTTCCCGAAGCA | CGTTCAATCTCCTTCGGCGTAATCG | TT | CCACTCAACTTTAACCCG |
| 13 | GTCCCTGCCTCTATATCT | TT | ACGCCTGTTCATATTTTTCTAGTTC | TCATTACGTGCGTCGTGAACTCATT | TT | CCACTCAACTTTAACCCG |
| 14 | GTCCCTGCCTCTATATCT | TT | TTTTGCCCTGTAATCCGAATGTTCG | GTGATAAATTTGCCTAATTTGTGCA | TT | CCACTCAACTTTAACCCG |

| | | | | | | |
|---|---|---|---|---|---|---|
| 15 | GTCCCTGCCTCTATATCT | TT | GAAGCCGCTGCTGACGTATTCGCCG | GCGTTTTCCGGCTGACCTGGTCCAC | TT | CCACTCAACTTTAACCCG |
| 16 | GTCCCTGCCTCTATATCT | TT | CGGCTACCCCTTCCGCTATGAGCAT | CCCCTGCGCCTCCACCCTTTTCGGG | TT | CCACTCAACTTTAACCCG |
| 17 | GTCCCTGCCTCTATATCT | TT | AGGCTCTTCTTCTTGTGTGTTCCGT | ATCTAACCTCATTAGTTGTGGATCT | TT | CCACTCAACTTTAACCCG |
| 18 | GTCCCTGCCTCTATATCT | TT | GAACAGTGCTGGCTTCATACGGTGA | TTTTTCCTTGATCTCGCAGAGCACA | TT | CCACTCAACTTTAACCCG |
| 19 | GTCCCTGCCTCTATATCT | TT | TCGTCCAAACTCTGGTCCGTAATAT | TTCAACGTATGCTTCCTGGCTTGCG | TT | CCACTCAACTTTAACCCG |

**Table S13.** Probe pairs designed for *Titanopuga salinarum dac* HCR in situ hybridization (B2 initiator).

| Pair | Initiator | Spacer | Hybridization | Hybridization | Spacer | Initiator |
|---|---|---|---|---|---|---|
| 1 | CCTCGTAAATCCTCATCA | AA | TGTCCGTTAATTGTACTACCGTGCA | TTTTTCTCTGCCCATGTTGCGGGG | AA | ATCATCCAGTAAACCGCC |
| 2 | CCTCGTAAATCCTCATCA | AA | CCAGGTGTTTGCGAAGTCCTTTTTT | GGGGAATGATTCGTGATTTGCGTGT | AA | ATCATCCAGTAAACCGCC |
| 3 | CCTCGTAAATCCTCATCA | AA | TATACAGACATGGGGCAGTCTTGCA | ATGGTTGGATGGGAGGTTGACTTCA | AA | ATCATCCAGTAAACCGCC |
| 4 | CCTCGTAAATCCTCATCA | AA | TCTCCATCTCCAATTCTTGCGTTAG | TCCTTTCAGCGTCCGGTCTGCTGCT | AA | ATCATCCAGTAAACCGCC |
| 5 | CCTCGTAAATCCTCATCA | AA | GGAAGTGTTCCGAAGTGCCTCTTCG | CTCGTTGAGTAGACGTAGCGTTTCG | AA | ATCATCCAGTAAACCGCC |
| 6 | CCTCGTAAATCCTCATCA | AA | AGCTGCTCGTGAAGCCGACGCCGAC | TGCGTTCGCTTCTTCATCTCGGACT | AA | ATCATCCAGTAAACCGCC |
| 7 | CCTCGTAAATCCTCATCA | AA | AGAGAATTCTCGTCCTCTGTTCTTC | TCTTCTCTTTCTTCAACCTCTTTTG | AA | ATCATCCAGTAAACCGCC |
| 8 | CCTCGTAAATCCTCATCA | AA | TTCCCTTTCCCGTAAGATCTCCATC | TAACTGTTTCTCTAAGTTCTCCCTG | AA | ATCATCCAGTAAACCGCC |
| 9 | CCTCGTAAATCCTCATCA | AA | CTTTCTTGCTGTCGGGCGTTGTCTG | AACTCCGCTTTCTCCAAGTTAATCT | AA | ATCATCCAGTAAACCGCC |
| 10 | CCTCGTAAATCCTCATCA | AA | TAAGAAGTGTTTCTATGGACGATAC | CTACCTTGAGCAAACCCTGGATGTT | AA | ATCATCCAGTAAACCGCC |
| 11 | CCTCGTAAATCCTCATCA | AA | TCCTGTGCTCTGGAAAGGCGCCGAT | TGGATTCGTGGTTGCGCTCCCATCG | AA | ATCATCCAGTAAACCGCC |
| 12 | CCTCGTAAATCCTCATCA | AA | GTACTACTGACGTCAGGGTTATCAC | CTGGAAGTGAGTCGATCGGAATTGG | AA | ATCATCCAGTAAACCGCC |

**Table S14.** Probe pairs designed for *Archegozetes longisetosus dac* HCR in situ hybridization (B2 initiator).

| Pair | Initiator | Spacer | Hybridization | Hybridization | Spacer | Initiator |
|---|---|---|---|---|---|---|
| 1 | CCTCGTAAATCCTCATCA | AA | TCCGTTTGTTGTTGTTGCTGTTGCG | TAATTGCAAAACTTTTAATCTACTG | AA | ATCATCCAGTAAACCGCC |
| 2 | CCTCGTAAATCCTCATCA | AA | CTTGTTGGTTTTGAGTGATTTGCTG | GGGGAGTCGGTGACGATTTCAGAGA | AA | ATCATCCAGTAAACCGCC |
| 3 | CCTCGTAAATCCTCATCA | AA | CTGCTGCTGATGTATCTCGTGCTCC | ATTCTGTGTATTGTGTGATATCTGT | AA | ATCATCCAGTAAACCGCC |
| 4 | CCTCGTAAATCCTCATCA | AA | CAAATGCTCTCTCATTTCTCGCTCC | TTTTTGTTCTTCATTCAGTTGTTTT | AA | ATCATCCAGTAAACCGCC |
| 5 | CCTCGTAAATCCTCATCA | AA | AGTTCAGCTTTTTGCAAATGCAATT | TCAAGTTGTGCCGCCAACTCAGCCC | AA | ATCATCCAGTAAACCGCC |
| 6 | CCTCGTAAATCCTCATCA | AA | CAATTGCCAAAAGCCCCTCAATATT | TTTGTTGTTGTCTAGCATTATGAGC | AA | ATCATCCAGTAAACCGCC |
| 7 | CCTCGTAAATCCTCATCA | AA | TTGTTGAGTTTGCTGTTGCTGTGGA | CAGCAATGTTTCCAAACAATGTTCC | AA | ATCATCCAGTAAACCGCC |
| 8 | CCTCGTAAATCCTCATCA | AA | TTGCGGTTTCACTCGCACCCCGTGT | GAGGAGTTGTCGAACTGTTTCGATT | AA | ATCATCCAGTAAACCGCC |
| 9 | CCTCGTAAATCCTCATCA | AA | GCTGACAGGGTTGTCGCTATTGTTA | ACACGATATACCGTTAAATCCAGAA | AA | ATCATCCAGTAAACCGCC |
| 10 | CCTCGTAAATCCTCATCA | AA | TCGTCGTCATCATCAGTAGTGTCAT | CTATTATTACTTTCCACGTCGTCTT | AA | ATCATCCAGTAAACCGCC |
| 11 | CCTCGTAAATCCTCATCA | AA | TGCGCTGCTGGTGTTACTGCTGCTG | ATTATTATTATTGTTTGTATTGTTG | AA | ATCATCCAGTAAACCGCC |
| 12 | CCTCGTAAATCCTCATCA | AA | GTTACGAGAGCTTAAGTTAAGTGCG | GTTATTTCCACTACTATTGACATTA | AA | ATCATCCAGTAAACCGCC |
| 13 | CCTCGTAAATCCTCATCA | AA | TGTTGCTGTGAAAGTAACCATAAAT | GAGTTCGGGACATGTTGTTGTTGCT | AA | ATCATCCAGTAAACCGCC |

| | | | | | | |
|---|---|---|---|---|---|---|
| 14 | CCTCGTAAATCCTCATCA | AA | CCAGAACCTTCCGATGTGGCGTTGT | GTTGGATTGAAATCTGAGGCCACGG | AA | ATCATCCAGTAAACCGCC |
| 15 | CCTCGTAAATCCTCATCA | AA | CTGGAGGCGGACTTGCATTTGAAAA | CATTGCTTCCGTGATGCGAAGCTAT | AA | ATCATCCAGTAAACCGCC |
| 16 | CCTCGTAAATCCTCATCA | AA | TGGTGCGGTTCCAGCGCTGAAGTGT | CACAAGTTGTTGCTGTTGCTGAGGA | AA | ATCATCCAGTAAACCGCC |
| 17 | CCTCGTAAATCCTCATCA | AA | TTCGCTGCTAGCATTGCTGCTGCGG | AAGAAAGGAAATGCGGCCGAATAAC | AA | ATCATCCAGTAAACCGCC |
| 18 | CCTCGTAAATCCTCATCA | AA | TATTAAGACTGCTATGCGTTTGCTT | CGGCACTTGTGTTACCACTACTATT | AA | ATCATCCAGTAAACCGCC |
| 19 | CCTCGTAAATCCTCATCA | AA | ATCATCAGAGAAACGGGGCTTTTTA | GTCTATATTTTCACTGGCATAGTCA | AA | ATCATCCAGTAAACCGCC |
| 20 | CCTCGTAAATCCTCATCA | AA | TTGTGTTGCTGTGGCCCTGAGGTGA | GTTGGGTATCCGAGATAGGGCACAA | AA | ATCATCCAGTAAACCGCC |
| 21 | CCTCGTAAATCCTCATCA | AA | TGACTGAACTTCTCTTGGGTGGACG | ATGATGGGACTGGGCTAACAATGCC | AA | ATCATCCAGTAAACCGCC |

**Table S15.** Probe pairs designed for *Ixodes scapularis exd* HCR in situ hybridization (B2 initiator).

| Pair | Initiator | Spacer | Hybridization | Hybridization | Spacer | Initiator |
|---|---|---|---|---|---|---|
| 1 | CCTCGTAAATCCTCATCA | AA | TTCCGAACTCTGCGGCGGACTGTAG | TCACAGCGGACTGAGCGGGGTTGAG | AA | ATCATCCAGTAAACCGCC |
| 2 | CCTCGTAAATCCTCATCA | AA | GATTGCACGTTGGCGCCCATGGCGT | ATGGCATGCATGCCGTGGGGAAGCT | AA | ATCATCCAGTAAACCGCC |
| 3 | CCTCGTAAATCCTCATCA | AA | TGTTGTACATGGAGTCCTGGGGCCC | GGTAGGAGTCGCCGCCGTTCATGCT | AA | ATCATCCAGTAAACCGCC |
| 4 | CCTCGTAAATCCTCATCA | AA | GACCCTGCGTCGTGGGCACCATGCT | GGGGCGGACTGATCATCTGACCCTG | AA | ATCATCCAGTAAACCGCC |
| 5 | CCTCGTAAATCCTCATCA | AA | GCTGCGTACAGGTTTGCCTCTTCTT | TTTTTCTTTTTCGGGAAGACTTTCT | AA | ATCATCCAGTAAACCGCC |
| 6 | CCTCGTAAATCCTCATCA | AA | TCCGCTTGTTGCCGAACCAATTAGA | CTTTCCCTATGTTCTTCTTGTATCG | AA | ATCATCCAGTAAACCGCC |
| 7 | CCTCGTAAATCCTCATCA | AA | GGCGAGTTCCTCCTTGGCCTCCTCC | CTGAGAGACGGTGATGCCGCACTTC | AA | ATCATCCAGTAAACCGCC |
| 8 | CCTCGTAAATCCTCATCA | AA | AAATACTCGTTCAGGATCTCCGTCG | GGGTACGGGTTGCTAAGGTGCGAGT | AA | ATCATCCAGTAAACCGCC |
| 9 | CCTCGTAAATCCTCATCA | AA | TTGCGTCGAGGAACCTGGACCTGAG | GCTTGCTGAAGTTGCGTCTCTTTCG | AA | ATCATCCAGTAAACCGCC |
| 10 | CCTCGTAAATCCTCATCA | AA | GAGCTGCACCTGGATGGAGTTGAAC | CATGACGGCTTCGCAGGTGCTCTGC | AA | ATCATCCAGTAAACCGCC |
| 11 | CCTCGTAAATCCTCATCA | AA | ATCTCCTTGGGCGTGATCGGCCGCG | TTGTGGATTATCTGCACCATGCGCT | AA | ATCATCCAGTAAACCGCC |
| 12 | CCTCGTAAATCCTCATCA | AA | CGTGCGTCGTGAACTCGTTGCACGC | GACTCTGCTCTCGCAAAAGGTTCAT | AA | ATCATCCAGTAAACCGCC |
| 13 | CCTCGTAAATCCTCATCA | AA | GATCTGCCTGATCTGGGCGAGCTTG | CTCGTACTTCTCCAGCTCCTGGTGG | AA | ATCATCCAGTAAACCGCC |
| 14 | CCTCGTAAATCCTCATCA | AA | CCGCTGCCCCGGCGCCGCCGCCCTT | CCCCGGAGGCCGCCGCAGACGCGTT | AA | ATCATCCAGTAAACCGCC |

**Table S16.** Probe pairs designed for *Ixodes scapularis dac-A* HCR in situ hybridization (B3 initiator).

| Pair | Initiator | Spacer | Hybridization | Hybridization | Spacer | Initiator |
|---|---|---|---|---|---|---|
| 1 | GTCCCTGCCTCTATATCT | TT | ACCGTTTTGTTGCAACGAGATCCTC | GCGGTCCGTACCTGTCGATCGAAAG | TT | CCACTCAACTTTAACCCG |
| 2 | GTCCCTGCCTCTATATCT | TT | AGTGTATTGCACCCGGAATCGAACG | AAGCGGATGGTTTTGCGGCTCGAGT | TT | CCACTCAACTTTAACCCG |
| 3 | GTCCCTGCCTCTATATCT | TT | GGCGTCTGGGTAATGCTGCGAAAGG | CGTCATCTATTGTGGTTTGAGGAAT | TT | CCACTCAACTTTAACCCG |
| 4 | GTCCCTGCCTCTATATCT | TT | CCCTGAACCAAATCGGTCGATGTCT | GCTGAAAGTGACGGCGCTGCGTTGC | TT | CCACTCAACTTTAACCCG |
| 5 | GTCCCTGCCTCTATATCT | TT | GCTTTCACCCCGTTCATACCAATTA | TTTCCACGAACTGCTTTTGAAAGCA | TT | CCACTCAACTTTAACCCG |
| 6 | GTCCCTGCCTCTATATCT | TT | GAATCGTAAAGCGCTCACTGTGTTT | TACGTAGGGAAGAACAGCCTTTGGC | TT | CCACTCAACTTTAACCCG |
| 7 | GTCCCTGCCTCTATATCT | TT | ATATTGCTTCTTGCAGTGGTGCAGT | AAAAACGAAACAGATGGAGGAACGG | TT | CCACTCAACTTTAACCCG |
| 8 | GTCCCTGCCTCTATATCT | TT | AGAGCAGCTTGCAGCGATTCACCCC | TGTAGAGCGCGTCAAAGTCCTTGCA | TT | CCACTCAACTTTAACCCG |

| | | | | | | |
|---|---|---|---|---|---|---|
| 9 | GTCCCTGCCTCTATATCT | TT | TTACACACGATCGGACTGATGCCCA | CCCCGCAGCACGCGGACTTGCTCGA | TT | CCACTCAACTTTAACCCG |
| 10 | GTCCCTGCCTCTATATCT | TT | AACCGCCCACCAGGTGCCTGAGGAA | GCTTGAGCTTTGTGTACACCGTGTG | TT | CCACTCAACTTTAACCCG |
| 11 | GTCCCTGCCTCTATATCT | TT | GTACTCGCCGCGGTACGCGAACGCC | CTCGAAGGCCTGGGGCAGGCAGAGC | TT | CCACTCAACTTTAACCCG |
| 12 | GTCCCTGCCTCTATATCT | TT | CTGCAGTCCGTCGCGTCGGCGGCCA | ATCTTCTCGCCGCGGTACTGGATGA | TT | CCACTCAACTTTAACCCG |
| 13 | GTCCCTGCCTCTATATCT | TT | CGTTGGACAAGGGGTCGAGACCCCG | ACCTCGACGAGGGACTCTTGAGCCT | TT | CCACTCAACTTTAACCCG |
| 14 | GTCCCTGCCTCTATATCT | TT | GCAGAGGGCGGCCGCGTTGAAAGCC | CCCGTTGATGACCGAGCAGAACTGG | TT | CCACTCAACTTTAACCCG |
| 15 | GTCCCTGCCTCTATATCT | TT | GACGGGGACGAGGAACCGCGGCTGT | GGGAGCGGTGGCGGCAGGTCGTGGG | TT | CCACTCAACTTTAACCCG |

**Table S17.** Probe pairs designed for *Ixodes scapularis dac-B* HCR in situ hybridization (B3 initiator).

| Pair | Initiator | Spacer | Hybridization | Hybridization | Spacer | Initiator |
|---|---|---|---|---|---|---|
| 1 | GTCCCTGCCTCTATATCT | TT | CGATCTGAATCAACTTCCGGTGACG | TTTTTGTCGGGTTCCTGTCTGCTTT | TT | CCACTCAACTTTAACCCG |
| 2 | GTCCCTGCCTCTATATCT | TT | TTGCGGTTGCTGCTGCTGCTGCGAT | GGACTCTGGACGTTGTAGAGGCTGC | TT | CCACTCAACTTTAACCCG |
| 3 | GTCCCTGCCTCTATATCT | TT | CCTCTTCGTACTGAGACCGTTTCTT | GCGGGAGTGTGGCCGTGGTTCGCAG | TT | CCACTCAACTTTAACCCG |
| 4 | GTCCCTGCCTCTATATCT | TT | CGCCGACAGCGGCGTTCTTTCTTGA | TCGAGTTCCAGCTGGTCCTGGATAC | TT | CCACTCAACTTTAACCCG |
| 5 | GTCCCTGCCTCTATATCT | TT | GCTCCTCCGTTAGCTGCTTTTCCAG | TCTTTTGGTAGAGCACCCTGGTCCG | TT | CCACTCAACTTTAACCCG |
| 6 | GTCCCTGCCTCTATATCT | TT | CTCCATCTTGAGCTCGGCCTTTTCG | TTCCCGAAGTCCTCGTTCTCGTAAG | TT | CCACTCAACTTTAACCCG |
| 7 | GTCCCTGCCTCTATATCT | TT | CACTTTGAGGAGTCCCTGGATGTTT | GTCTTGGTGCCTGGCGTTGTCGGCG | TT | CCACTCAACTTTAACCCG |
| 8 | GTCCCTGCCTCTATATCT | TT | ATTTCTTGGTCTCTGTCGTCGGTGT | GGGCTGCTGCTTTCTTCGCCGCCGC | TT | CCACTCAACTTTAACCCG |
| 9 | GTCCCTGCCTCTATATCT | TT | TCCGTGCGGTGAGCGTGTTGCGACA | GCGCCATCTTGGTATTCCCTAGGCT | TT | CCACTCAACTTTAACCCG |
| 10 | GTCCCTGCCTCTATATCT | TT | ATGCCGACGAAGACAGACCACCATT | TCAGCACCGGAGGACCATGTCCATT | TT | CCACTCAACTTTAACCCG |
| 11 | GTCCCTGCCTCTATATCT | TT | TCCATCGTAGGACGAGAACATCCGG | GAAGGCAGCCGAGTCTTTGAGTCGG | TT | CCACTCAACTTTAACCCG |
| 12 | GTCCCTGCCTCTATATCT | TT | CCGTCTCGCTCTTGTTGTGATCAAA | CGCTGCCGTGGGGCTTGTCCGCATG | TT | CCACTCAACTTTAACCCG |
| 13 | GTCCCTGCCTCTATATCT | TT | TGCACCACCATCAACGTGGTCGAGT | CACCTCAGGTACCGATACCGACACG | TT | CCACTCAACTTTAACCCG |
| 14 | GTCCCTGCCTCTATATCT | TT | CCTTTTCGATGCTGATCTGTCGGTC | GTGGCGATGATTCCAAAGTCCACTG | TT | CCACTCAACTTTAACCCG |
| 15 | GTCCCTGCCTCTATATCT | TT | AGGAGTCCCTGGATGTTTCGAAGCA | TGCCTGGCGTTGTCGGCGGCCACTT | TT | CCACTCAACTTTAACCCG |
| 16 | GTCCCTGCCTCTATATCT | TT | CACCGCCGCTCATTTCTTGGTCTCT | CATGGGGTCCCGGGCTGCTGCTTTC | TT | CCACTCAACTTTAACCCG |
| 17 | GTCCCTGCCTCTATATCT | TT | GTGCGGTGAGCGTGTTGCGACAGGT | CCATCTTGGTATTCCCTAGGCTCGT | TT | CCACTCAACTTTAACCCG |
| 18 | GTCCCTGCCTCTATATCT | TT | CCGACGAAGACAGACCACCATTGAT | GGACCGGAGGACCATGTCCATTGGA | TT | CCACTCAACTTTAACCCG |
| 19 | GTCCCTGCCTCTATATCT | TT | ATCGTAGGACGAGAACATCCGGTCC | GGCAGCCGAGTCTTTGAGTCGGCTT | TT | CCACTCAACTTTAACCCG |
| 20 | GTCCCTGCCTCTATATCT | TT | TCGATGCGCCCCTTCTTGCTCATCA | TGACCGTTCTCGTAGCCGGGGTAGT | TT | CCACTCAACTTTAACCCG |
| 21 | GTCCCTGCCTCTATATCT | TT | TGGAATGTGTTTCCCGTTTGGGGAA | GGGGACGGGCTTGGTTCGCAACTCG | TT | CCACTCAACTTTAACCCG |
| 22 | GTCCCTGCCTCTATATCT | TT | TCGATGCGCCCCTTCTTGCTCATCA | TGACCGTTCTCGTAGCCGGGGTAGT | TT | CCACTCAACTTTAACCCG |
| 23 | GTCCCTGCCTCTATATCT | TT | GGTAGACCCTGGTAGCTTCTCCGAA | GACGGGCTTGGCGGGTCTGGAAACT | TT | CCACTCAACTTTAACCCG |
| 24 | GTCCCTGCCTCTATATCT | TT | TGCAGTCCTTGTAGAGCGCATCAAA | ACAGGCGGCTGGCAGATCTCGCCGT | TT | CCACTCAACTTTAACCCG |
| 25 | GTCCCTGCCTCTATATCT | TT | ACGATCGGCGTGATGTCCAGCCTCT | AGGATCCTGACCTGCTCGACGTTGC | TT | CCACTCAACTTTAACCCG |
| 26 | GTCCCTGCCTCTATATCT | TT | ACCAGGTGCTTGAGGAACAGCTCGA | TTCGTGTACACCGTGTGCAGCCCTC | TT | CCACTCAACTTTAACCCG |

**Table S18.** Probe pairs designed for *Pycnogonum litorale exd* HCR in situ hybridization (B1 initiator).

| Pair | Initiator | Spacer | Hybridization | Hybridization | Spacer | Initiator |
|---|---|---|---|---|---|---|
| 1 | GAGGAGGGCAGCAAACGG | AA | GCCTAAAGCTGCCTAATTTAACAAA | AAAAATAAGCTCGCATCGTTTTTAA | TA | GAAGAGTCTTCCTTTACG |
| 2 | GAGGAGGGCAGCAAACGG | AA | TTAAAACAATGTTGCCATTAATTCA | TTTTTAAAAGTGGCGAGGTTTCATC | TA | GAAGAGTCTTCCTTTACG |
| 3 | GAGGAGGGCAGCAAACGG | AA | ACAGTCAGCCAAATAATAAATATTT | TTTGTAAAATTATATTAAATATTAT | TA | GAAGAGTCTTCCTTTACG |
| 4 | GAGGAGGGCAGCAAACGG | AA | ATTATTTATTTAAAATTTTAATAAC | AAAAATCTTTACGTTTATGATACCT | TA | GAAGAGTCTTCCTTTACG |
| 5 | GAGGAGGGCAGCAAACGG | AA | ATCTTTTAAATTGTATTTTAACACA | AAAAATTTTTCAAAACAGAATCCAA | TA | GAAGAGTCTTCCTTTACG |
| 6 | GAGGAGGGCAGCAAACGG | AA | TAAATATATATACTTGGTTTGTTAC | AAAAATAAAGGTATCATAACGTCAT | TA | GAAGAGTCTTCCTTTACG |
| 7 | GAGGAGGGCAGCAAACGG | AA | TCACTAGGACTGGCTGCAGATTGTG | TGTGGTGGGACGCTATTGGCTGACA | TA | GAAGAGTCTTCCTTTACG |
| 8 | GAGGAGGGCAGCAAACGG | AA | GGTGCATATCGTACATCCCACCTGA | TCGACGAATGATTATATCCTCCAAC | TA | GAAGAGTCTTCCTTTACG |
| 9 | GAGGAGGGCAGCAAACGG | AA | TCCAGTCTGTGATTGTACGTTGGCA | ATGTGCTGGTAATCCGTCTGAGTAT | TA | GAAGAGTCTTCCTTTACG |
| 10 | GAGGAGGGCAGCAAACGG | AA | CCTCCATTCATACTCATACTTCCAG | ATTGAAGAAATTTGTTGGTAAGAAT | TA | GAAGAGTCTTCCTTTACG |
| 11 | GAGGAGGGCAGCAAACGG | AA | TCATTTGACCTTGACCCTGTGATGA | ACATCTGGTCACCCGGAGGAGGACT | TA | GAAGAGTCTTCCTTTACG |
| 12 | GAGGAGGGCAGCAAACGG | AA | GACCTCAGTATCATGACGGCTTCGC | TTTTTCCTCCTAGCATCTAAGAACC | TA | GAAGAGTCTTCCTTTACG |
| 13 | GAGGAGGGCAGCAAACGG | AA | TCTATTTCTTTCGGCGTAATTGGCC | TTTTTGTGAATAATTTGGACCATTC | TA | GAAGAGTCTTCCTTTACG |
| 14 | GAGGAGGGCAGCAAACGG | AA | TCACGTGCGTCGTAAACTCGTTACA | TTCTACTTTGTTCTCTCAGAAGGTT | TA | GAAGAGTCTTCCTTTACG |
| 15 | GAGGAGGGCAGCAAACGG | AA | GTATATTTGTCGAATTTGAGCTAAT | CTGTTCGTATTTTTCCAATTCCTGA | TA | GAAGAGTCTTCCTTTACG |
| 16 | GAGGAGGGCAGCAAACGG | AA | CTGTCCGGTTGTCCAGGACCACCTG | GCGCGATAATCGGAATGCTCGATAG | TA | GAAGAGTCTTCCTTTACG |
| 17 | GAGGAGGGCAGCAAACGG | AA | CAGCACCTCCGCCTTTTTCTGGACC | CTGCAGCCGATGCATTAGCCGCCGC | TA | GAAGAGTCTTCCTTTACG |
| 18 | GAGGAGGGCAGCAAACGG | AA | CAACCGCATTAGTTGTGGATCGGGC | GACACCTTCCGCTATTAACATGTTA | TA | GAAGAGTCTTCCTTTACG |
| 19 | GAGGAGGGCAGCAAACGG | AA | GAACAATGCCGGTTTCATACGATGA | TTTTTCTTTAATTTCGCACAATACA | TA | GAAGAGTCTTCCTTTACG |
| 20 | GAGGAGGGCAGCAAACGG | AA | TCATCTAAGCTTTGATCGGTAATGT | TTGAGTGTATGTTTTCTCGCTTGCG | TA | GAAGAGTCTTCCTTTACG |
| 21 | GAGGAGGGCAGCAAACGG | AA | CTTGTTTACGGCCATCCTGATCGTG | TAATTTGTTGTAATATTTCACCAAT | TA | GAAGAGTCTTCCTTTACG |
| 22 | GAGGAGGGCAGCAAACGG | AA | TGATGGGGCTTGAAGACCGTATCCA | CGGTGGTTGATGGCCTGGATCACCG | TA | GAAGAGTCTTCCTTTACG |
| 23 | GAGGAGGGCAGCAAACGG | AA | ACAACCGCATGGCCGTGCTGCTGCA | TGAGGTACGACATGACCAGCCATAC | TA | GAAGAGTCTTCCTTTACG |
| 24 | GAGGAGGGCAGCAAACGG | AA | TTACTCGACTAATGTATAAATCTAG | TTCTTTGTTGCTCGTCCATATCCTA | TA | GAAGAGTCTTCCTTTACG |
| 25 | GAGGAGGGCAGCAAACGG | AA | AAAACTATTCAATACACTAACAGCG | AAAAACTAGCGAAGGAGAGAACGAC | TA | GAAGAGTCTTCCTTTACG |
| 26 | GAGGAGGGCAGCAAACGG | AA | CTTCAATCCACGACACACTCACACT | ACTTTATCGTAGTAGTCACTGCTTC | TA | GAAGAGTCTTCCTTTACG |

**Table S19.** Probe pairs designed for *Pycnogonum litorale dac* HCR in situ hybridization (B5 initiator).

| Pair | Initiator | Spacer | Hybridization | Hybridization | Spacer | Initiator |
|---|---|---|---|---|---|---|
| 1 | CTCACTCCCAATCTCTAT | AA | CCGTAATATTTACAATTTATATTTA | GTTCAACAATAATACTACTTTATAA | AA | CTACCCTACAAATCCAAT |
| 2 | CTCACTCCCAATCTCTAT | AA | TGAAATTATGAAACGACCCCTACGT | ATGAAGAATCTTTTTCGCGTTTTAA | AA | CTACCCTACAAATCCAAT |
| 3 | CTCACTCCCAATCTCTAT | AA | GCCAACGATTCGTTAAGTATTCTTA | TTGTTTCTGTCCCGTTCTATGTCGT | AA | CTACCCTACAAATCCAAT |
| 4 | CTCACTCCCAATCTCTAT | AA | CTGATCCTTTGTTCTTCCAACAATT | TTTTTGATCCGTTTTTGGTACAAAA | AA | CTACCCTACAAATCCAAT |
| 5 | CTCACTCCCAATCTCTAT | AA | TTGCCTGGCATTGTCGGCTGCCACT | TTTCTCGTAGTTGCTTTGTCGTTCT | AA | CTACCCTACAAATCCAAT |

| | | | | | | |
|---|---|---|---|---|---|---|
| 6 | CTCACTCCCAATCTCTAT | AA | TGTTGTTCAACATTCCGACAGCTTG | CGGCCGTCATGGTGCCCATGCCAGG | AA | CTACCCTACAAATCCAAT |
| 7 | CTCACTCCCAATCTCTAT | AA | TCTTCTTCGGTGTCGTTTTCGTCTT | AAATCGTGTTCTCTGTCGTCGTCGT | AA | CTACCCTACAAATCCAAT |
| 8 | CTCACTCCCAATCTCTAT | AA | CTTCGCTACCACTGTGTTCTGTTCC | CGTCTTCTCCGACGTCATTGAAAGC | AA | CTACCCTACAAATCCAAT |
| 9 | CTCACTCCCAATCTCTAT | AA | TTGTGACAGATTTAATACTGAGCTG | ATTTCCGCCGCCATGGTTACGGTTG | AA | CTACCCTACAAATCCAAT |
| 10 | CTCACTCCCAATCTCTAT | AA | GAATCTTTGATCCTGTGGTGCTCGT | CCATTCGTCGAACCGTTCGCGAATG | AA | CTACCCTACAAATCCAAT |
| 11 | CTCACTCCCAATCTCTAT | AA | TGTCGCTCGATTCGCATCGTTCTTC | ATGCAGGCGAACTGTAAAGTCGCTC | AA | CTACCCTACAAATCCAAT |
| 12 | CTCACTCCCAATCTCTAT | AA | GGCGCTACTGATGGCCGAACGGTGG | TGATCTGGATGAATTGATGGCGTCG | AA | CTACCCTACAAATCCAAT |
| 13 | CTCACTCCCAATCTCTAT | AA | CTGGTCGTTGGTAGGAGTCCTGACA | TTGTTGGAAGTGACGTCATTGGATC | AA | CTACCCTACAAATCCAAT |
| 14 | CTCACTCCCAATCTCTAT | AA | GACTAGAATTTGATAATGCGCTCGG | GGTGGTGGTGATGATGATGATTTGG | AA | CTACCCTACAAATCCAAT |
| 15 | CTCACTCCCAATCTCTAT | AA | TCCACCTTGATGATGGTGGTGGTGA | TAACAGACTCGATGCTCCACCACCA | AA | CTACCCTACAAATCCAAT |
| 16 | CTCACTCCCAATCTCTAT | AA | ATCGGTATGAATGGTAAAGAATTGA | CCTCCGTGGACACCATGATGACTTG | AA | CTACCCTACAAATCCAAT |
| 17 | CTCACTCCCAATCTCTAT | AA | CTGCATGGTGACTGTAACCATTGGC | GTGCAGCTGCAGCTGCTACCTGGGC | AA | CTACCCTACAAATCCAAT |
| 18 | CTCACTCCCAATCTCTAT | AA | TGCACGAAGAAATTCGGCTGTGATG | CAATGGATTCTTCTCCAATTGTTCG | AA | CTACCCTACAAATCCAAT |
| 19 | CTCACTCCCAATCTCTAT | AA | TTATCAATGCGACATTTCTTGGCCT | CCGTTTTCGTATCCCAGGTAATCTC | AA | CTACCCTACAAATCCAAT |
| 20 | CTCACTCCCAATCTCTAT | AA | GGCTCATACCGTCAATTAATGAGGC | TTATTGCGCTATCAGATGGATGTGT | AA | CTACCCTACAAATCCAAT |
| 21 | CTCACTCCCAATCTCTAT | AA | CCGATTGAGCCAACGCCACCGCCAC | TTTTTAGCACTAGGGTCACAATGAC | AA | CTACCCTACAAATCCAAT |
| 22 | CTCACTCCCAATCTCTAT | AA | AAACCGATGGCCGATGATGGTGATG | CACCTGTAGATTGCGGTGGACTGTT | AA | CTACCCTACAAATCCAAT |
| 23 | CTCACTCCCAATCTCTAT | AA | CATACTCGGAACCATGCTCGATATA | ATGGTGATGCTGTTCTAATGCTTTG | AA | CTACCCTACAAATCCAAT |
| 24 | CTCACTCCCAATCTCTAT | AA | TTCGAAGCCACAGACGCTGCTAAAG | GAAGCACTGAGTGCTCCGTACGGGA | AA | CTACCCTACAAATCCAAT |
| 25 | CTCACTCCCAATCTCTAT | AA | CGTTATCGCCGTGAGACGAAGTGAC | GCGGTAACATCGGCGGCGATGGACT | AA | CTACCCTACAAATCCAAT |
| 26 | CTCACTCCCAATCTCTAT | AA | TGACATTGGACTCGTACTATTCTCG | AGTCGTGAGGCCAGTGGCAGGACCA | AA | CTACCCTACAAATCCAAT |
| 27 | CTCACTCCCAATCTCTAT | AA | TCCAGTGCTCCTATCGATGGAGGCT | GGCGACGGTTGCTTCATGTTCAGCA | AA | CTACCCTACAAATCCAAT |
| 28 | CTCACTCCCAATCTCTAT | AA | TCAGCTGACCATTGGTACTCGGAAG | GCGGCGATCCGAGAGGCGTCGACGG | AA | CTACCCTACAAATCCAAT |
| 29 | CTCACTCCCAATCTCTAT | AA | CGACGACGATGCAGCATCAGTTTGA | CGGCGGCGGCGACGGCGACGGCGAA | AA | CTACCCTACAAATCCAAT |
| 30 | CTCACTCCCAATCTCTAT | AA | ATGATGAATGAATGAATGAATAATG | CCAGAGCTAACAGGCTGCTGCTGCT | AA | CTACCCTACAAATCCAAT |

**Table S20.** Primer pairs used in RNAi experiments. Primer pairs for *Po-dac* were previously published in Sharma et al. (2013).

| Primer ID | Forward Primer | Reverse Primer |
|---|---|---|
| *Po-exd* | 5'-GCGCAAGCTAGGAAACACAC-3' | 5'-ATTCCTCCTTGGCTTCTTCG-3' |
| *Po-N* | 5'-CGGTTTCAAAGGTGTCGATT-3' | 5'-TGACGCAAGGATTGCTGTAG-3' |
